# Structure and Mechanism of a Two-component Lanthipeptide Toxin

**DOI:** 10.64898/2026.05.13.724966

**Authors:** Ryan Moreira, Constantin Giurgiu, Imran R. Rahman, MacKenzie Patterson, Stefan T. Huber, Yi Yang, Kevin Jeanne Dit Fouque, Francisco Fernandez-Lima, Ge Liu, Ruihan Guo, Payam Kelich, Po-Chao Wen, Emad Tajkhorshid, Alex G. Johnson, Wilfred A. van der Donk

## Abstract

The enterococcal cytolysin is a two-component lanthipeptide natural product that co-operatively kills mammalian cells and bacteria. It is a virulence factor in typically commensal strains of gut-dwelling *Enterococcus faecalis* and causes a fatal form of alcoholic hepatitis. Despite its clinical significance, little is known about the mechanism of action. We present here the molecular details of cytolysin’s remarkable bioactivity. A high-resolution cryo-electron microscopy structure revealed highly ordered tubular assemblies comprising the two subunits. We further demonstrate these structures disrupt the membranes of eukaryotic and bacterial cells. These results provide the first high-resolution structure of two distinct lanthipeptides interacting with one another and offer an explanation for the unique bioactivity of the cytolysin toxin.

## Main Text

The enterococcal cytolysin is a two-component ribosomally translated and post-translationally modified peptide (RiPP) natural product that is a virulence factor in *E. faecalis* and is responsible for the cytotoxic activity of virulent strains (*1*). A member of the human gut microbiome, *E. faecalis* is a widespread commensal organism and a substantial fraction of the human population is believed to host cytolysin-producing (C+) *E. faecalis* (*2*). C+ *E. faecalis* is not a danger to its host when it is in equilibrium with other members of the gut community; however, certain disease states are exacerbated by commensal C+ *E. faecalis* (*3*). Recently, cytolysin was shown to produce a severe form of alcoholic hepatitis that is often fatal (*4*). To date, no treatment for cytolysin-induced disease states has been reported.

Cytolysin is a member of the lanthipeptide class of RiPPs and both components contain rigidifying thioether bridges called lanthionines or methyllanthionines. Lanthionines are generated from the dehydration of Ser to dehydroalanine (Dha) followed by the Michael-type addition of a cognate Cys (*5*). Methyllanthionines are biosynthesized by an analogous pathway that starts with dehydration of Thr to dehydrobutyrine (Dhb). Cytolysin is composed of a long subunit CylL_L_″ (38 residues, 3.4 kDa) and a short subunit CylL_S_″ (21 residues, 2 kDa) that possess non-overlapping ring patterns (Fig. 1A) (*6*). Solution NMR studies revealed that CylL_L_″ possessed an α-helical conformation conferred by its lanthionine rings akin to a natural form of peptide stapling (*7, 8*). Cytolysin displays a unique bioactivity profile as it is capable of damaging mammalian and bacterial cells (*9, 10*). Previous studies demonstrated that CylL_S_″ and CylL_L_″ act in a 1:1 ratio at nanomolar concentrations to kill gram-positive bacteria and cause hemolysis (*11*). The bioactivity of cytolysin is unusually potent. An analysis of the activities of 2,644 antimicrobial peptides (AMPs) revealed that only six had antimicrobial activity similar to cytolysin, with none of these displaying nanomolar activity against erythrocytes (*12, 13*). Despite its clinical significance, little is known about the mode of action (MoA) of cytolysin.

**Fig. 1.**
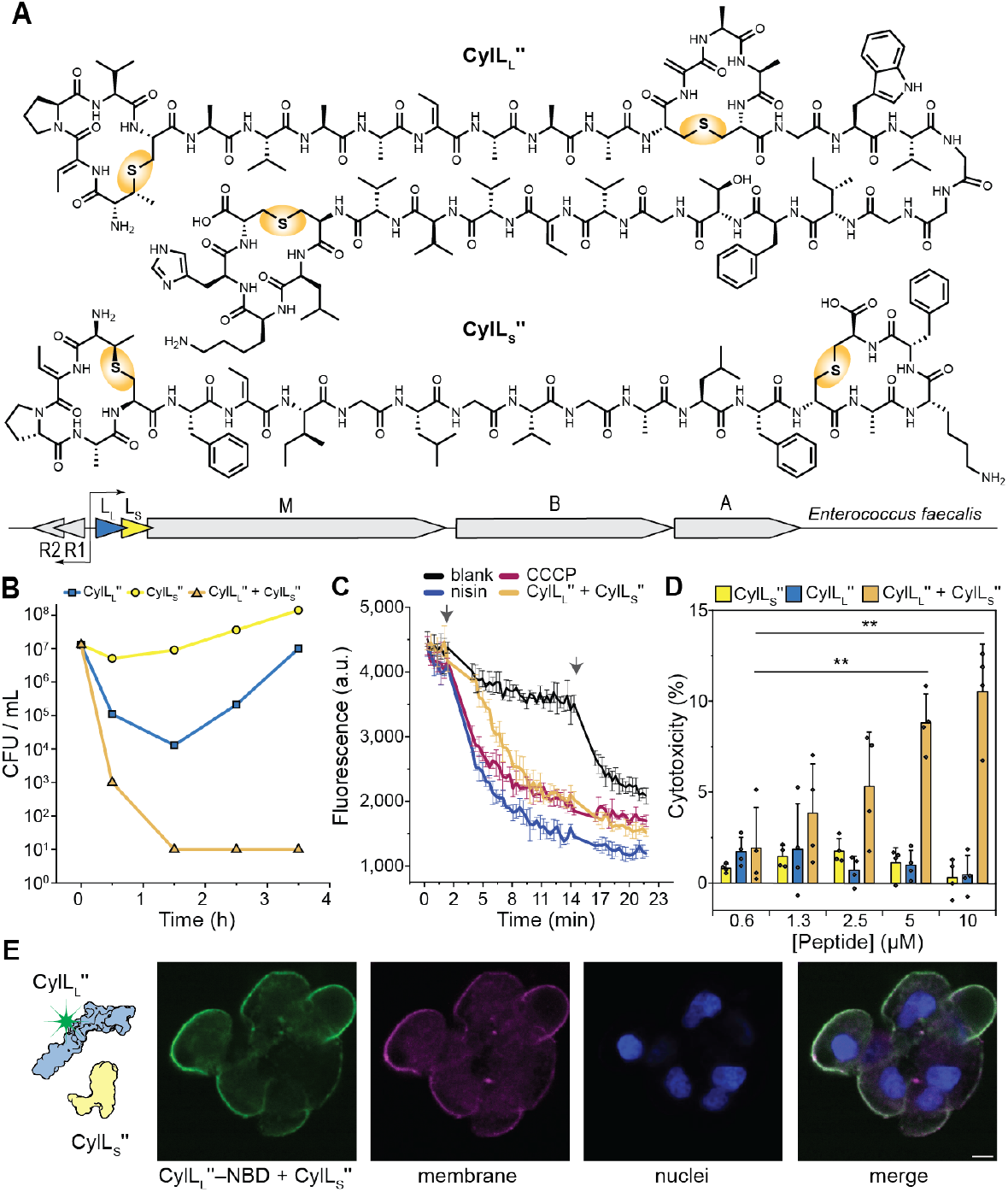
CylL_L_″ and CylL_S_″ co-operatively disrupt the membranes of bacteria and liver cells. A) The chemical structures of CylL_L_″ and CylL_S_″. The thioether crosslinks are highlighted. B) Change in the number of viable colonies of *L. lactis subsp. cremoris* over time after treatment with CylL_L_″, CylL_S_″ or CylL_L_″ and CylL_S_″ mixed in a 1:1 ratio. The total peptide concentration was 32 nM. C) pH-sensitive fluorescent dyes were used to monitor changes in the intracellular pH of *L. lactis subsp. cremoris* over time. Cells were treated with both cytolysin peptides (2 × MIC), nisin (2 × MIC), or CCCP. Chronologically, the first arrow indicates when peptide or CCCP was added and the second arrow indicates when CCCP was added (for the black, blue and yellow lines). Data from three experiments are presented (*n* = 3). D) LDH activity from the media of HepG2 cells treated with CylL_L_″:CylL_S_″ (1:1), CylL_S_″ or CylL_L_″ 3 h prior to activity measurements. Data from four trials are presented (*n* = 4). Total peptide concentrations are listed on the horizontal axis. E) Confocal microscopy of HepG2 cells treated with a 1:1 mixture of CylL_L_″-nitrobenzoxadiazole (NBD) and CylL_S_″. The total peptide concentration was equal to 10 μM. H33342 = Hoechst33342 for nuclear staining. The membrane was stained with CellBrite® Orange.

Considering the clinical impact of cytolysin, along with its unique structure and bioactivity, elucidation of its MoA is valuable to guide potential intervention strategies. Here, we show that CylL_S_″ and CylL_L_″ co-operatively form assemblies in solution that disperse into cellular membranes causing rapid membrane destabilization and depolarization. A high-resolution structure of these assemblies shows that the two subunits evolved to form an unusual helical supramolecular arrangement of high stability. This structure provides an unparalleled glimpse into the interaction between two distinct lanthipeptides and offers a compelling explanation for the cross-kingdom toxicity of cytolysin.

The high-content of hydrophobic residues in CylL_S_″ and CylL_L_″ suggested a membrane-target. To probe this hypothesis, CylL_S_″ and CylL_L_″ were prepared in *E. coli* using the biosynthetic enzymes (*6*) and their antibiotic activity was evaluated against *Lactococcus lactis subsp. cremoris*, a cytolysin-susceptible strain (minimum inhibitory concentration (MIC) = 8 nM). The individual subunits of cytolysin possessed modest bactericidal activity against *L. lactis subsp. cremoris* (Fig. 1B). In contrast, a stoichiometric mixture of CylL_S_″:CylL_L_″ (32 nM) reduced the number of viable bacteria by four orders of magnitude within 30 minutes (Fig 1B). Peptide antibiotics that target the membrane are often bactericidal by dissipating the proton motive force (*14*). The effect of cytolysin on the membrane integrity of *L. lactis subsp. cremoris* was monitored using the DNA intercalating dye SYTOX green (*15*). Cytolysin permeabilized the membranes of *L. lactis subsp. cremoris* within minutes at two times the MIC (Fig. S1) and half maximal activity was observed ten minutes after treatment. We used carboxyfluorescein diacetate (5(6)-CFDA) to assess the impact of cytolysin on the proton motive force (*16*). A stoichiometric mixture of CylL_S_″ and CylL_L_″ dissipated the pH gradient across the bacterial membrane to the same extent as the protonophore carbonyl cyanide *m*-chlorophenyl hydrazone (CCCP) within ten minutes of treatment at two times the MIC (Fig. 1C). Compared to the known, potent, pore forming lanthipeptide nisin (*17*), the action of cytolysin proceeded after a short lag time.

Genomic and transcriptional analysis of resistant mutants of *L. lactis* supported a membrane-targeted mechanism. Genomic analysis revealed few changes at the genetic level and none that were common (Fig. S1). The transcriptome of a spontaneous resistant strain of *L. lactis* was analyzed using RNA-Seq and suggesting a passive resistance mechanism that involved the upregulation of genes previously associated with resistance to cationic antimicrobial peptides and detergents (Fig. S1) (*18, 19*). Peptides that target the cellular membrane typically permeabilize model membranes at peptide:lipid (P:L) ratios of 1:50 to 1:500 (*20*). Cytolysin permeabilized liposomes composed of 1,2-dioleoyl-sn-glycero-3-phosphocholine (DOPC) at a P:L ratios of 1:123 to 1:1111 (Fig. S1). At a P:L ratio of 1:250, cytolysin fully permeabilized liposomes with compositions that modelled bacteria and epithelial cells (Fig. S1) consistent with cytolysin’s potent cross-kingdom activity.

To determine whether a similar mechanism could explain cytolysin’s liver toxicity (*4*), the hepatocellular carcinoma epithelial cell line HepG2 was used as a model (*21*). Acute leakage of cellular contents was monitored through assessment of lactate dehydrogenase (LDH) activity in the growth media after treatment with CylL_S_″, CylL_L_″ or a stoichiometric mixture of CylL_S_″:CylL_L_″ (Fig. 1D). Only the combination of both subunits caused a dose-dependent LDH leakage consistent with findings using primary hepatocytes (*4*). A fluorescent form of cytolysin was prepared by labelling CylL_L_″ with a fluorophore. A stoichiometric combination of labelled CylL_L_″ and unlabelled CylL_S_″ possessed similar bioactivity to wild type (WT) (Figs. S2) and localized to the cell membrane (Fig. 1E).

Some membrane active peptides assemble in solution to form amyloid fibrils. The most acutely toxic form of these peptides are oligomers that assemble concomitantly with larger aggregates (*22-25*). CylL_S_″ and CylL_L_″ co-operatively form aggregates in solution in a target-independent fashion (*26*) suggesting that a heterooligomeric form of cytolysin could be the bioactive form. High-resolution mass spectrometry analysis of a mixture CylL_S_″ and CylL_L_″ showed the formation of heterooligomeric species consistent with assembly occurring through an alternating copolymerization (Fig. S3). Negative staining transmission electron microscopy (TEM) analysis revealed that the individual subunits of cytolysin, which are not bioactive, assembled into spherical aggregates in solution (Fig. S4). In contrast, a 1:1 mixture of CylL_S_″:CylL_L_″ formed well-defined nanotubes with water filled channels that were approximately 5–6 nm in diameter and a length of >10 nm, which co-existed with toroidal ring-shaped features (length <10 nm) (Fig. 2A). Analysis of 10,913 features showed that rings outnumbered tubes 8,986 to 1,927. The tube length was normally distributed, with an average of 104 ± 39 nm (Fig. S4). SDS-PAGE analysis showed a well-defined band at approximately 130 kDa that greatly exceeded the sum of the masses of the two peptides and suggested that an assembled form of cytolysin was stable in environments that mimic a membrane (*26*) (Fig. S4). The nanotubes and rings were stable to extraction from liposomes with mild detergents (Fig. S4). CylL_S_′′ and CylL_L_′′ are made by proteolytic processing of CylL_S_′ and CylL_L_′, which are not bioactive (*27*) (Figs. S5 and S6). Consistent with bioactivity being imparted by the assemblies, CylL_S_′ and CylL_L_′ did not form nanotubes or rings (Fig. S6). Importantly, SDS-PAGE analysis of several cytolysin analogs prepared by site-directed mutagenesis of the biosynthetic precursors showed a clear correlation between the ability to form the 130 kDa assemblies and their bioactivity (Fig. S6) (*28*). To investigate whether the nanotubes are involved in antimicrobial activity, *L. lactis subsp. cremoris* was treated with both subunits of cytolysin, and then negatively stained and analyzed by TEM. Features that strongly resembled the nanotubes and rings were evenly distributed on the surface of the cells (Fig. 2B). These nanotubes were in different orientations relative to the surface of the cell with some parallel and others perpendicular.

**Fig. 2.**
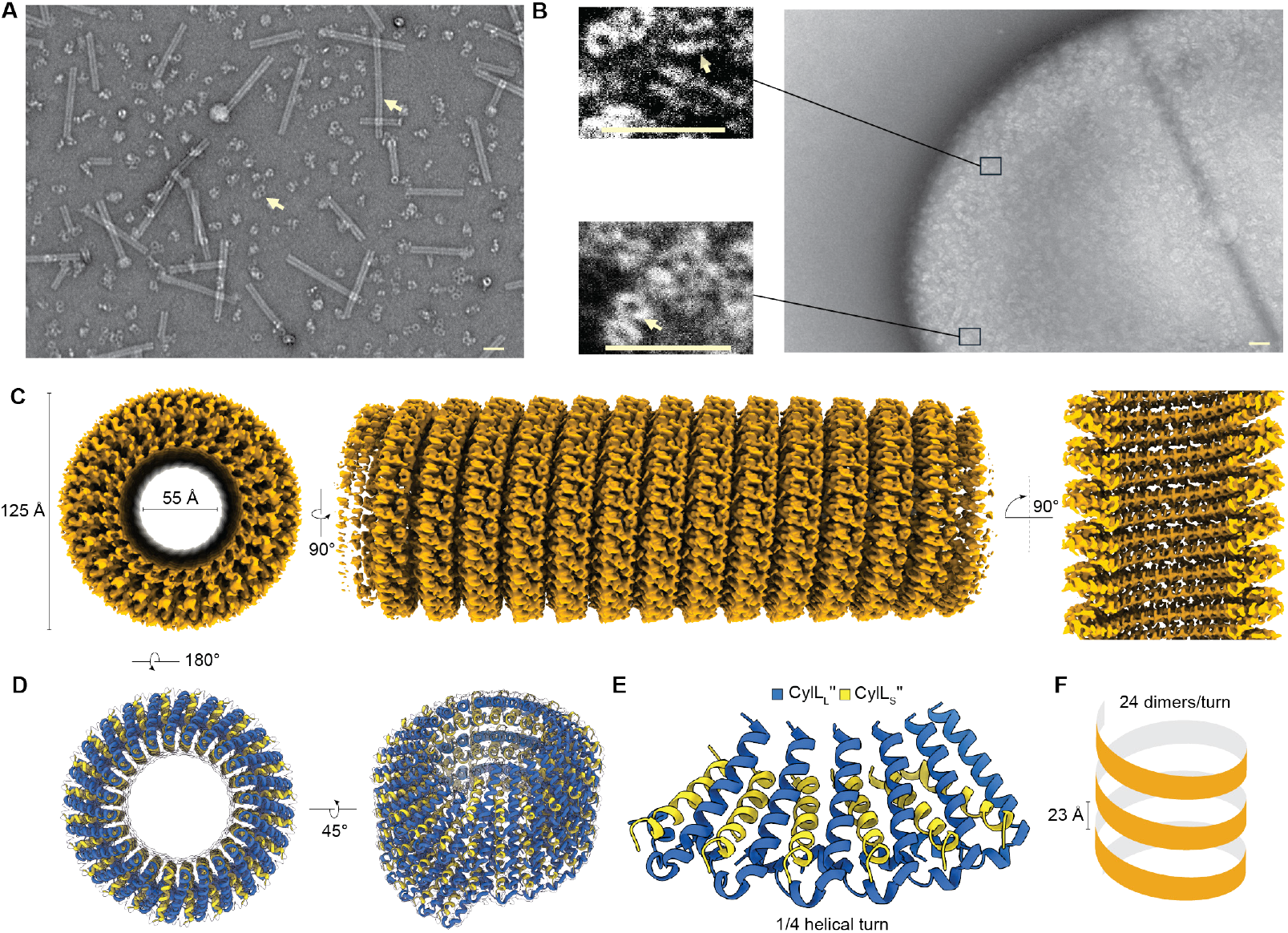
High-resolution cryo-electron microscopy structure map and the model of the cytolysin nanotube. A) Negatively stained samples of CylL_S_″: CylL_L_″ in a 1:1 ratio analyzed by TEM. Arrows indicate the two classes of features: rings and tubes. B) Negatively stained *L. lactis subsp. cremoris* after treatment with 1:1 CylL_S_″:CylL_L_″. Insets provide a magnified view of the highlighted region. The contrast was linearly increased. Colored arrows indicate nanotube and ring features. For both A) and B) the scale bars represent 50 nm. C) A cryo-EM map of the superhelical assembly with a resolution of 2.0 Å. D) Map-model overlay showing 72 units of CylL_L_″ and 72 units of CylL_S_″ fitting within the map. E) Six consecutive dimers composed of CylL_S_″ (yellow) and CylL_L_″ (blue) modelled into one quarter of a helical turn of the map. F) Cartoon of the superhelix formed during assembly. The helical pitch is equal to 23 Å and 24 dimers comprise each turn.

No high-resolution structural data has described the interaction between two different lanthipeptides. Therefore, cryo-electron microscopy (cryo-EM) was used to probe the structure of cytolysin assemblies at near-atomic resolution. Two-dimensional classification revealed tubes and ring-shaped structures under the conditions used for self-assembly and vitrification (see Supplementary Information). The tubes possessed helical symmetry, which allowed the reconstruction of a 3D map of the assembled form of cytolysin at a resolution of 2.0 Å (Figs. 2C, S7, and S8). The highly helical nature of the cytolysin nanotubes are reminiscent of the helices in the pore-forming cholesterol-dependent protein cytolysins and gasdermins (*29, 30*) and the ESCRT-III-like polymers that remodel bacterial membranes (*31-33*). The density map showed a left-handed helix with a pitch of 23 Å that formed a nanotube with a constant internal diameter of 55 Å (Fig. 2C). Model building revealed a shorter peptide and a longer peptide with mostly α-helical backbone conformations (Fig. 2D and S8). The density map possessed morphology that deviated from α-helical in areas that contained the lanthionine/methyllanthionine rings and dehydroamino acids, consistent with the atypical backbone conformations imbued by side chain modifications (*34*). Within a single dimer, CylL_S_″ forms one continuous α-helix while two helical regions bridged by a jackknife-like hinge comprises CylL_L_″ (Fig. 2E). The helical axis of CylL_S_″ forms acute angles with the two helical regions of CylL_L_″, facilitating the interdigitation of branched hydrophobic side chains spanning the lengths of both subunits (Fig. 2E) and supporting additional contacts with adjacent dimers. The constructed model is composed of dimers of CylL_S_″ and CylL_L_″, which arrange into a helix that contains 24 dimers per turn (Fig. 2F). The top and bottom faces of the helix and the exterior surface are composed of hydrophobic moieties while the lumen is relatively less hydrophobic (Fig. S9). This pattern is consistent with the membrane activity of these assemblies.

A two-dimensional map of the ring-shaped assemblies showed that they did not possess helical symmetry. A low-resolution 3D map demonstrated that the ring was cylindrical, ∼69 Å long and ∼120 Å in diameter with an internal diameter of ∼55 Å (Fig. S7). Assuming that the ring-shaped feature is a cyclic array of ∼24 CylL_S_″ and CylL_L_″ dimers, it would account for the 130 kDa assemblies observed by SDS-PAGE (Fig. S4C).

The predominantly helical backbones of CylL_L_″ and CylL_S_″ are similar to those determined in the solution phase through NMR characterization of the individual subunits (*8*), suggesting that the lanthionine rings serve to template the backbone into an assembly-ready conformation (Fig. 3A). The backbone of CylL_L_″ resembles a jackknife because the two helices containing the N- or C-termini are separated by a hinge region that places them at an acute angle. Each α-helix is conformationally seeded by a (methyl)lanthionine ring near the terminus (*35*). The central lanthionine ring enforces the hinge between the two helical axes. The backbone of CylL_S_″ is entirely helical in the nanotubes and does not contain the disordered region observed in the NMR structures (Fig. 3B). The transition of disordered to helical results in the formation of more intrachain hydrogen bonds, which may help pay the enthalpic cost of desolvation and drive assembly formation.

**Fig. 3.**
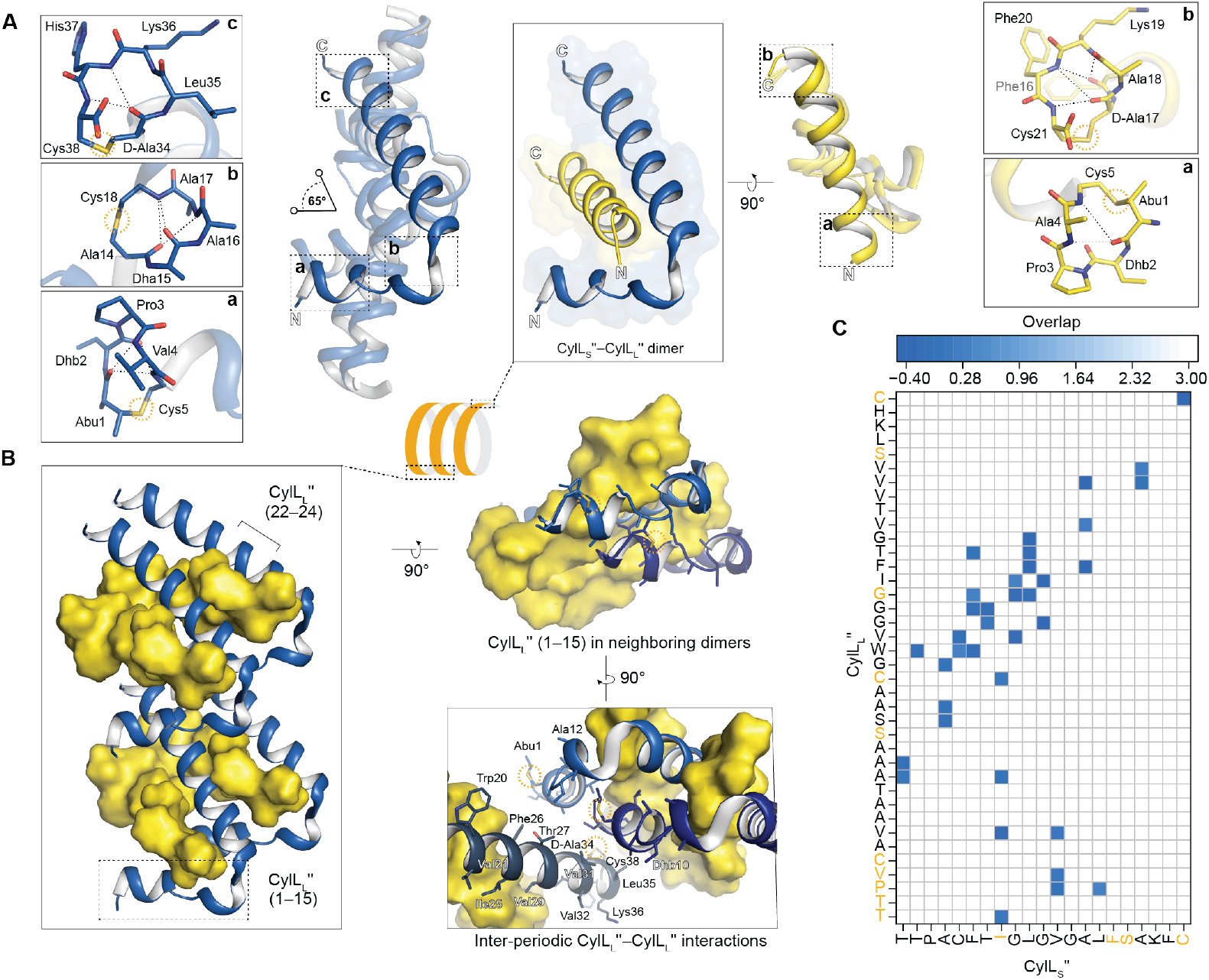
Structural analysis of the cytolysin oligomeric helical assembly. CylL_S_″ is shown in yellow and CylL_L_″ in blue. A) Details of the structure of the CylL_L_″-CylL_S_″ dimer. Overlay of the low energy structures of CylL_L_″ in the solution-phase (high-transparency) with the structure in the assembly (low-transparency). The structures of (methyl)lanthionine rings are shown as insets. Rings are labelled alphabetically starting at the N-terminus. B) CylL_L_″ scaffolds the assembly through a network of interactions conferred by the N-terminal helix of CylL_L_″ (residues 1–15; one of the four such segments in the figure is boxed). This section of CylL_L_″ is involved in intra- and inter-dimer contacts. It also participates in inter-periodic contacts with molecules in the adjacent turn. C) A two-dimensional heatmap illustrating important contacts between CylL_L_″–CylL_S_″. Grid squares are colored according to overlap, which was calculated in ChimeraX (*36*). Negative values are associated with stronger contacts. Positions highlighted in orange when mutated to Ala caused a loss in both antibiotic and hemolytic activity.

Based on the atomic model, assembly is hydrophobically driven and coordinated by the relative positions of branched hydrophobic side chains, which in turn are dictated by the (methyl)lanthionine rings. The α-helix formed by CylL_S_″ fills a gap between two CylL_L_″ units flanked on either side by Gly22, Gly23, and Gly24 of CylL_L_″ (Fig. 3B and Fig. S10). Branched hydrophobic side chains between the N- and C-terminal rings of CylL_S_″ make contacts with CylL_L_″ units on each flank (Fig. 3B and Fig. S10). The C-termini of adjacent CylL_L_″ chains come closer together as their backbones move away from the gap-filling CylL_S_″ unit. This proximity promotes the intercalation of the branched hydrophobic residues of two adjacent CylL_L_″ peptides (Fig. 3B and Fig. S10). The shorter α-helix of CylL_L_″ (residues 1–15) makes contacts with adjacent dimers within a superhelical period and makes contacts with subunits inter-periodically (Fig. 3B). Two CylL_L_″ helices composed of residues 1–15 from adjacent dimers create hydrophobic pockets that accommodate branched hydrophobic side chains found in CylL_S_″ (Fig. 3B), and CylL_L_″ chains interact inter-periodically through the C- and N-terminal lanthionine-enclosed rings in each chain (Fig. 3B). The overall result is a highly intertwined architecture that resembles a rope braid.

The structure was used to gain more insights into the structure-activity relationship of cytolysin (*28*). The CylL_L_″ 1–15 helix forms a network of crucial interactions that hold the assembled structure together. Consistent with this central role, mutations that removed the N-terminal methyllanthionine-containing ring of CylL_L_″ reduced assembly stability and decreased bioactivity (Fig. 4C and Fig. S6). In contrast, the N-terminal ring of CylL_S_″ makes few contacts with adjacent chains and is presented on the solvent exposed face of the assembly. Indeed, mutating this ring did not have a severe impact on assembly stability or bioactivity (Fig. 3C and Fig. S6). Mutating non-ring forming residues to Ala generally did not severely impact antibiotic and hemolytic activity with a handful of exceptions (*28*) that coincide with positions that form central contacts in the assembly. CylL_S_″ Phe16 interdigitates with residues in adjacent CylL_S_″ chains (Fig. S10) and CylL_S_″ Ile8 slots into a hydrophobic cavity generated by adjacent CylL_L_″ 1– 15 helices. Mutating these residues to Ala severely reduced antibiotic and hemolytic activity (*28*). The high-resolution structure also explains why CylL_L_’ and CylL_S_’ are inactive because adding six amino acids to the N-termini of either CylL_L_″ or CylL_S_″ would interfere with the tight packing observed in this section of the structure. In general, the ring forming residues that confers the secondary structures of CylL_S_″ and CylL_L_″ are more important for bioactivity than the side chains of individual residues that are not involved in lanthionine ring formation (Fig. 3C). The structure also agrees with the activity of WT CylL_S_’’ with CylL_L_’’ that carries an NBD group on Lys36. The side chain of Lys36 does not make any critical contacts and the structure can accommodate the NBD group on its periphery, or the NBD group can interdigitate with nearby branched hydrophobic side chains. An increase in NBD fluorescence was observed immediately after combining NBD-labeled CylL_L_’’ with WT CylL_S_″ (Fig. S11), consistent with the NBD group interacting with nearby hydrophobic residues (*37*).

**Fig. 4.**
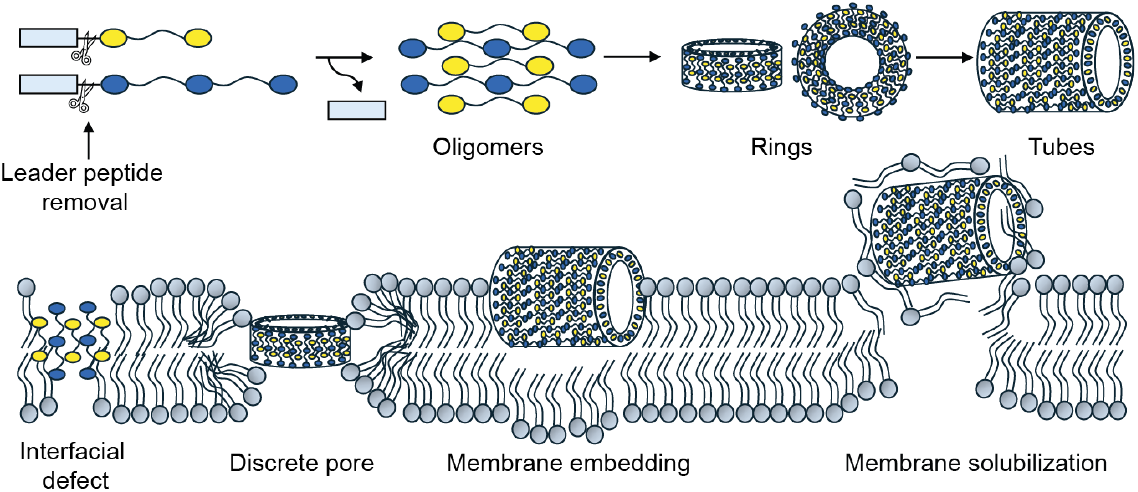
The mechanism of action of cytolysin. Leader peptide cleavage initiates oligomerization which progresses to yield hydrophobic rings and tubes that attack cellular membranes. Each oligomeric state has the potential to disrupt the membrane through different means. Smaller species may produce interfacial defects similar to antimicrobial peptides. Rings can form discrete pores similar to pore-forming protein toxins. Tubes can embed into the outer layer of the membrane causing permeabilization through membrane solubilization similar to amyloid forming peptides.

Our findings led to the development of heuristics for biasing genome mining towards α-helical lanthipeptides and uncharacterized cytolysin-like BGCs. The α-helical conformations of CylL_S_″ and CylL_L_″ are encoded into their sequences by the S/T(X)_3_C motifs. These motifs predict *i* to *i*+4 linkages which are known to template α-helical backbone conformations (*38*), especially when located at the N- and C-termini of a sequence (*35, 39*) (Fig. S12). Using genome mining, over 100 non-redundant substrates of class II lanthipeptide synthetases that possess residues primed to make *i* to *i*+4 linkages at the N- and C-termini were discovered (Data S1 and Fig. S12). The majority possess sequences consistent with CylL_S_″-like ring patterns and resemble the lanthipeptides cerecidin and bibacillin 1 (*40, 41*). Some of these are encoded in BGCs that contain a substrate with a CylL_L_″-like ring pattern, suggesting cytolysin-like synergistic activity. The taxonomic distribution of the cytolysin-like clusters was evaluated with a phylogenetic tree (Fig. S12). Most sequences are from Bacillota, but examples were found in Actinomycetota, Pseudomonadota, and Myxococcota. These bacteria occupy ecological niches split evenly between commensals and free-living species. Considering that some of these microbes play roles in human health and disease, the harmful impact of enterococcal cytolysin-like systems may extend beyond alcoholic hepatitis.

The enterococcal cytolysin was first described in the 1930s and its deleterious effects on human health have been long known (*42*). Its mode of action and the mechanism of its high potency against both bacterial and eukaryotic cells was thus far unresolved. The data in this study show that unlike other highly potent pore-forming natural products that recognize specific molecular targets in the membrane of susceptible cells (*17, 43-46*), the enterococcal cytolysin derives its potency by recognition of its partner to make highly ordered structures that perturb both bacterial and eukaryotic cells. The only requirement for assembly into bioactive oligomers is the presence of both subunits of the toxin, explaining its cross-kingdom activity as well as the short lag phase observed in its activity (Fig. S1A). Our findings suggest a mechanism in which different oligomeric forms of cytolysin possess different activities. Smaller oligomers, such as the rings observed in Fig. 2A, could insert into the membrane forming defects or discrete pores (Fig. 4) (*20*). Larger oligomers, such as the tubes observed in Fig. 2A, may cause membrane disruption through a detergent-like mechanism that proceeds through membrane embedding, lipid extraction, and membrane solubilization (Fig. 4) (*47*). To gain more insights into the molecular weight of the active species, we performed in-vitro dialysis experiments, which revealed that the active species has a molecular weight exceeding 10 kDa (*48*) (Fig. S13).

The structure and mechanism of cytolysin are intermediate between those of membrane-active peptides and proteins. Our structure places the enterococcal cytolysin within a small but growing family of hetero-oligomeric pore-forming proteins that include the membrane attack complex (*49, 50*) and the RCD-1 gasdermin pore (*51*). The MoA proposed here for cytolysin has similarities and differences to the model that was recently proposed for candidalysin, a 31-residue unmodified cationic peptide that functions as a virulence factor in *Candida albicans* (*52*). Candidalysin forms octameric bundles of α-helices that function as monomers in the polymerization of the peptide into linear, branched or looping chains that insert into phospholipid membranes with little backbone perturbation (*53, 54*). Compared to candidalysin, cytolysin self-assembly is considerably more structured and rigid and involves two post-translationally modified peptides. For the cytolysin assembly to occur, a leader peptide must be removed, making the proteases that cleave off the leader peptide potential targets for therapeutic intervention (*27*). In addition, the thioether crosslinks in CylL_L_″ and CylL_S_″ are critical for assembly suggesting that the enzyme CylM (*55*) that installs these structures is a potential target. Structural and mutational studies also demonstrate that the N-terminal ring of CylL_L_″ is central to bioactivity and assembly. A single nanotube would have these features exposed to solvent at the ends of the tube, making this ring also a promising target for neutralizing antibodies.

In summary, the high-resolution structure of the enterococcal cytolysin assembly provides a unique glimpse into how post-translation modifications can program bioactivity into peptide sequences. This structure provides a starting point for discovering or designing new bioactive molecules and self-assembling systems.

## Acknowledgments

The authors thank Steven Hobbs and Prof. Sayee Anakk (UIUC) for providing HepG2 cells. The research presented in this manuscript was carried out in part in the Material Research Laboratory Central Research Facilities and the Core Facilities at the Carl R. Woese Institute for Genomic Biology at the University of Illinois. We thank the Brandeis Electron Microscopy Facility and the Harvard Center for Cryo-Electron Microscopy. This study is subject to HHMI’s Open Access to Publications policy. HHMI laboratory heads have previously granted a nonexclusive CC BY 4.0 license to the public and a sublicensable license to HHMI in their research articles. Pursuant to those licenses, the author-accepted manuscript of this article can be made freely available under a CC BY 4.0 license immediately upon publication.

## Funding

This study was supported by the Howard Hughes Medical Institute (HHMI) (W.A.v.d.D.).

R.M. was supported by a postdoctoral fellowship from the Life Sciences Research Foundation.

A.G.J. is supported by a grant from the Rita Allen Foundation and start-up funds from Brandeis University.

## Author contributions

Conceptualization: RM, WAvdD

Methodology: RM, WAvdD, AGJ, CG, IRR, KJDF, FFL

Investigation: RM, WAvdD, AGJ, STH, MP, CG, IRR, GL, RG, PK, PW, ET

Visualization: RM, AGJ, MP

Funding acquisition: RM, WAvdD, AGJ

Project administration: RM, WAvdD, AGJ

Supervision: RM, WAvdD

Writing – original draft: RM, WAvdD, AGJ

Writing – review & editing: RM, WAvdD, AGJ

## Competing interests

The University of Illinois and Brandeis University have filed a provisional patent application on the work described in this study.

## Data, code, and materials availability

Atomic coordinates and structure factors for the reported cryo-EM structure have been deposited with the Protein Data Bank under accession numbers 11PC and the Electron Microscopy Data Bank under accession number EMD-75911.

## Supplementary Materials

### Supplementary Materials

Materials and Methods

Figures S1 to S13

Table S1

References 56–91

Data S1 and S2

