## Supplementary figures and images for "Structure and Mechanism of a Two-component Lanthipeptide Toxin"

### 20241126_REM_bacteria_Cyl_040.tif

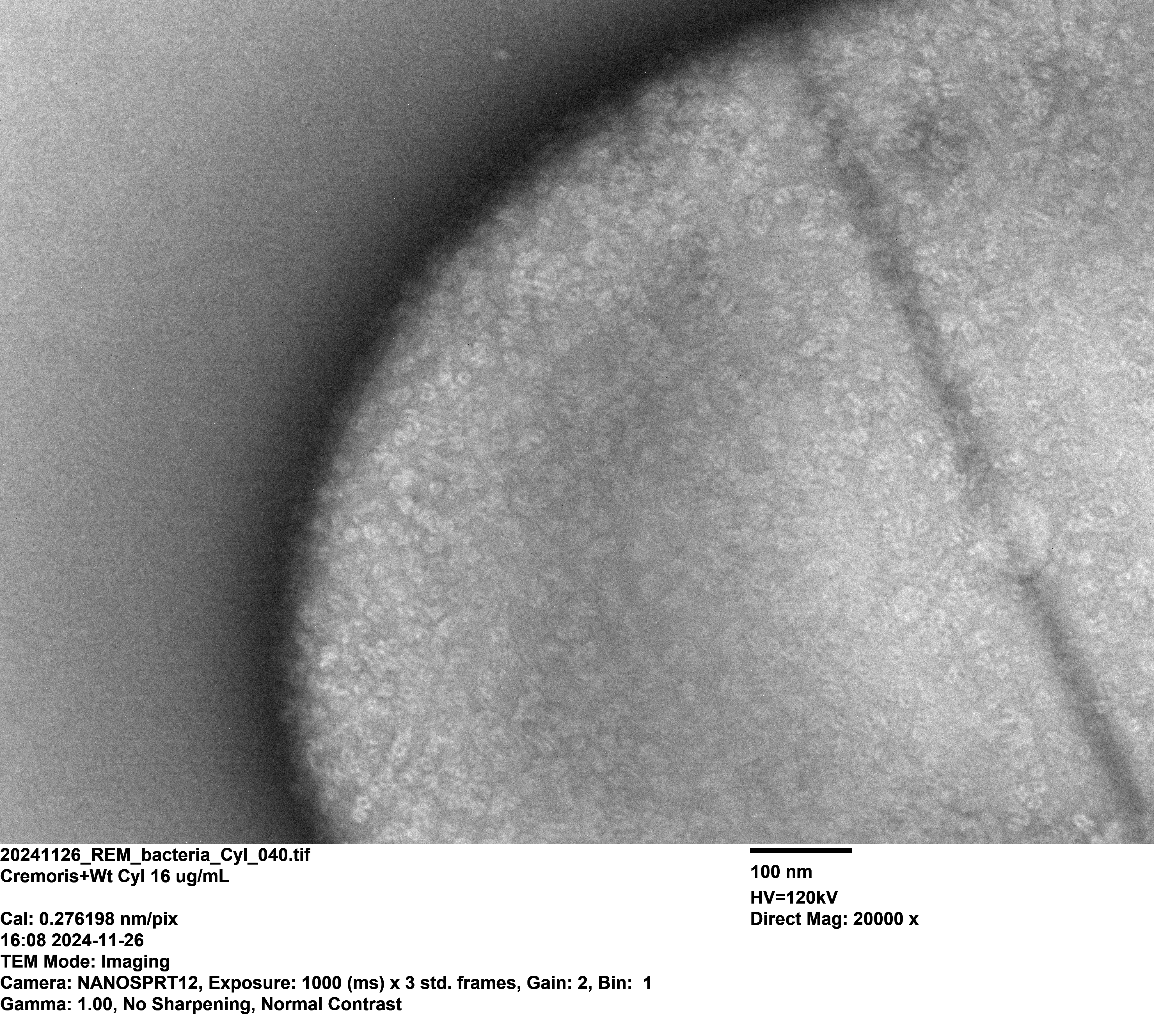

### 20250523_REM_CylLS_ovn_1_003.tif

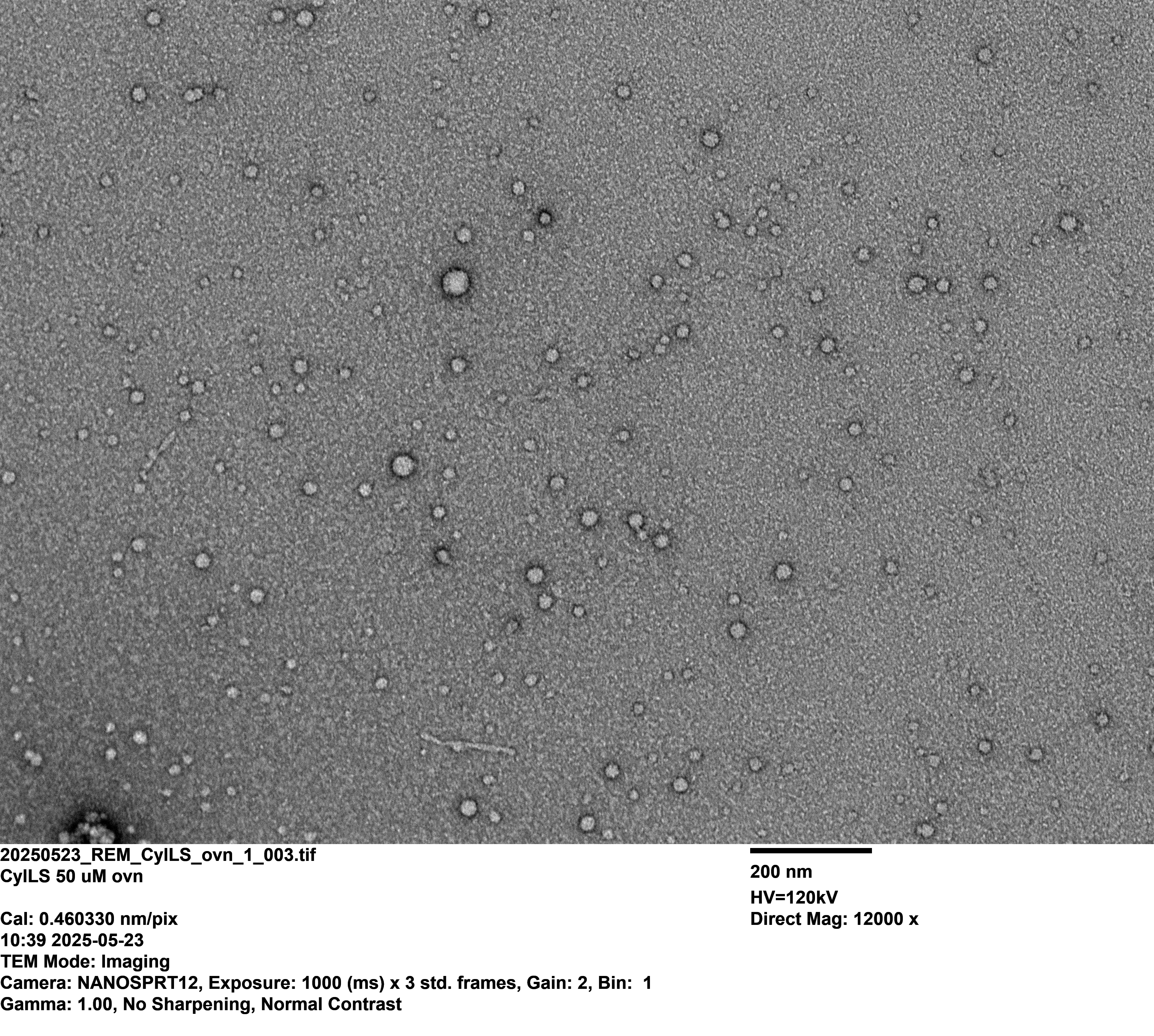

### 20250523_REM_CylLS_ovn_1_011.tif

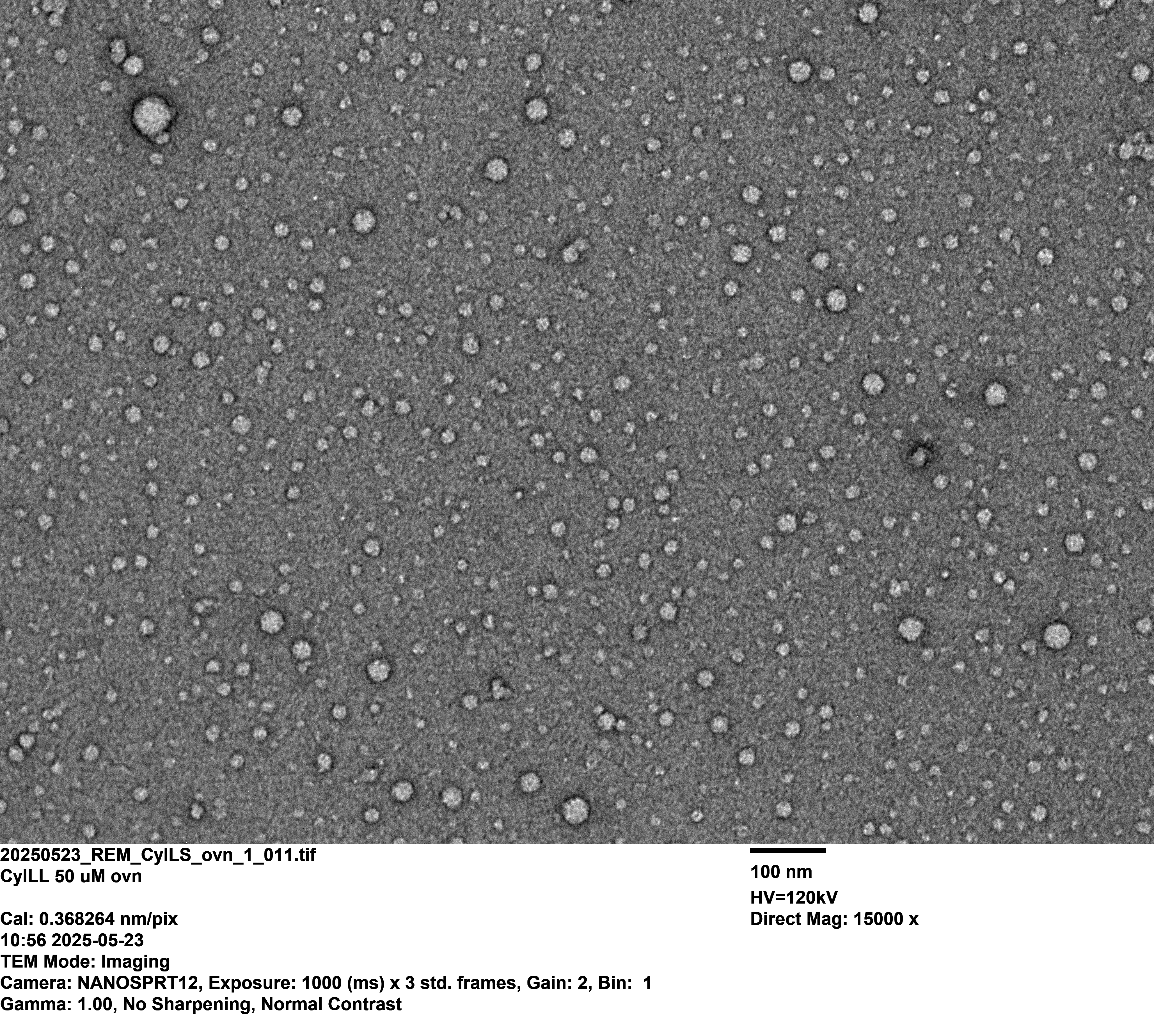

### 20250602_REM_Singleprimes_Conc_Cyl_1_003.tif

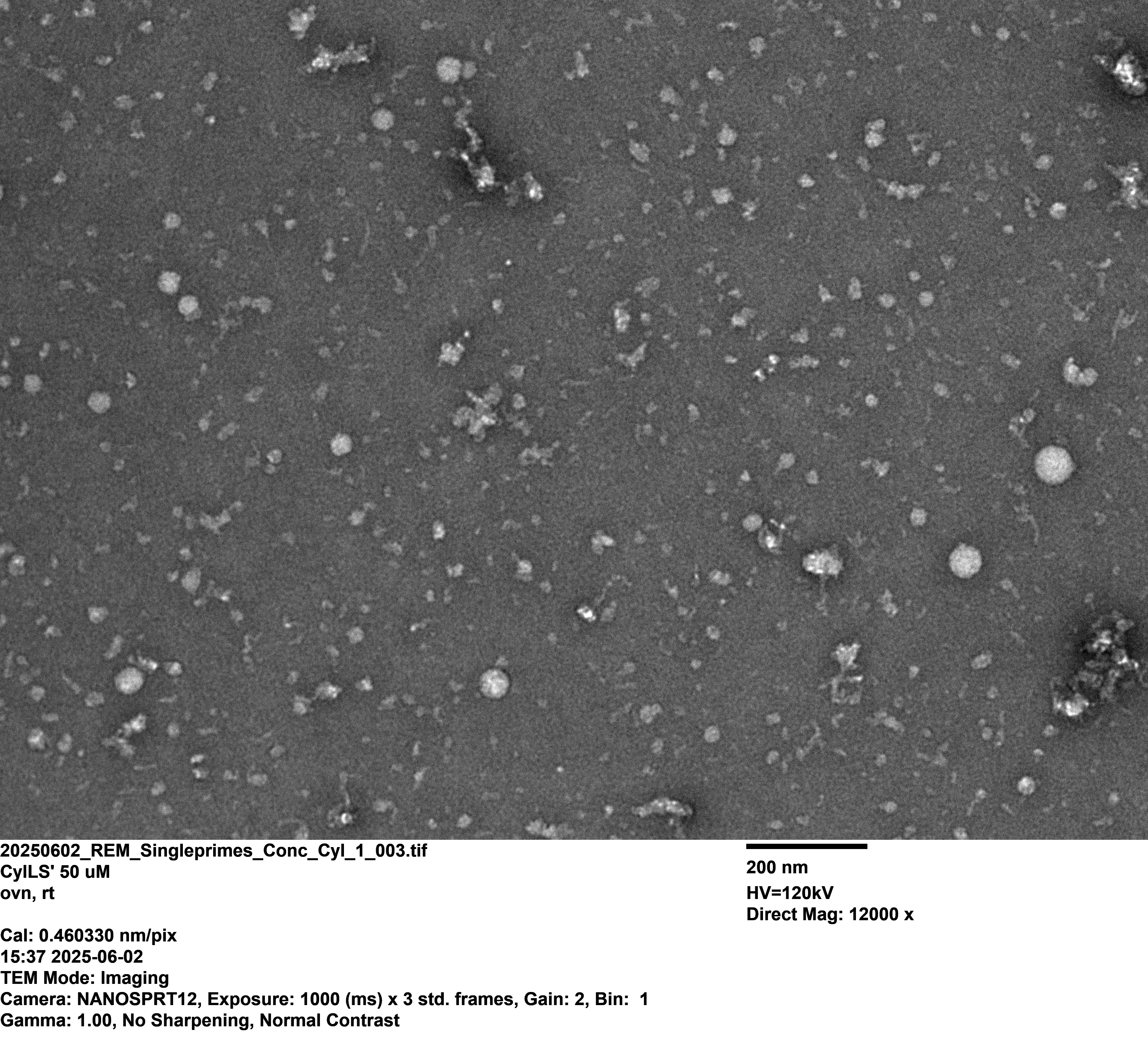

### 20250602_REM_Singleprimes_Conc_Cyl_1_010.tif

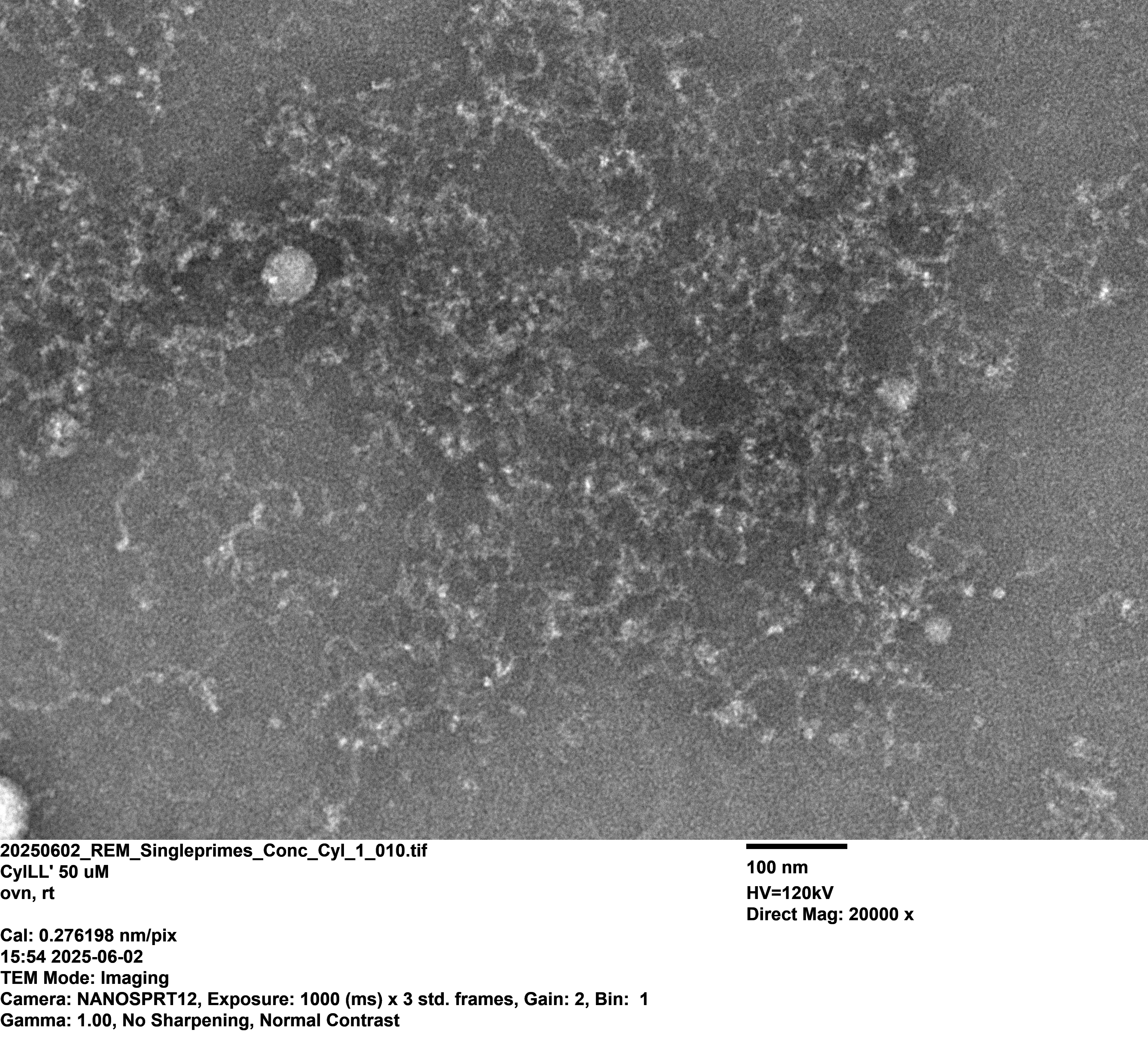

### 20250602_REM_Singleprimes_Conc_Cyl_1_013.tif

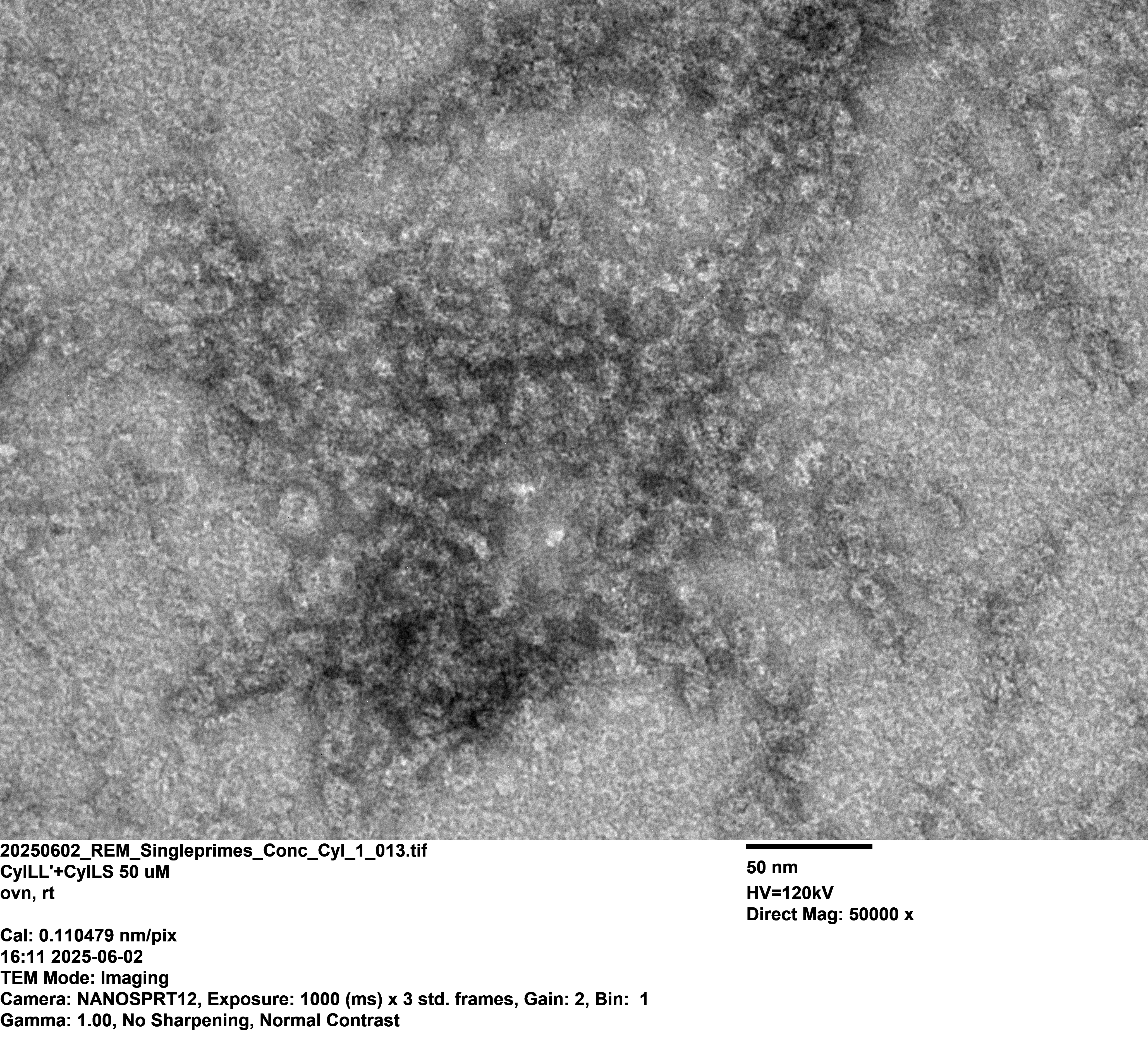

### 20250602_REM_Singleprimes_Conc_Cyl_1_019.tif

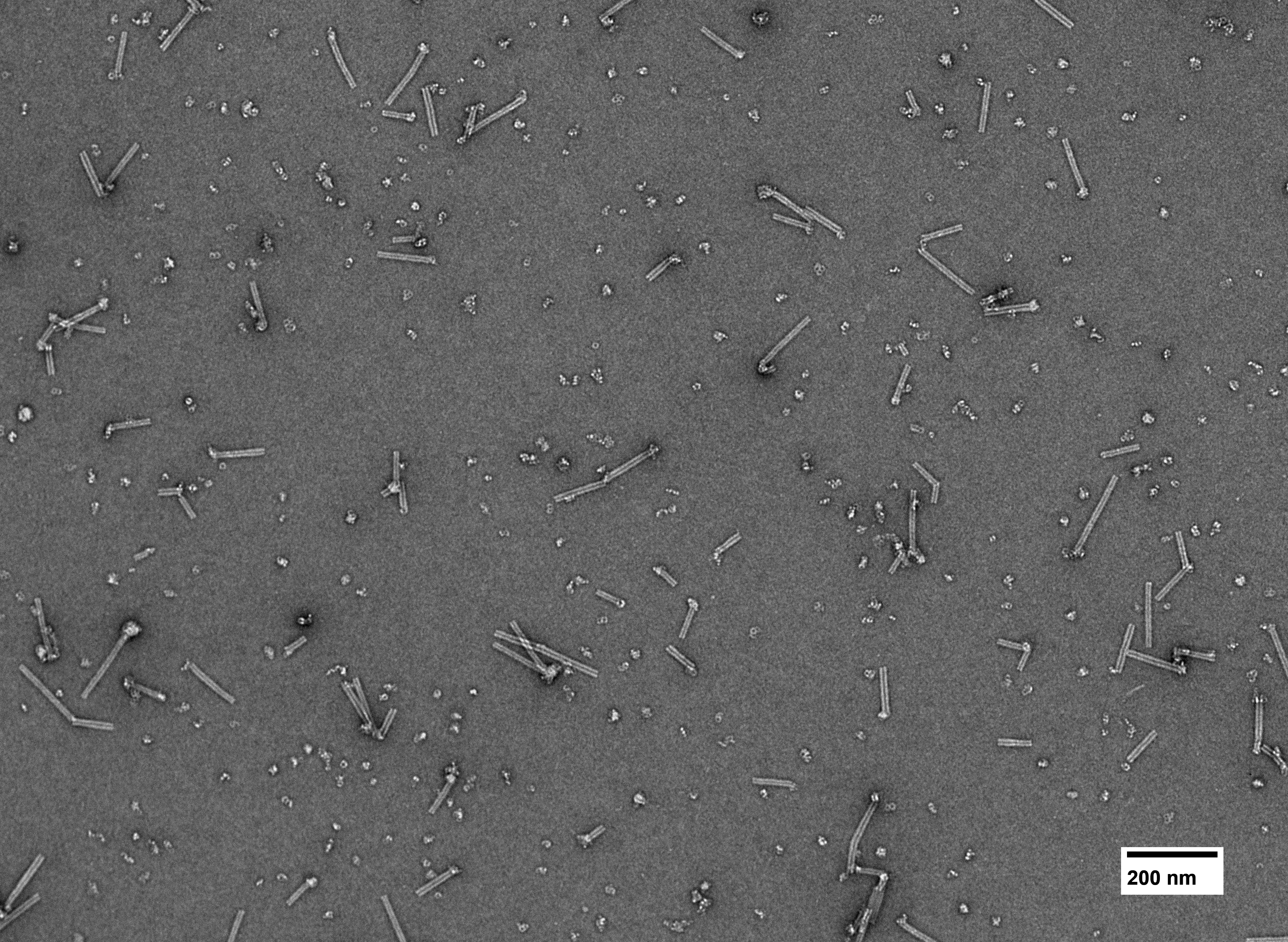

### 20250602_REM_Singleprimes_Conc_Cyl_1_020.tif

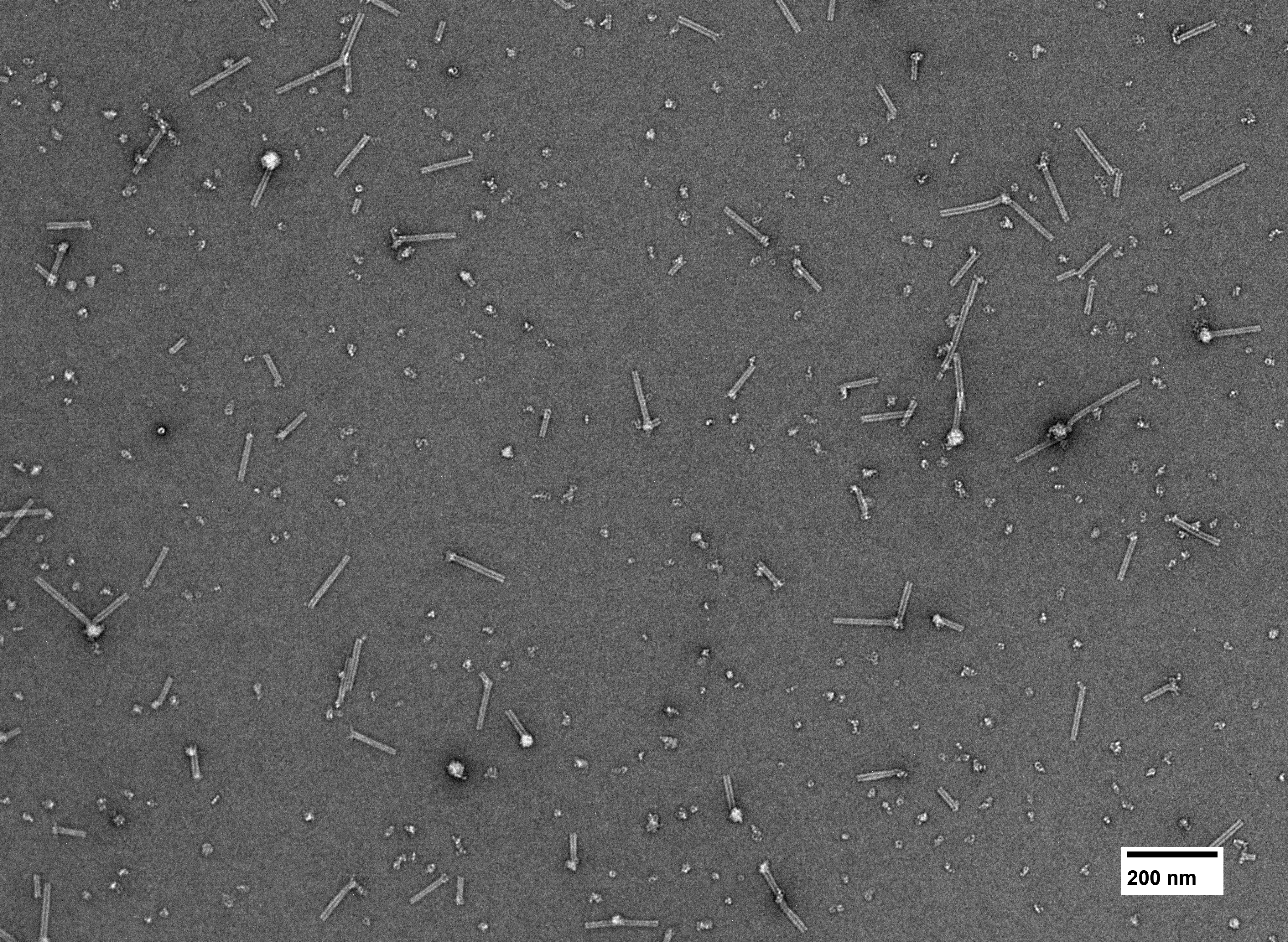

### 20250602_REM_Singleprimes_Conc_Cyl_1_022.tif

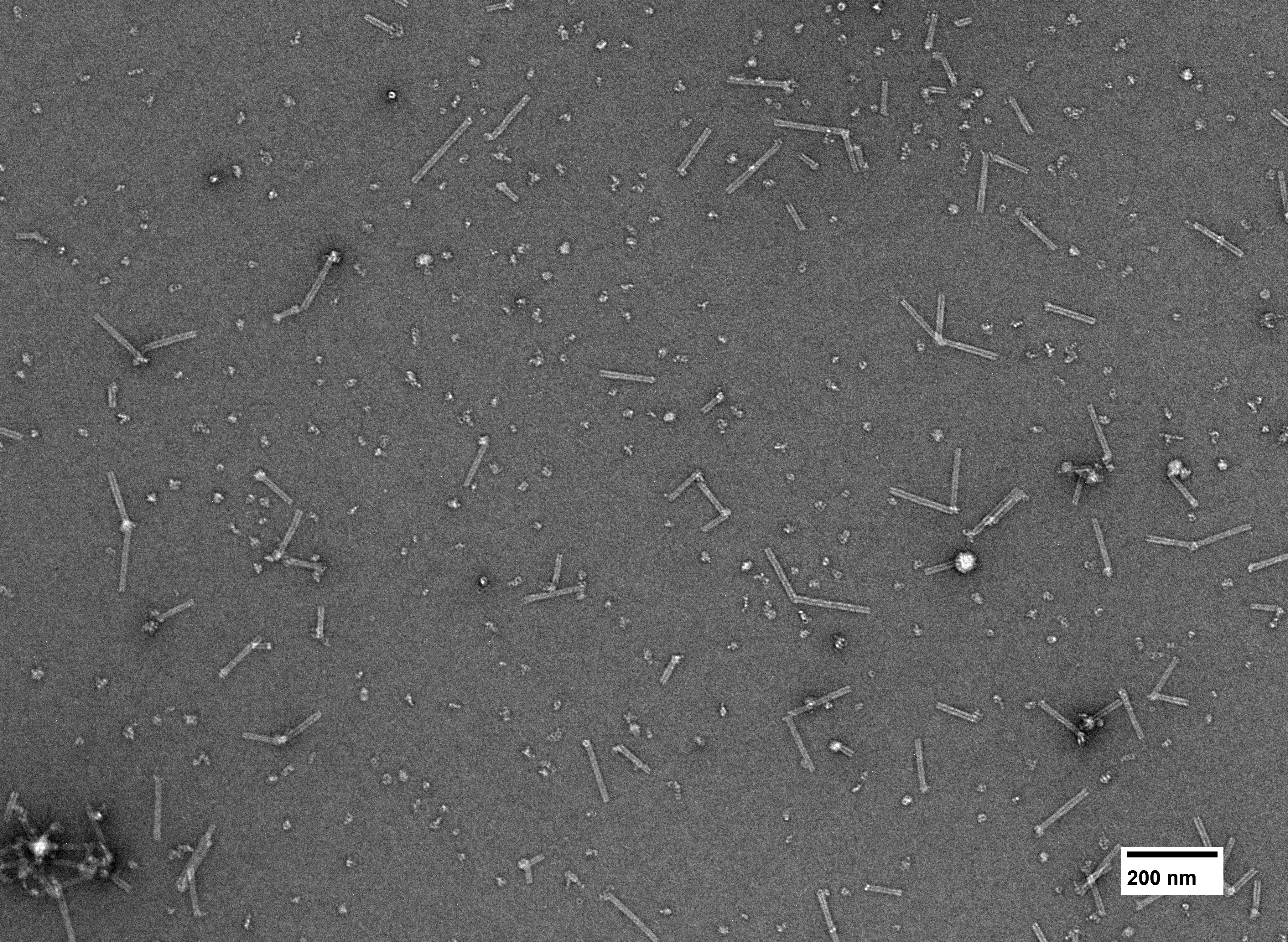

### 20250602_REM_Singleprimes_Conc_Cyl_1_023.tif

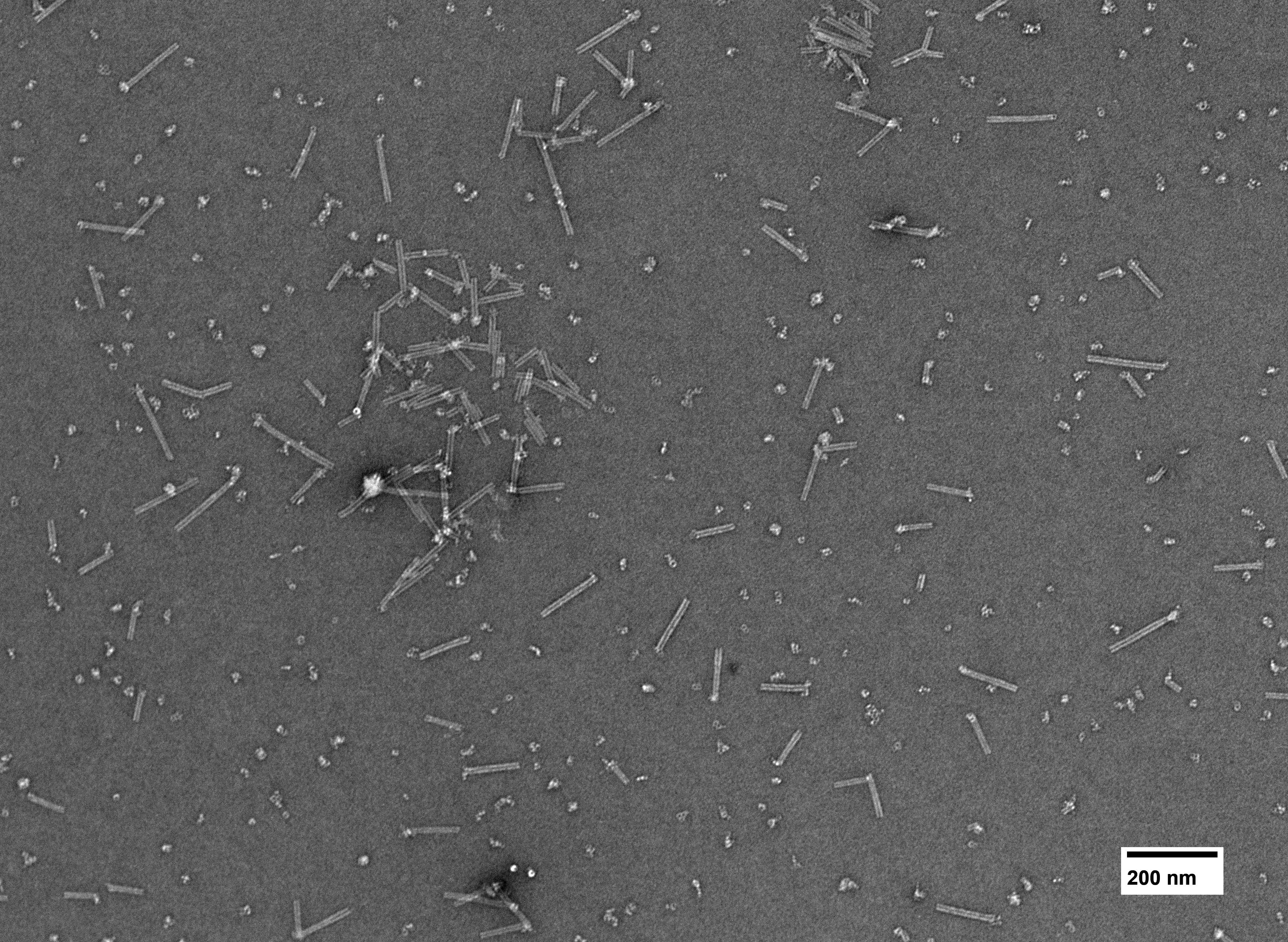

### 20250602_REM_Singleprimes_Conc_Cyl_1_024.tif

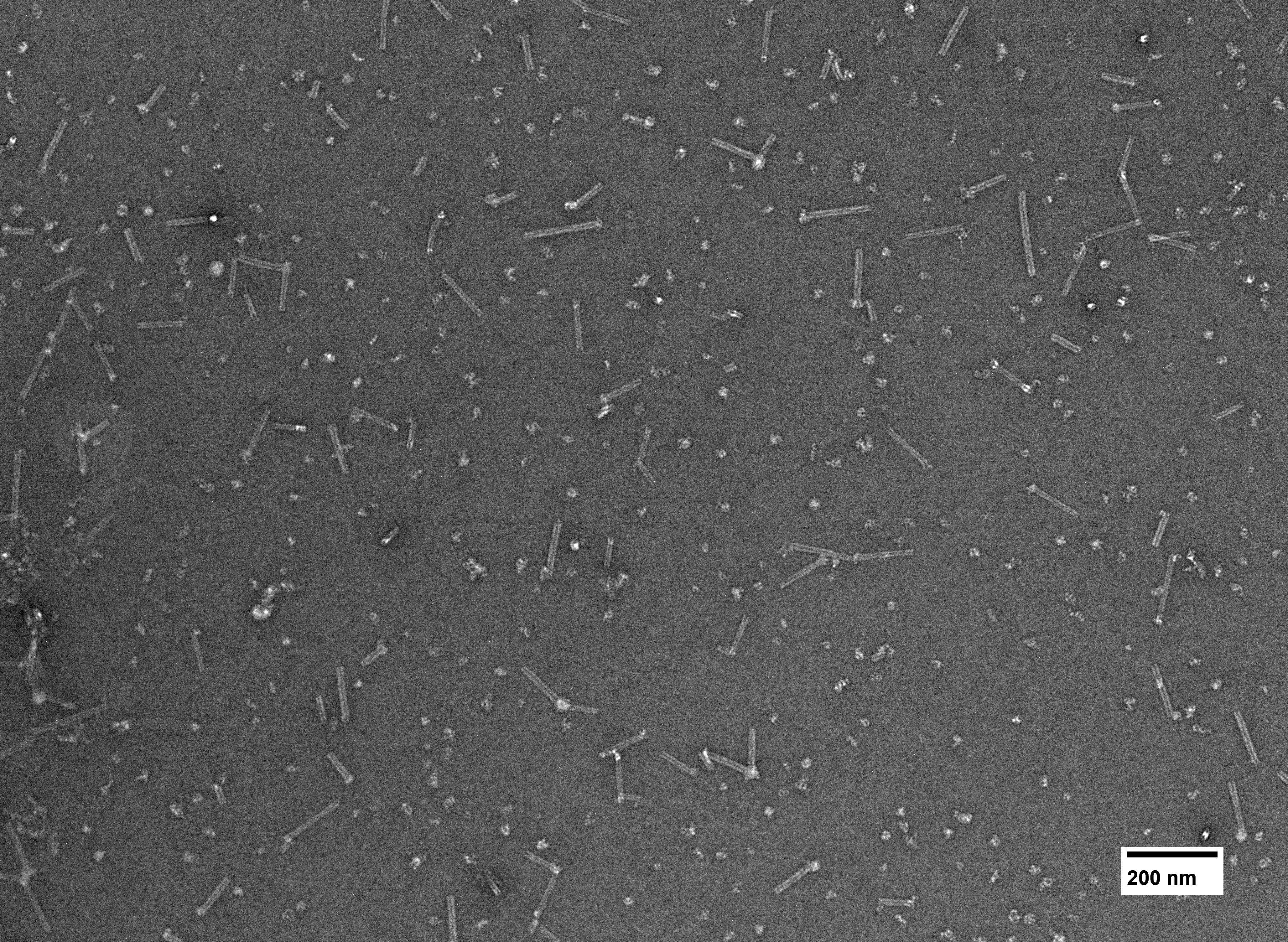

### 20250602_REM_Singleprimes_Conc_Cyl_1_026.tif

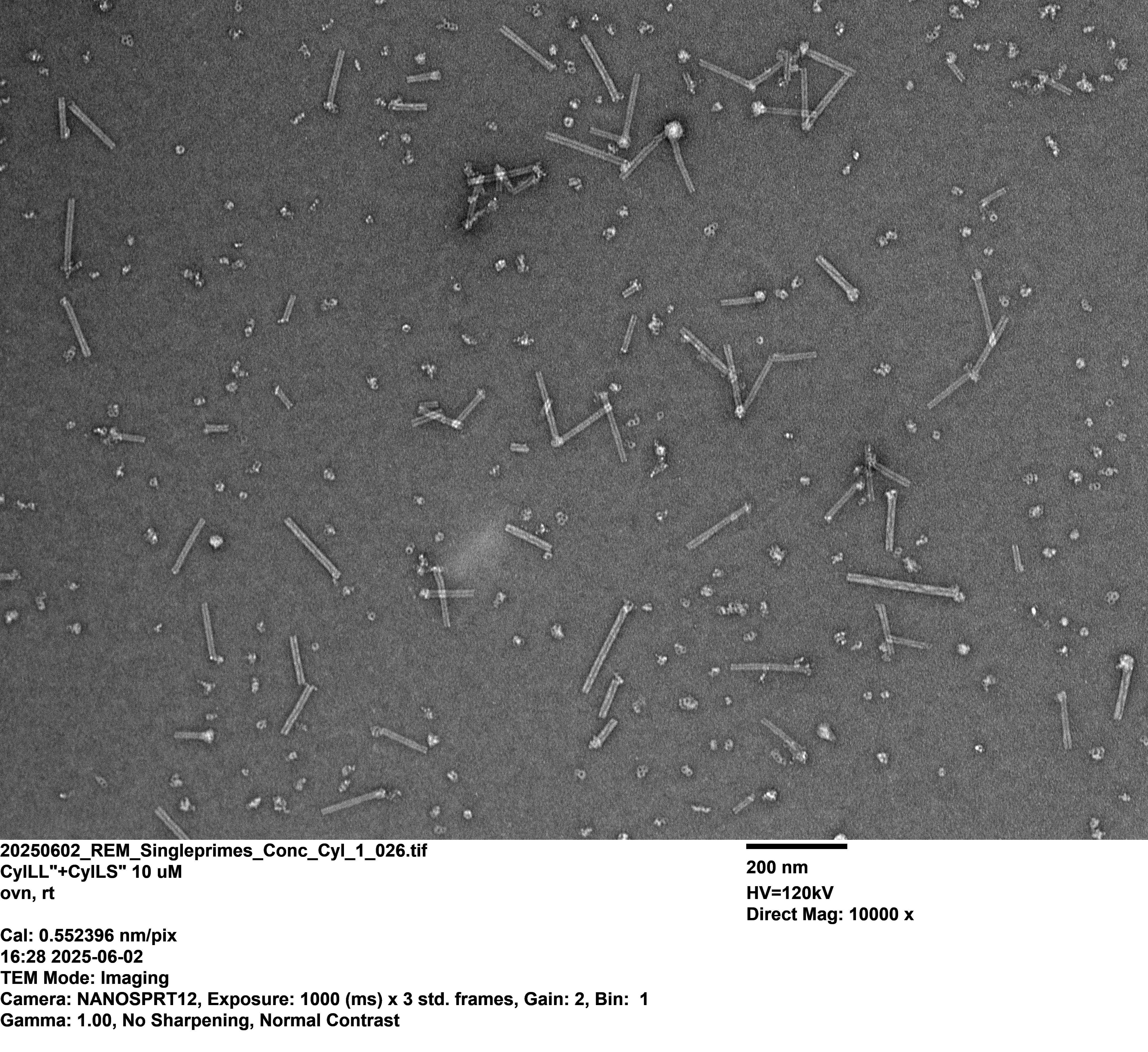

### 20250602_REM_Singleprimes_Conc_Cyl_1_028.tif

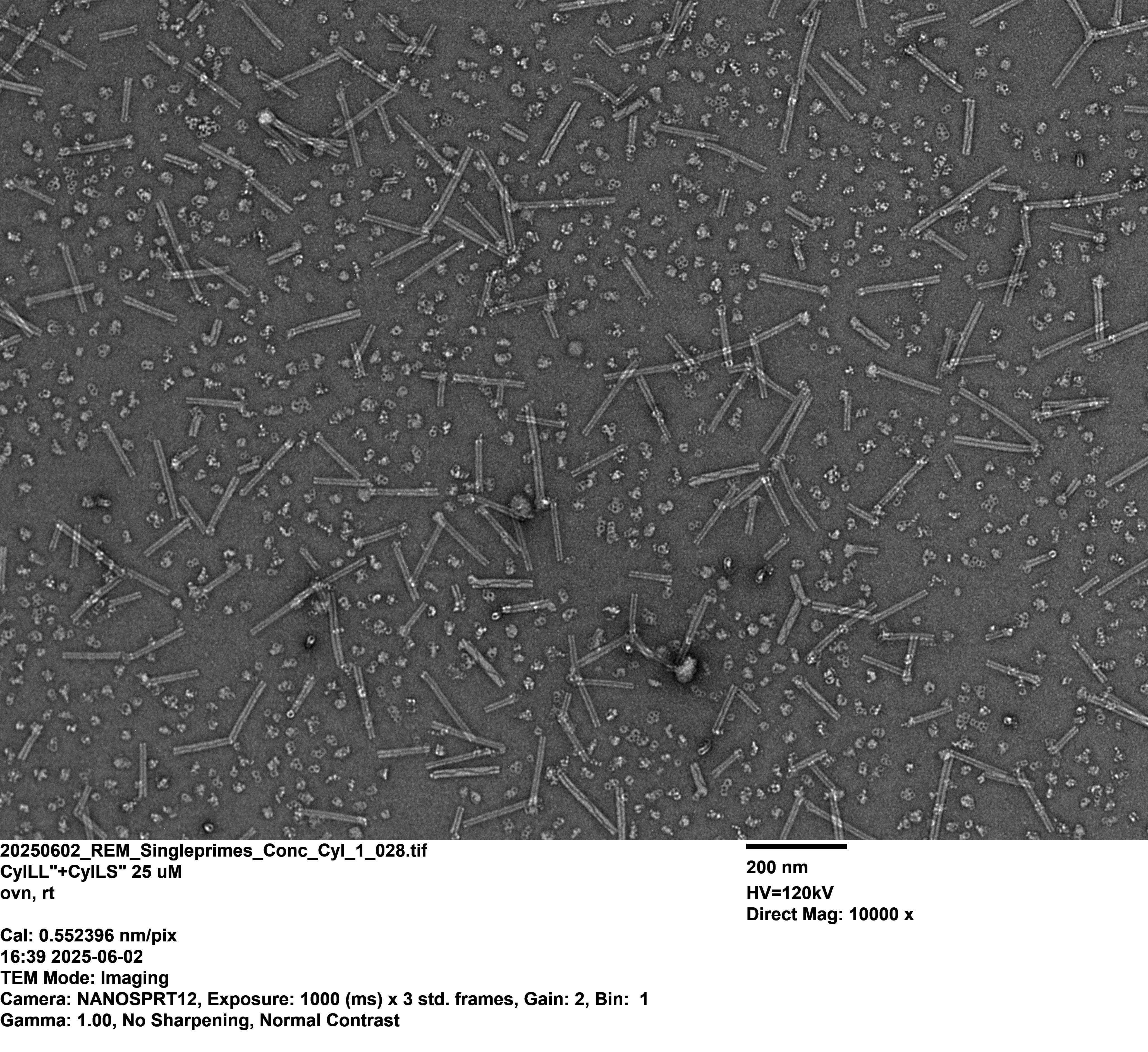

### 20250602_REM_Singleprimes_Conc_Cyl_1_029.tif

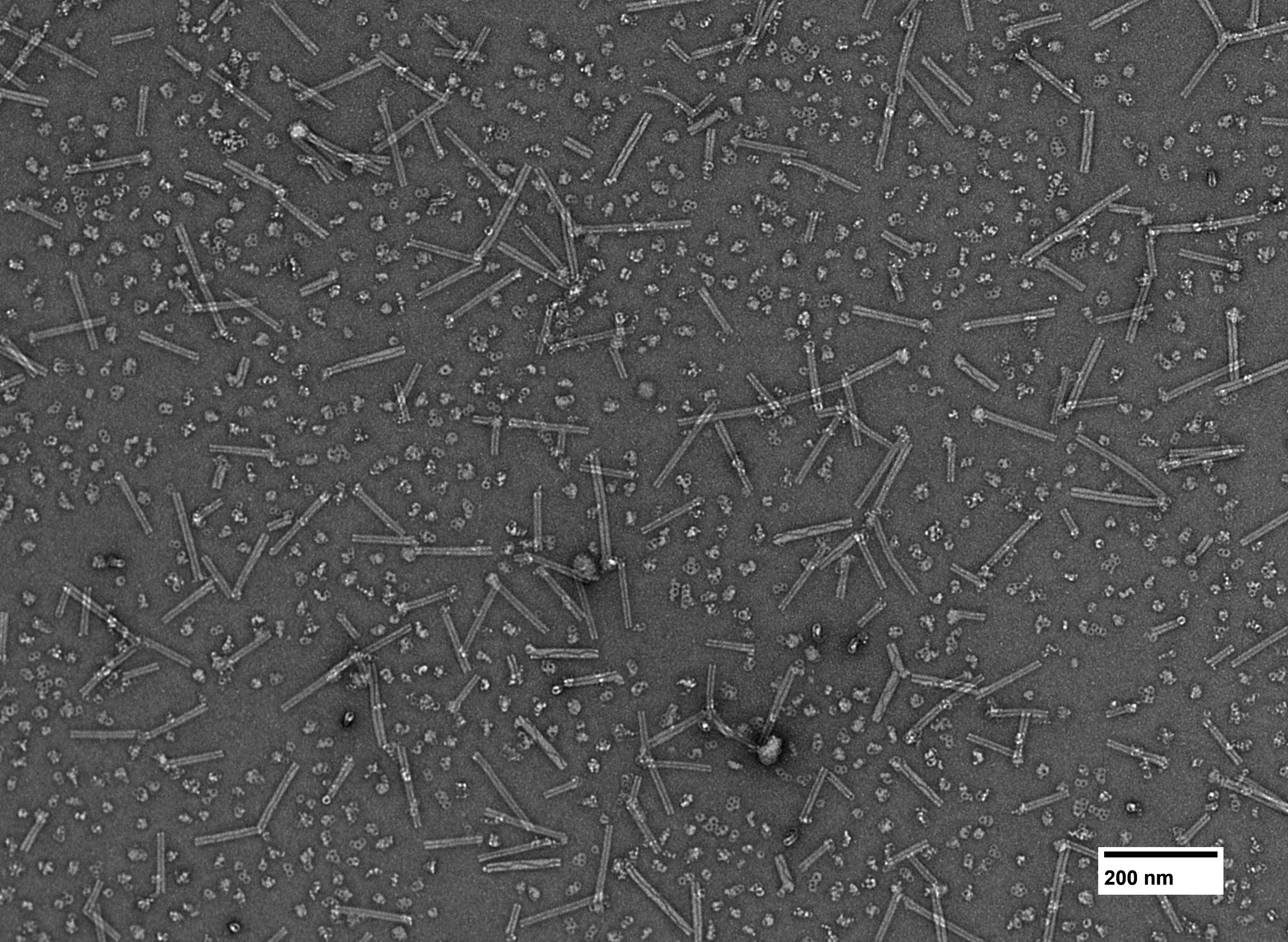

### 20250602_REM_Singleprimes_Conc_Cyl_1_030.tif

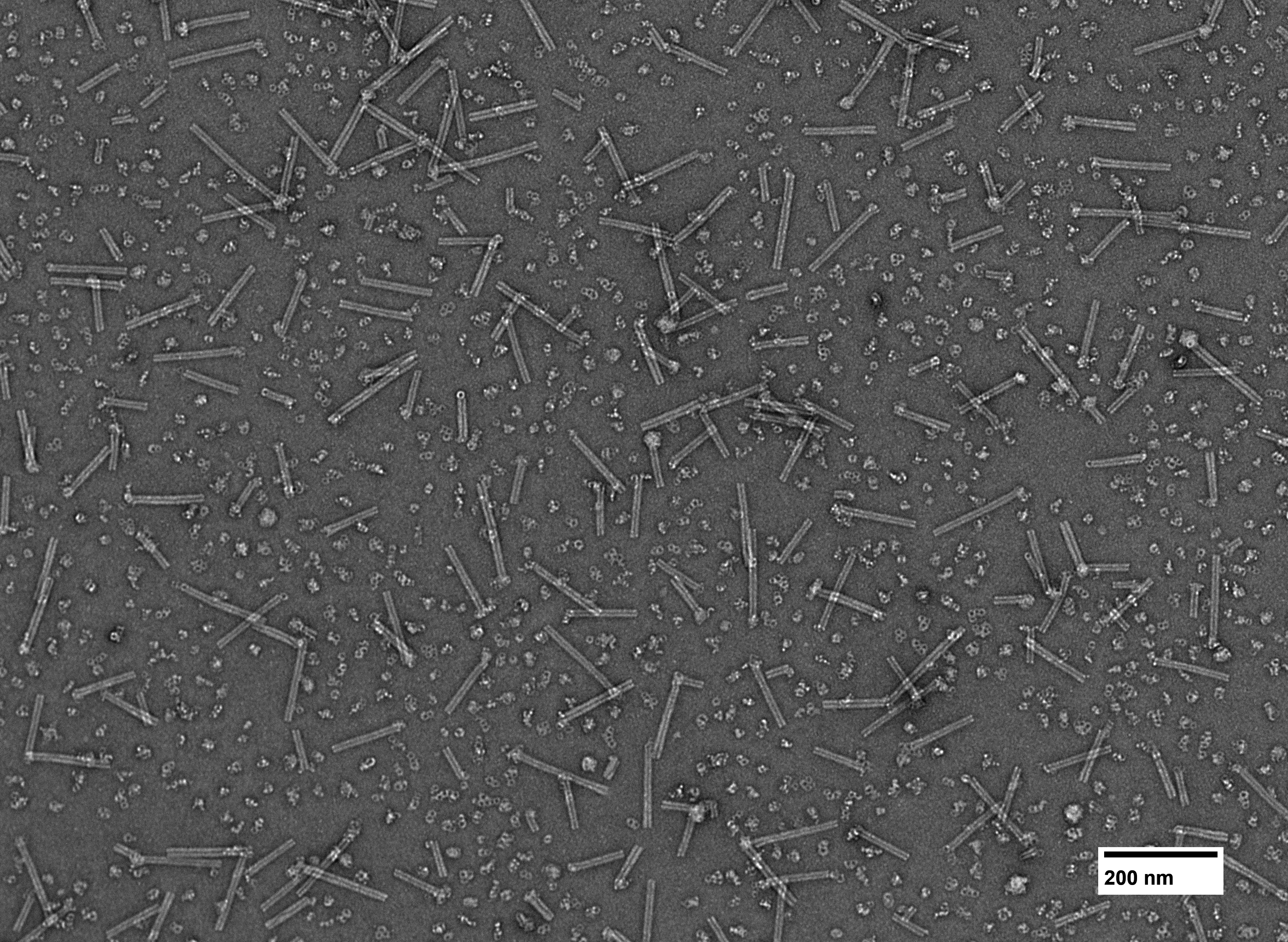

### 20250602_REM_Singleprimes_Conc_Cyl_1_031.tif

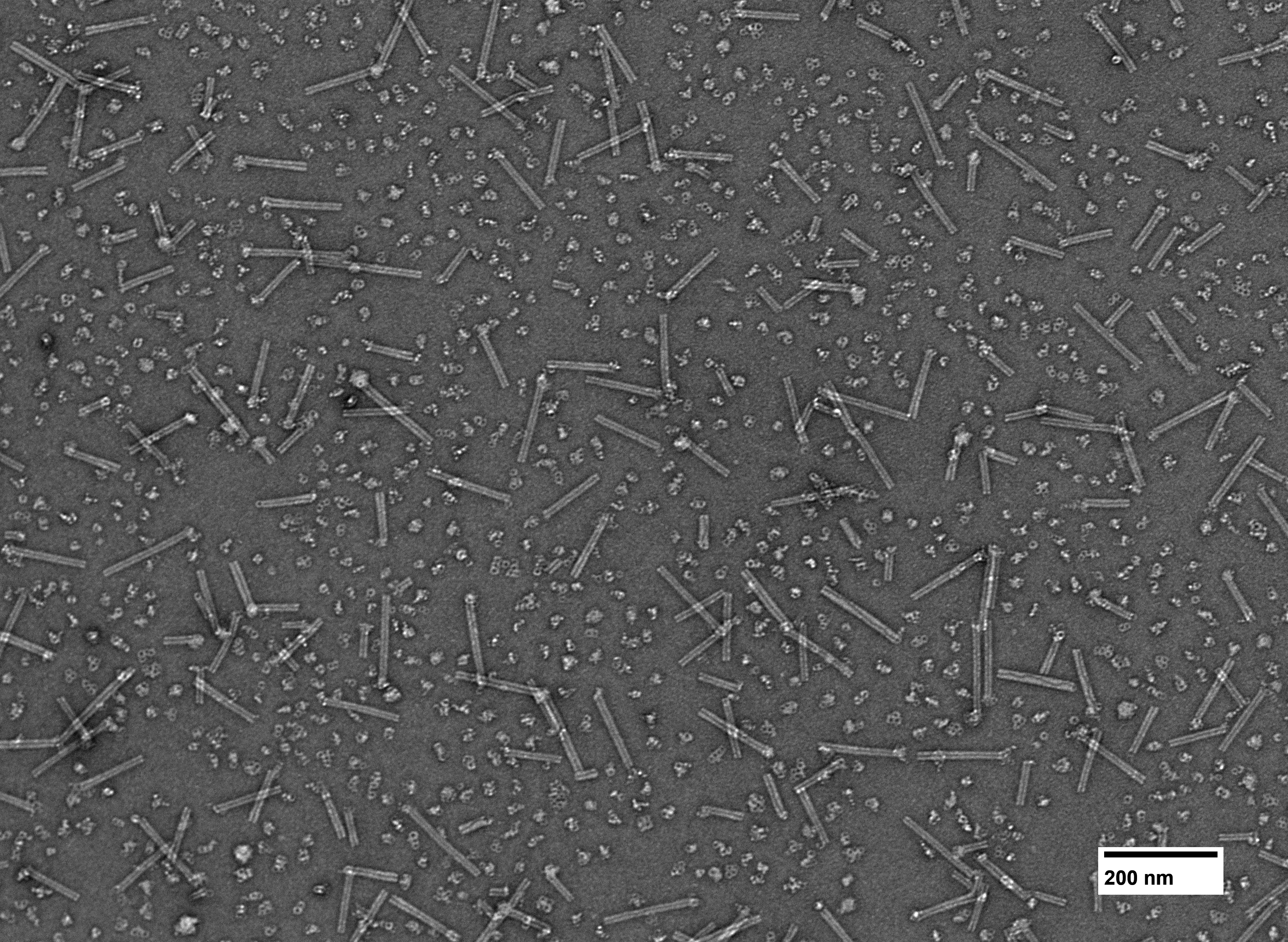

### 20250602_REM_Singleprimes_Conc_Cyl_1_032.tif

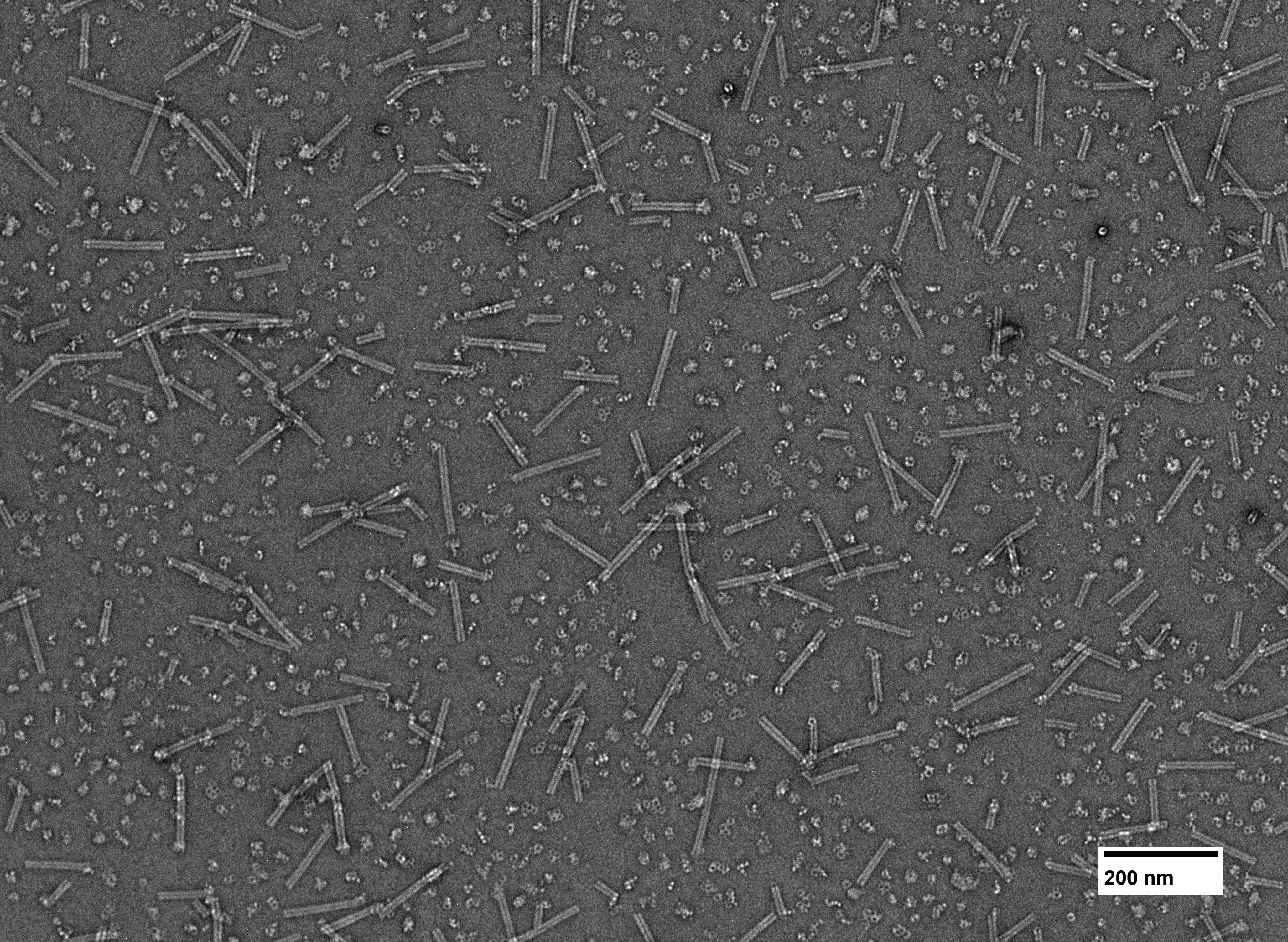

### 20250602_REM_Singleprimes_Conc_Cyl_1_033.tif

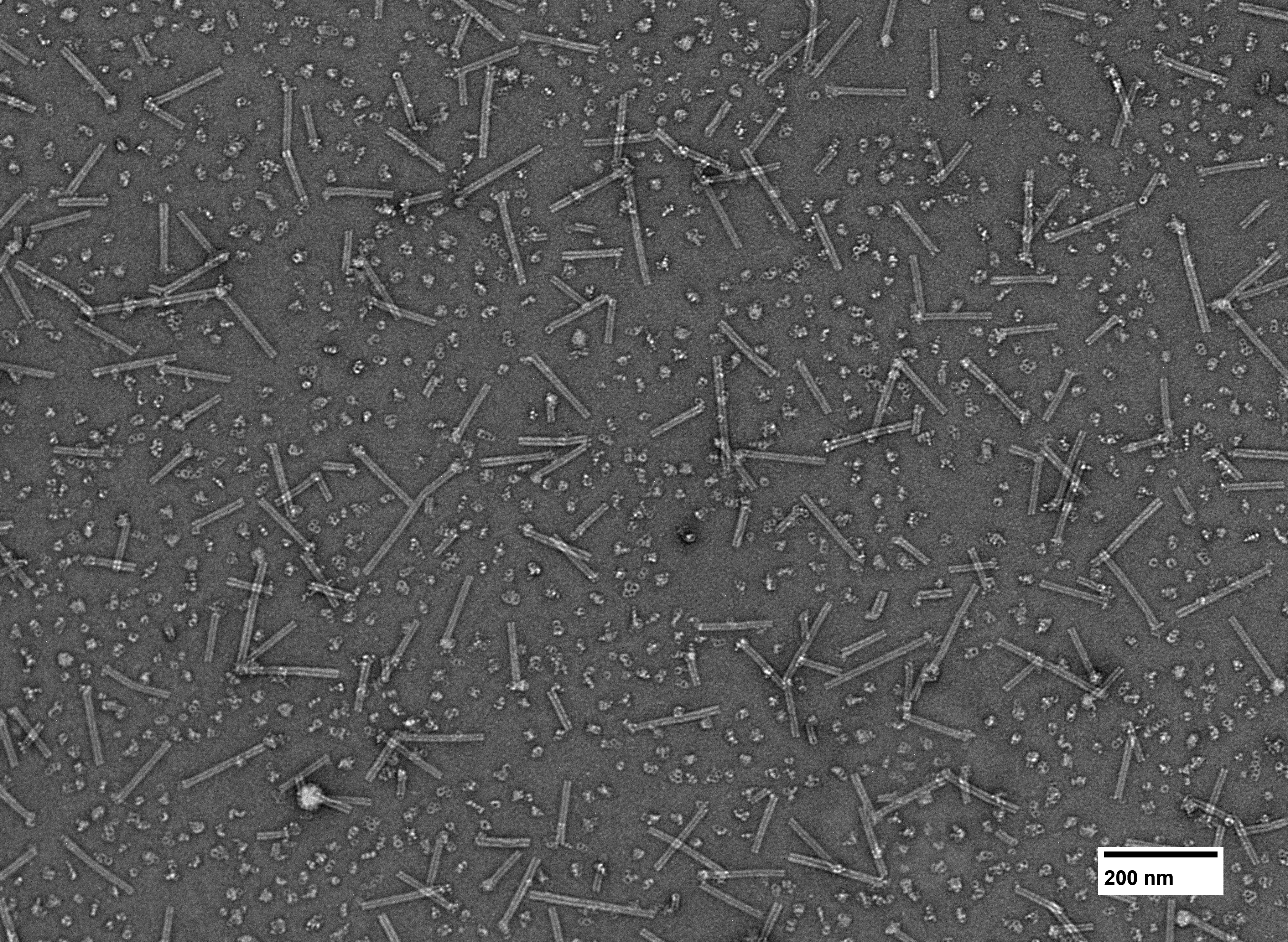

### 20250602_REM_Singleprimes_Conc_Cyl_1_034.tif

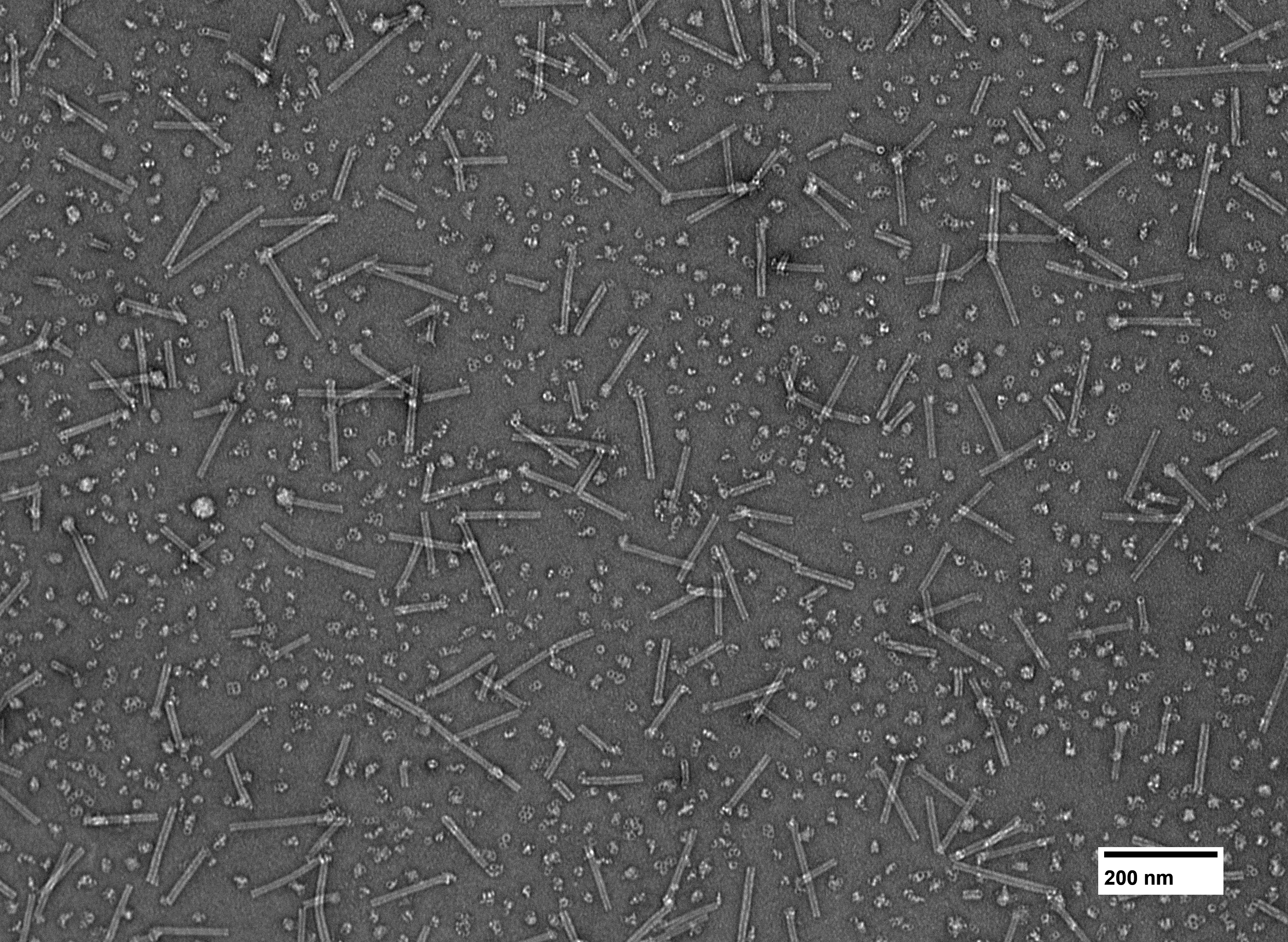

### 20250602_REM_Singleprimes_Conc_Cyl_1_035.tif

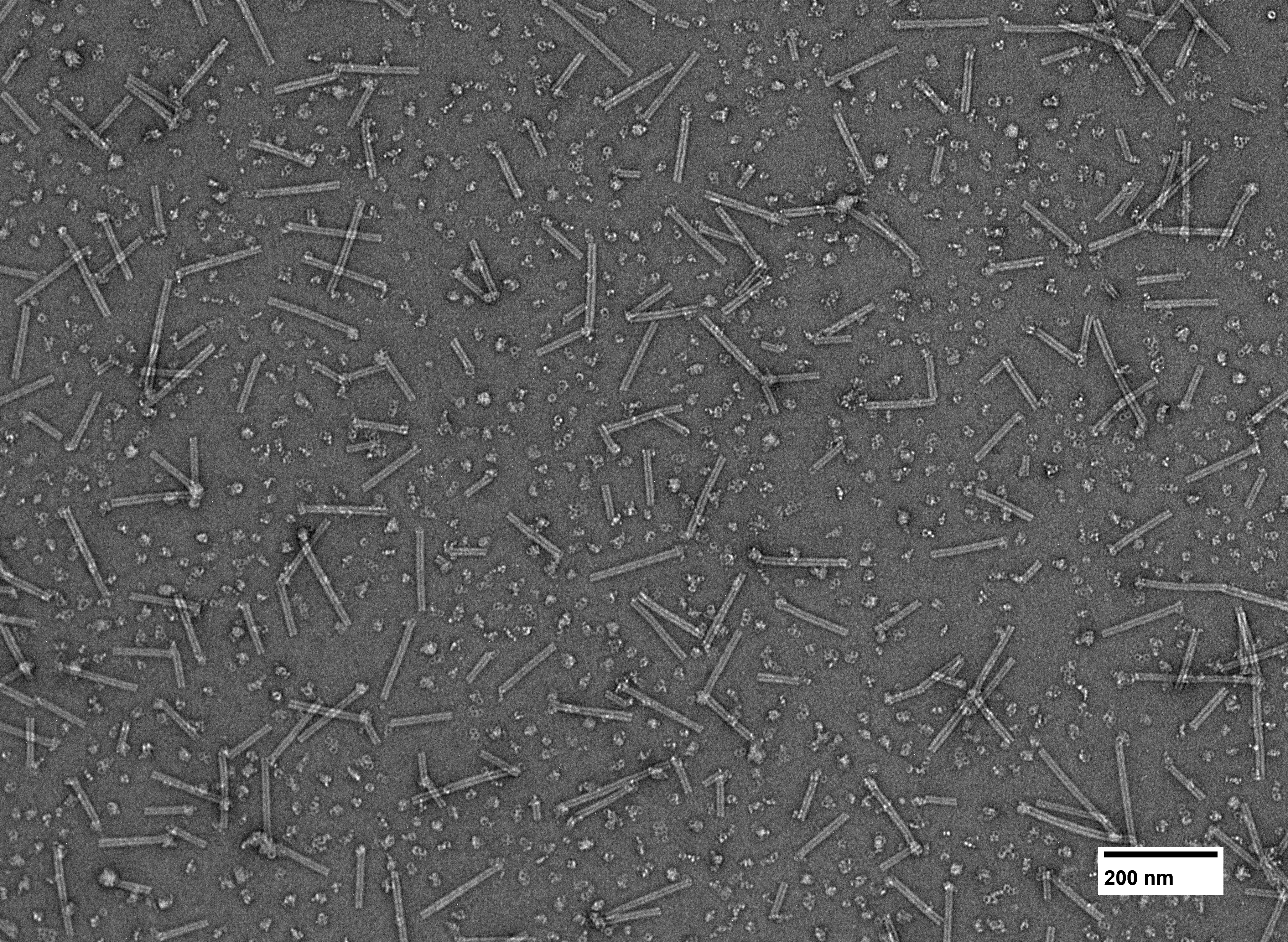

### 20250602_REM_Singleprimes_Conc_Cyl_1_036.tif

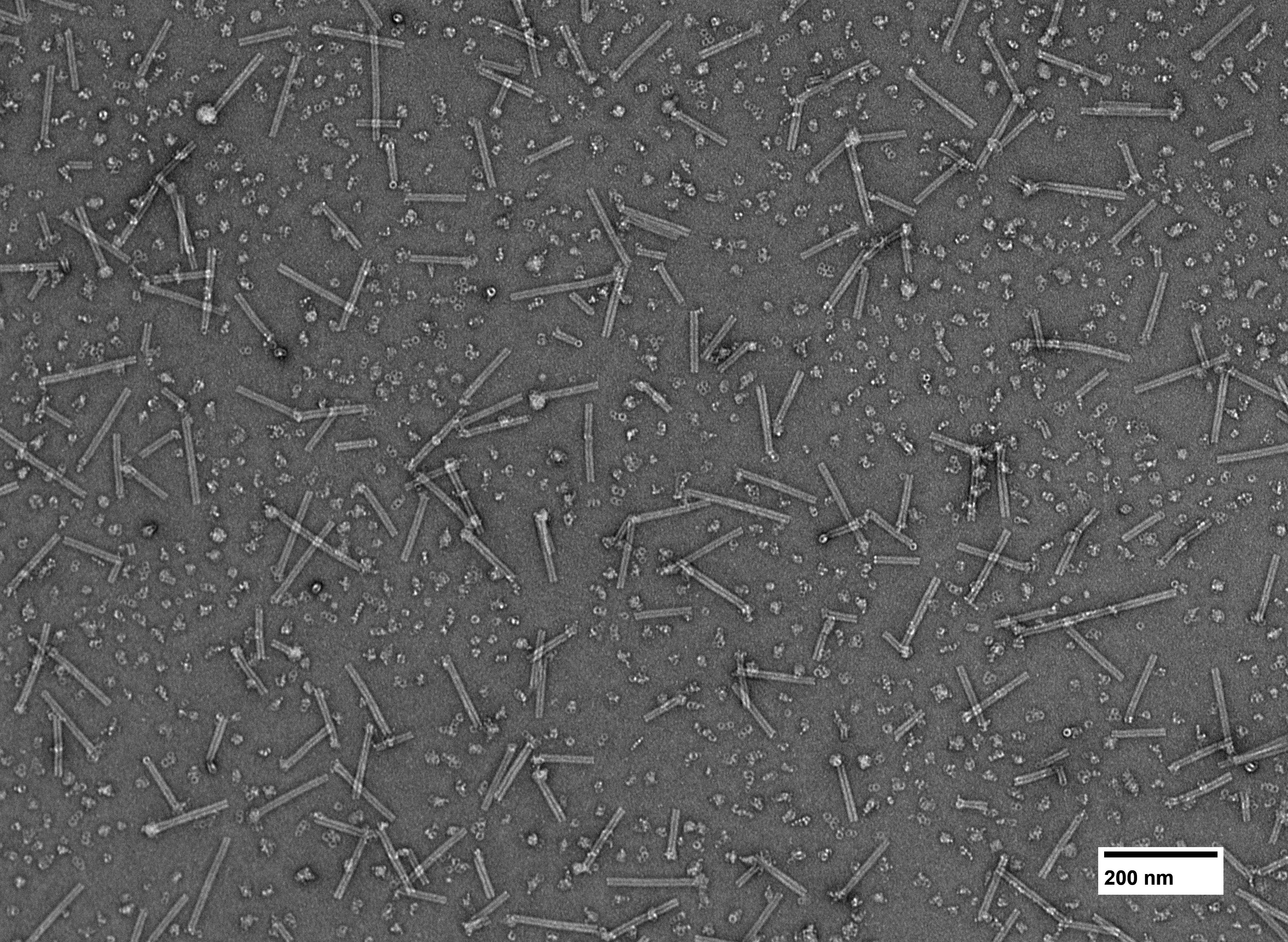

### 20250602_REM_Singleprimes_Conc_Cyl_1_037.tif

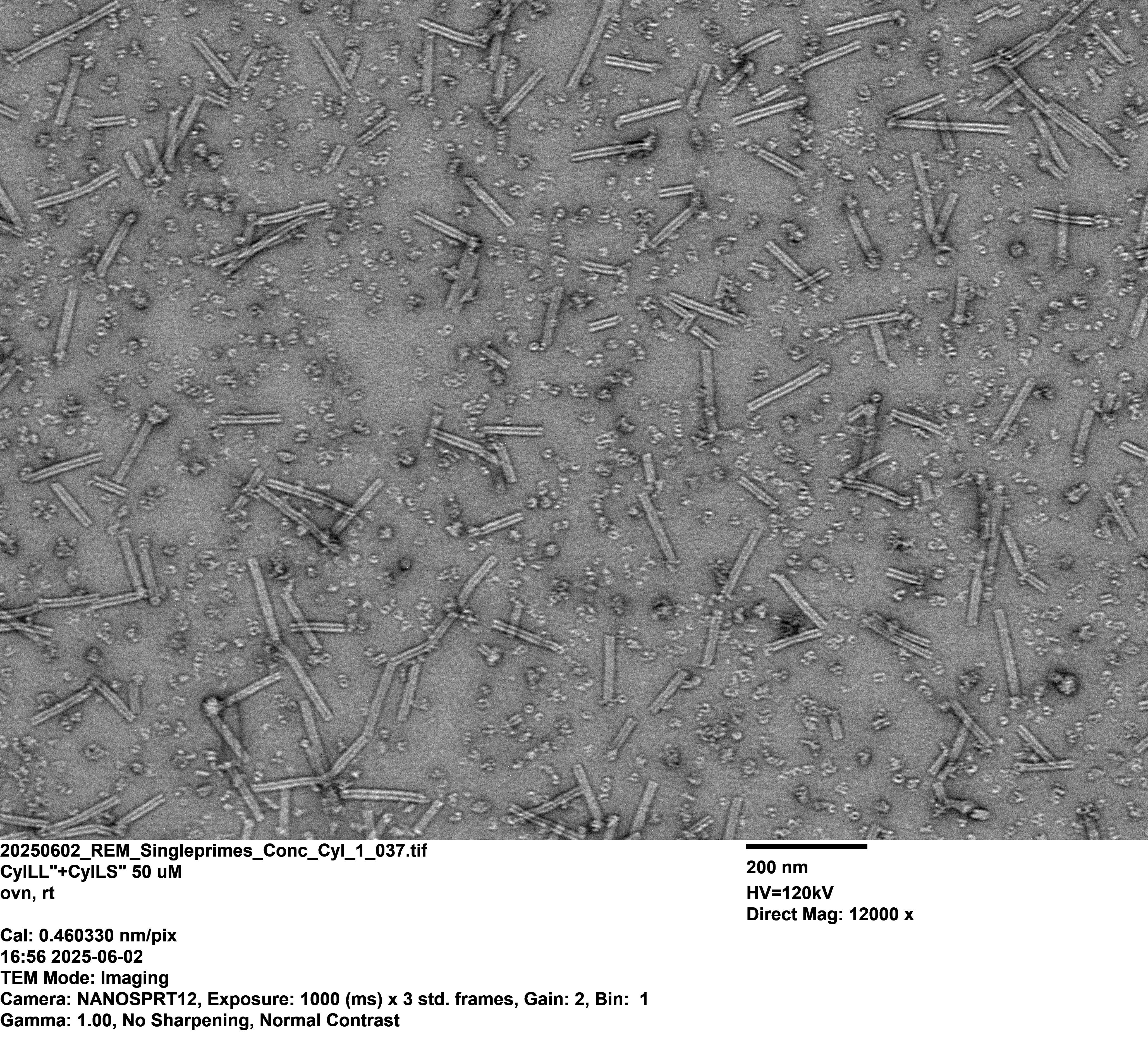

### 20250602_REM_Singleprimes_Conc_Cyl_1_038.tif

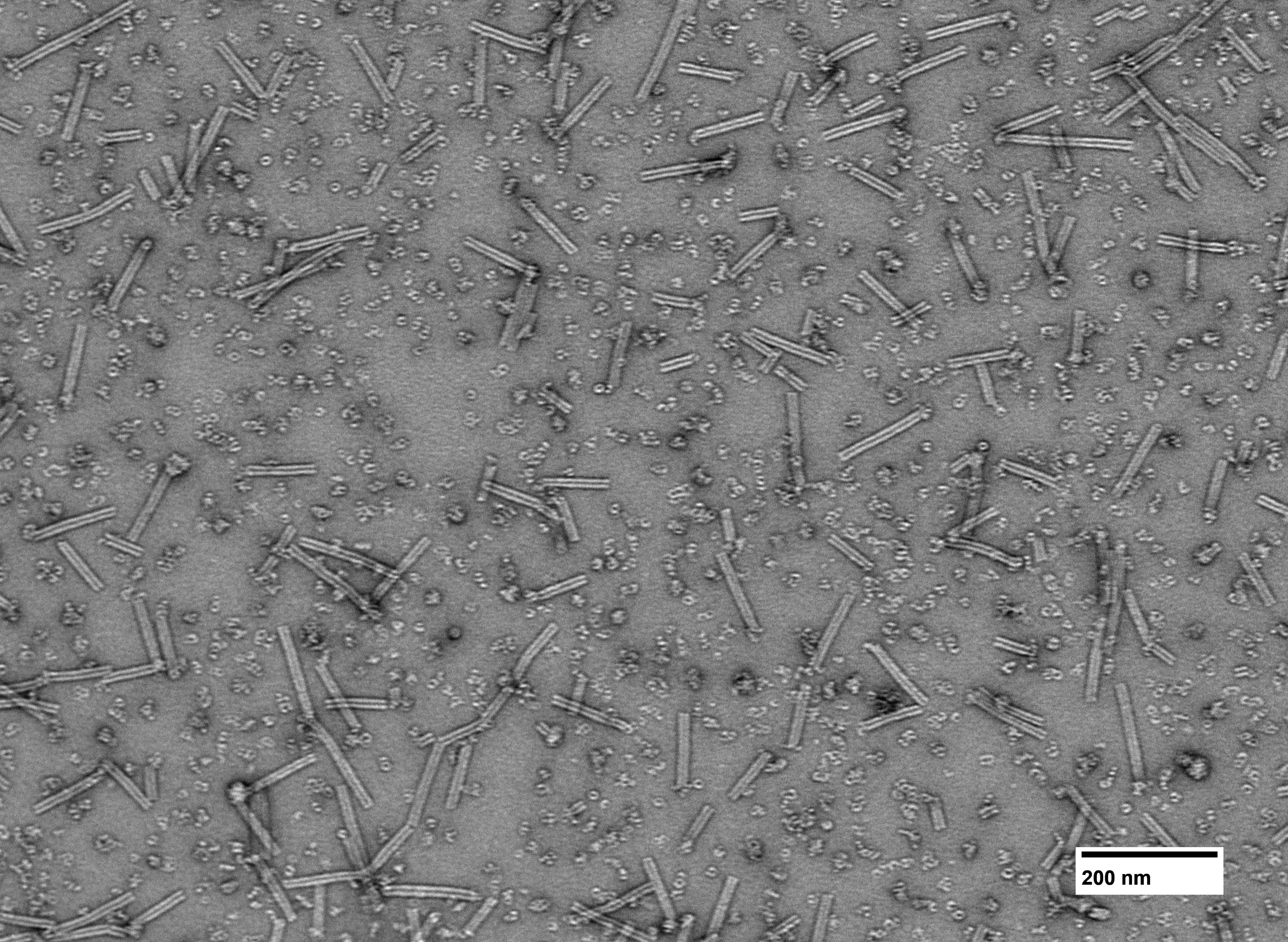

### 20250602_REM_Singleprimes_Conc_Cyl_1_039.tif

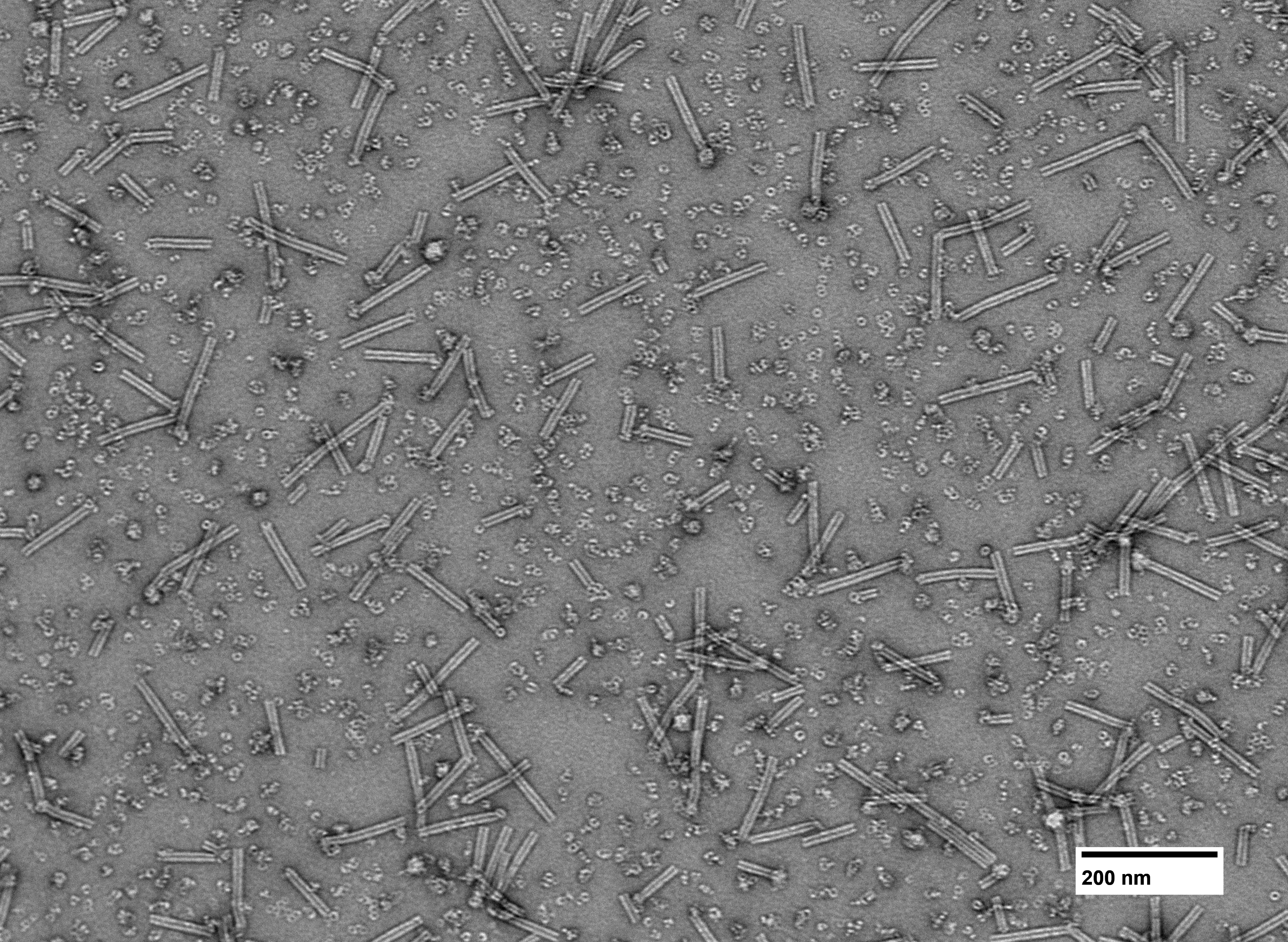

### 20250602_REM_Singleprimes_Conc_Cyl_1_040.tif

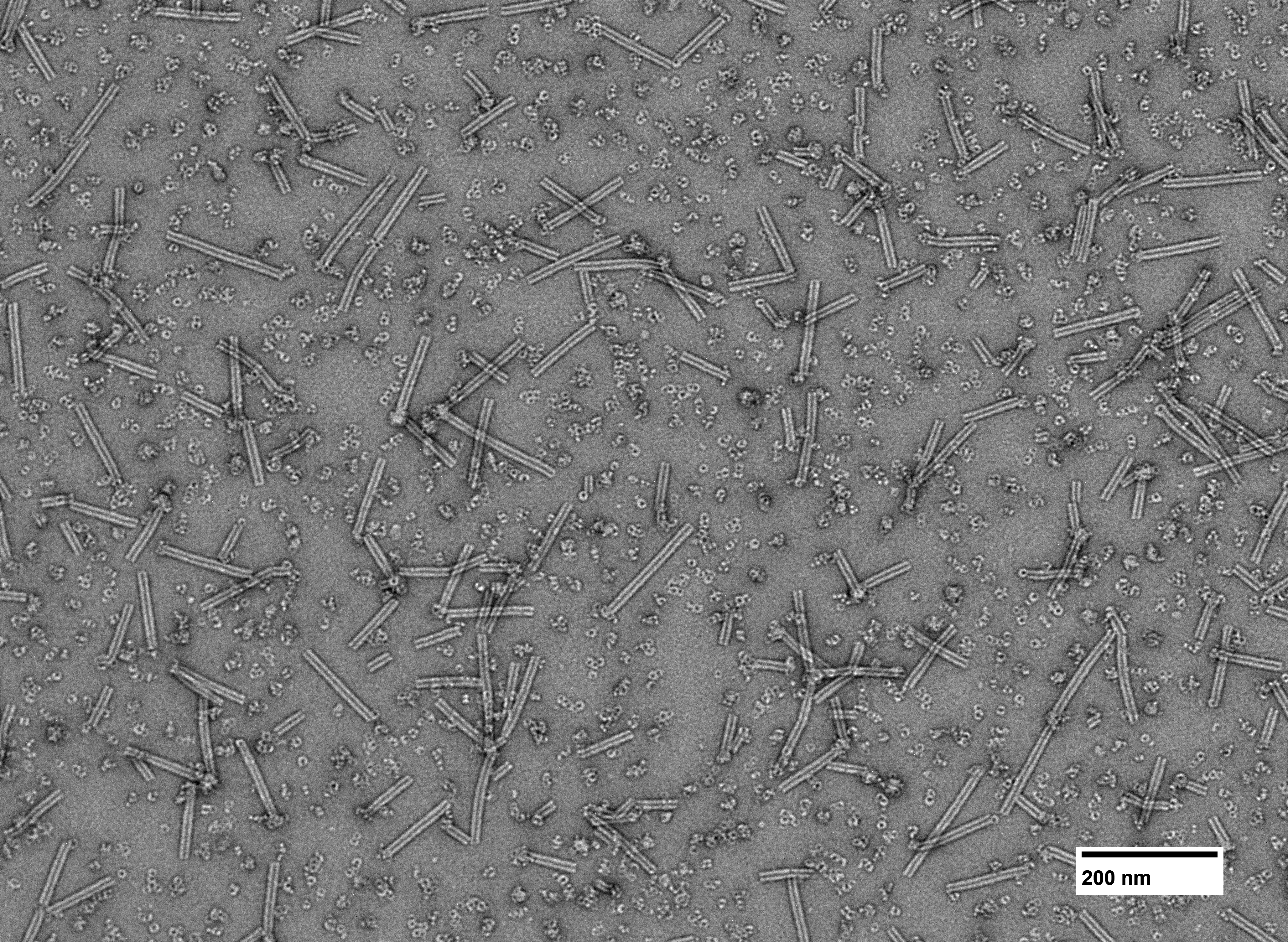

### 20250602_REM_Singleprimes_Conc_Cyl_1_041.tif

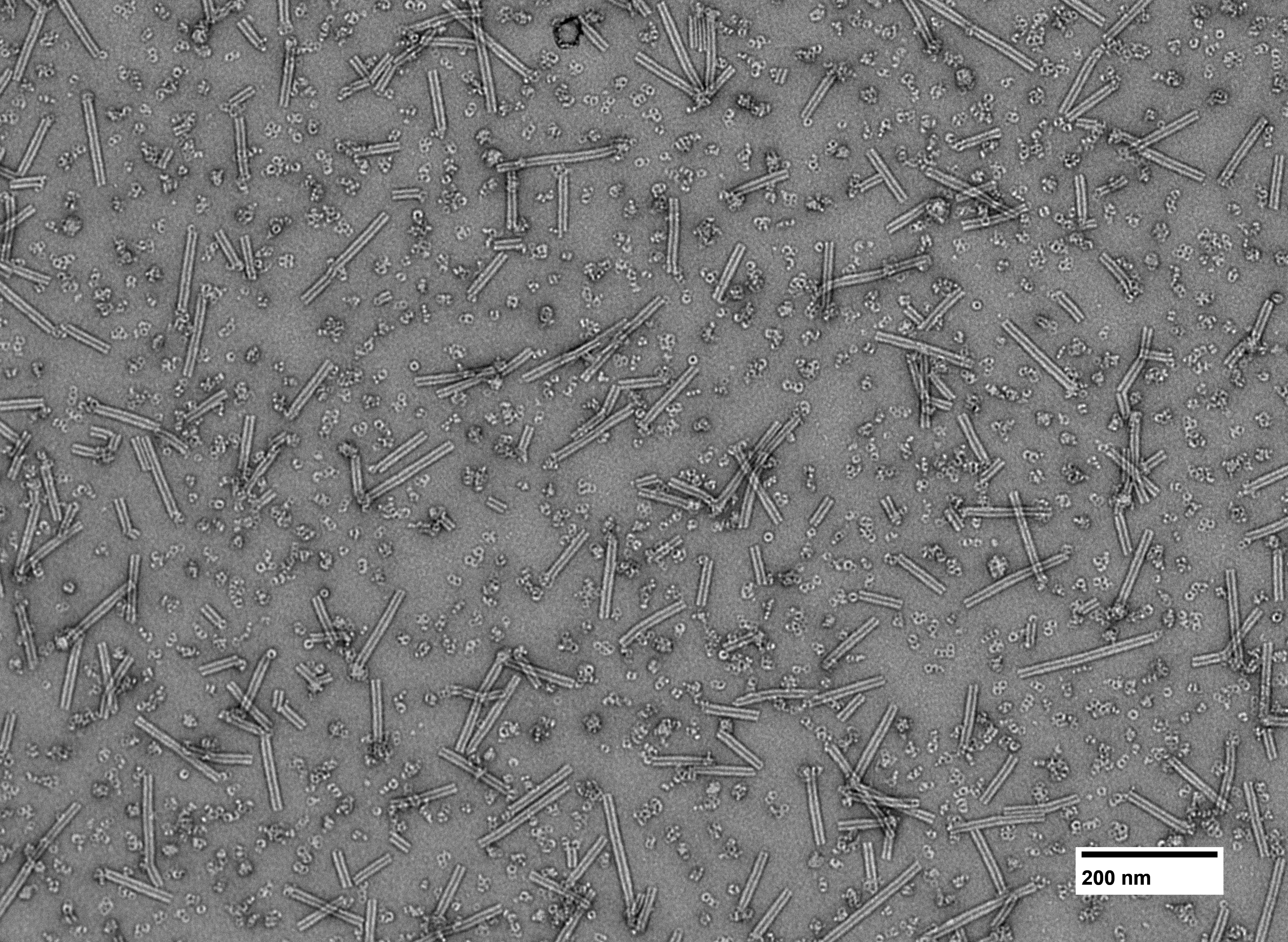

### 20250602_REM_Singleprimes_Conc_Cyl_1_042.tif

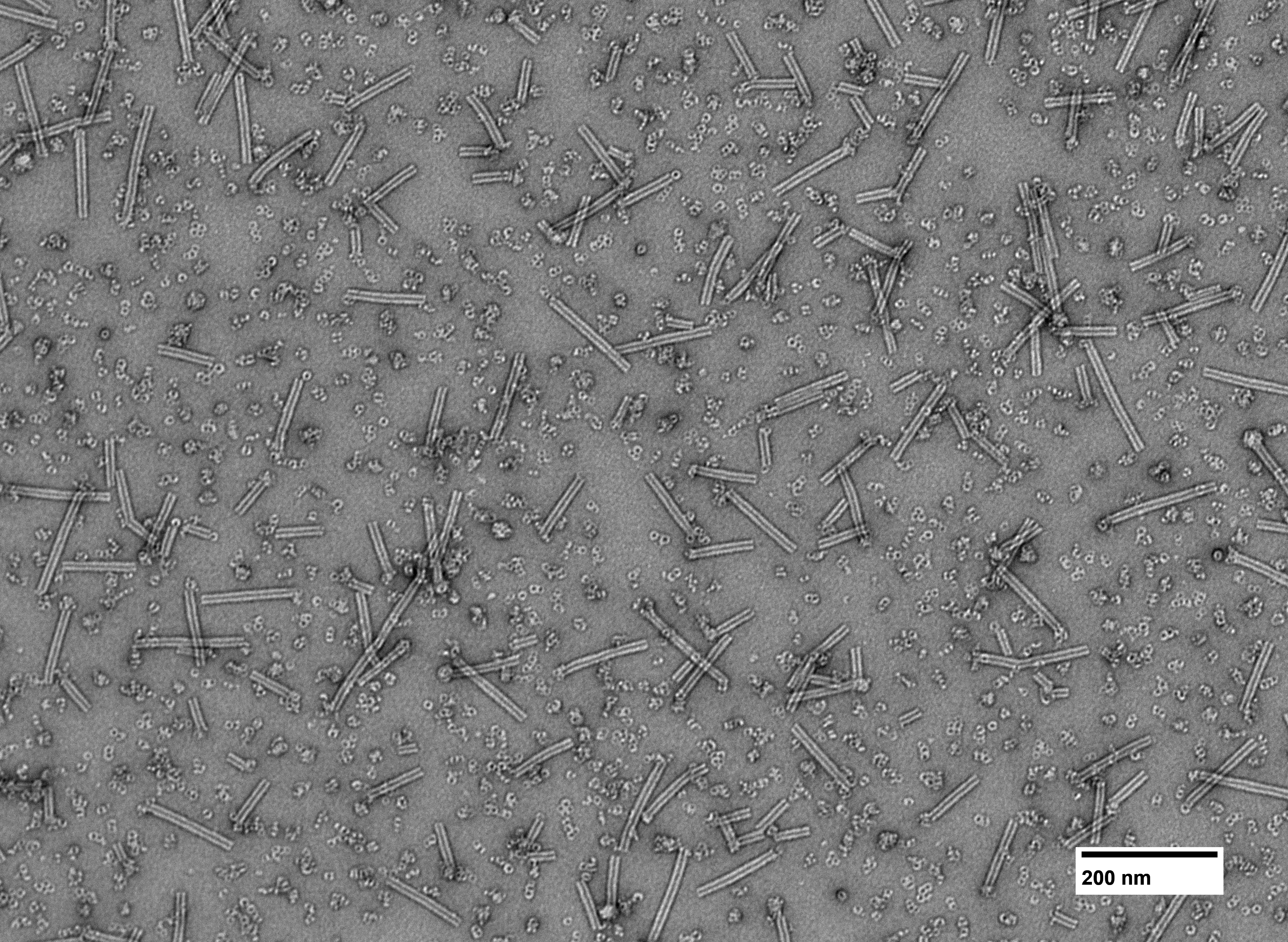

### 20250602_REM_Singleprimes_Conc_Cyl_1_043.tif

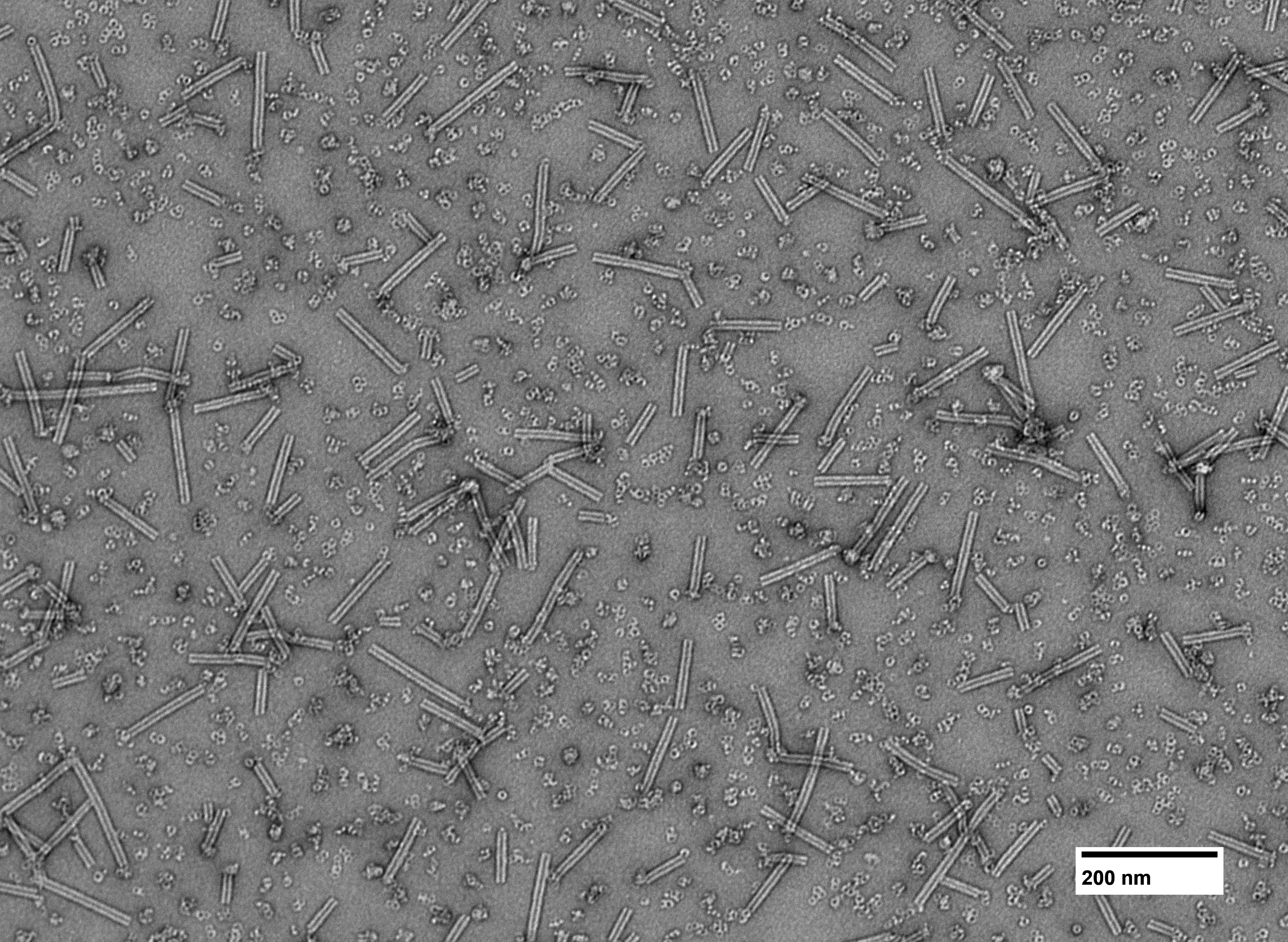

### 20250602_REM_Singleprimes_Conc_Cyl_1_044.tif

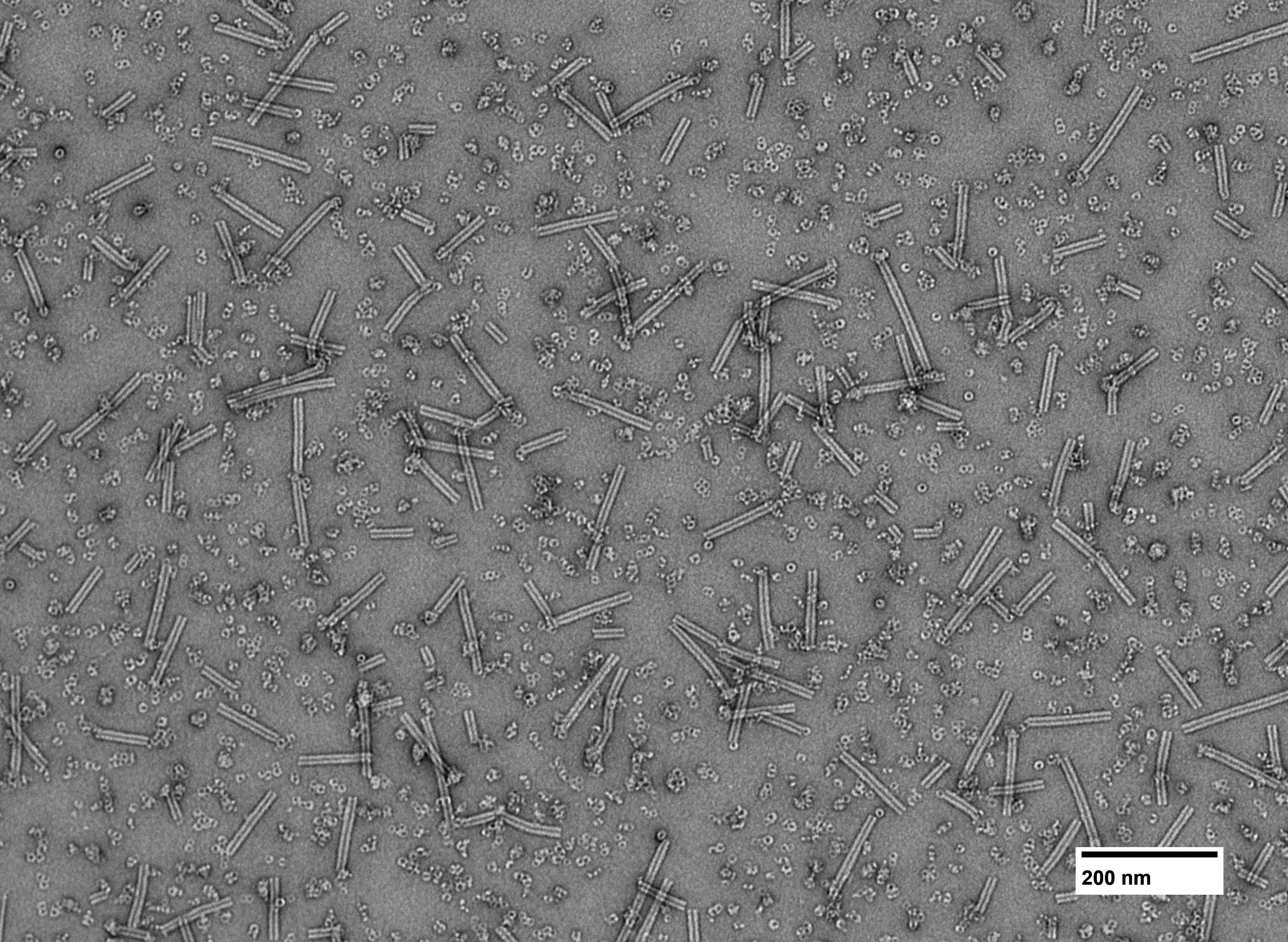

### PXL_20241125_230927301.jpg

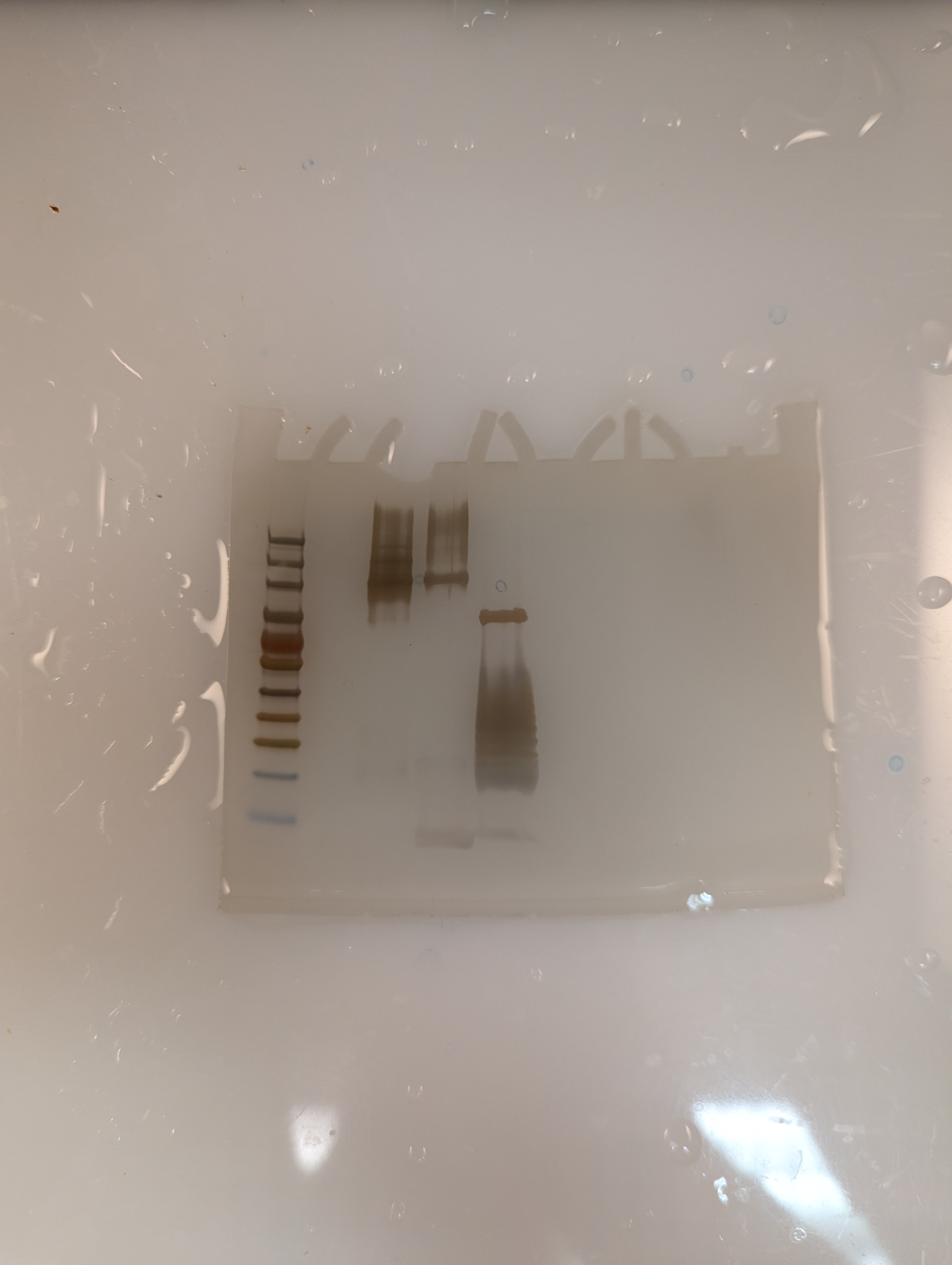
