## Supplemental Materials for "Structure and Mechanism of a Two-component Lanthipeptide Toxin"

**Affiliations:**
[1]Department of Chemistry and Howard Hughes Medical Institute, University of Illinois at Urbana-Champaign, Urbana, IL, USA
[2]Department of Biochemistry, Brandeis University, Waltham, MA, USA
[3]Harvard Cryo-EM Center for Structural Biology, Boston, MA, USA
[4]Department of Chemistry and Biochemistry, College of Arts and Sciences, Florida International University, Miami, FL, USA
[5]Thomas M. Siebel Center for Computer Science, University of Illinois at Urbana-Champaign, Urbana, IL, USA
[6] Theoretical and Computational Biophysics Group, NIH Resource for Macromolecular Modeling and Visualization, Beckman Institute for Advanced Science and Technology, Department of Biochemistry, and Center for Biophysics and Quantitative Biology, University of Illinois Urbana-Champaign, Urbana, IL, USA

**The PDF file includes:**

Materials and Methods
Figs. S1 to S13
Table S1
References 56-91

**Other Supplementary Materials for this manuscript include the following:**
Data sets S1 and S2

**Materials and Methods**

General

All organic solvents and other chemicals were purchased from Sigma Aldrich unless another source is specified. Nisin was isolated as described previously (*56*). Cytolysin was prepared and isolated as described previously (*57*). All buffers were prepared in Milli-Q water and filtered prior to use. Peptides were characterized using Bruker Daltonics UltrafleXtreme MALDI TOF/TOF instrument at the University of Illinois School of Chemical Sciences Mass Spectrometry Facility. Negative stain transmission electron microscopy was carried out in part in the Materials Research Laboratory Central Research Facilities at the University of Illinois.

High-Resolution Tandem Mass Spectrometry

High-performance liquid chromatography coupled to mass spectrometry (LC-MS) utilized an Agilent 1290 LC-MS QToF instrument and a 150 × 2.1 mm Kinetex 2.6 µm C8 column heated to 45 °C. The LC method utilized a flow rate of 0.4 mL/min and two solvents: Solvent B (acetonitrile containing 0.1% formic acid (FA)) and Solvent A (water containing 0.1% FA). The method started with 2 min at 5% Solvent B/95% Solvent A then a linear gradient to 95% Solvent B/5% Solvent A over 6 min then 2 min at 95% Solvent B/5% Solvent A. High-resolution mass spectra were collected in the positive mode. Tandem-MS fragmentation utilized normalized collision energies of 20 and 30.

Preparation and analysis of stocks of cytolysin

Stocks of $CylL_S''$ and $CylL_L''$ were prepared in methanol. Stocks were analyzed by LC-MS using a Kinetex 1.7 µm C8 100 Å, 150 × 2.1 mm LC Column. For concentration determination, an external standard of CylL$_L$″ whose concentration was determined by absorbance at 280 nm was used. The absorbances at 205 nm of the analyte and standard peptide peaks were compared and concentration was determined using the following equation:

$$C_{Anl} = \frac{Abs_{Anl} * n_{Std}}{Abs_{Std} * n_{Anl}} * C_{Std}$$

$C_{Anl}$ and $C_{Std}$ represent the concentration of the analyte and standard peptide, respectively. $Abs_{Anl}$ and $Abs_{Std}$ are absorbance of the analyte and standard peptide at 205 nm, respectively. The number of amide bonds present in analyte and standard peptides were represented by $n_{Anl}$ and $n_{Std}$ , respectively. Peptides were stored separately at –20°C.

<u>Preparation of CylL$_L$″-C5A, CylL$_S$″-C5A, CylL$_S$′ and CylL$_L$′</u>

CylL$_L$″-C5A was prepared following a previously reported approach (*57*). The full length, modified precursor His$_6$-mCylL$_S$″-C5A was prepared by a previously described approach (*57*). After immobilized metal affinity purification and protease treatment, the solution containing crude peptide was acidified to a pH equal to 4 with trifluoroacetic acid (TFA) and incubated at room temperature for 30 min before centrifugation (4,000 ×g for 10 min). The supernatant was removed, and the pellet was extracted with 90% methanol/10% water. The extract was purified following the previously reported HPLC method (*57*).

The full length, modified precursors His$_6$-mCylL$_S$″ and His$_6$-mCylL$_L$″ were prepared by a previously described approach (*57*). CylL$_S$′ and CylL$_L$′ were prepared by treatment of the full-length precursors with the protease LahT$_{150}$ (10 μM), incubation for 16 h at room temperature, acidification to a pH equal to 4 using TFA, and purification by HPLC (*58*). The HPLC method used was described previously for purifying cytolysin analogs (*57*). CylL$_S$′ was stored as a solution in methanol. CylL$_L$′ was stored as solution in 66:33.9:0.1 acetonitrile:water:TFA. Samples of

$CylL_S'$ and $CylL_L'$ were analyzed by LCMS. See Fig. S5 for MS-MS characterization data. Raw data can be found in Data S1.

Preparation of $CylL_L''$-NBD

A solution containing 1 mM $CylL_L''$, 15 mM 4-chloro-7-nitrobenzofurazan, and 20 mM *N*,*N*-diisopropylethylamine in methanol was incubated at 45 °C for 240 min then incubated at room temperature overnight. $CylL_L''$-NBD was purified with a 250 × 4.6 mm Hypersil GOLD 5 µm C8 column attached to an Agilent 1200 analytical HPLC system. The separation method used a flow rate of 1 mL/min, started with 10 min at 5% acetonitrile containing 0.1% TFA/95% water containing 0.1% TFA which was followed by a linear gradient that started from 5% acetonitrile containing 0.1% TFA/95% water containing 0.1% TFA and progressed to 95% acetonitrile containing 0.1% TFA/5% water containing 0.1% TFA over 30 min. $CylL_L''$-NBD eluted after 28 min. Product containing fractions were combined and lyophilized. To prepare stocks of $CylL_L''$-NBD the residue was dissolved in methanol and the concentration was determined from the absorbance in methanol using a molar absorptivity of 22,000 $M^{-1}cm^{-1}$. A sample of purified $CylL_L''$-NBD was analyzed by LC-MS. See Fig. S2 for characterization data.

Generating cytolysin resistant mutants of *Lactococcus lactis*

*Lactococcus lactis* CNRZ 481 (*59*) was grown overnight to an $OD_{600}$ equal to 1.0 in GM17 media and plated on GM17 agar plates supplemented with 2 × MIC (16 nM) cytolysin. The plates were incubated overnight at 30 °C for 3 d and colonies were observed. Single colonies were used to inoculate 5 mL of GM17 media supplemented with 2 × MIC cytolysin and incubated at 30 °C for 24 h. The cultures that grew were plated on GM17 agar supplemented with 4 × MIC cytolysin and incubated at 30 °C for 24 h. Single colonies were picked and used to inoculate 5 mL of GM17 media supplemented with 4 × MIC cytolysin and incubated at 30 °C for 24 h. The cultures that grew were used to inoculate another 5 mL of GM17 media supplemented with 4 × MIC cytolysin and incubated at 30 °C for 24 h. The cultures that grew were used to inoculate 5 mL of GM17 media that did not contain cytolysin and incubated at 30 °C for 24 h. The cultures were then used to inoculate 5 mL of GM17 media supplemented with 4 × MIC cytolysin and incubated at 30 °C for 24 h. The cultures that grew were plated on GM17 agar supplemented with 4 × MIC cytolysin and incubated at 30 °C for 24 h. The resulting colonies were used to prepare glycerol stocks. In total, seven resistant clones were subjected to genome sequencing.

Genomic DNA Extraction and sequencing of WT and resistant clones

Glycerol stocks of the parental *L. lactis* were used to inoculate 5 mL of GM17 media and the cells were incubated at 30 °C for 24 h. Glycerol stocks of the cytolysin-resistant strains were used to inoculate 5 mL of GM17 media supplemented with 4 × MIC cytolysin and incubated at 30 °C for 24 h. The cultures were centrifuged at 5000 ×g for 10 min and genomic DNA was isolated using PureLink™ Genomic DNA Mini Kit (ThermoFisher) according to the manufacturer's protocols. The concentration of DNA was quantified by Qubit dsDNA BR Assay (ThermoFisher) and submitted to the Roy J. Carver Biotechnology High-Throughput Sequencing and Genotyping Unit (UIUC) for library preparation and 2×150 bp paired-reads on an Illumina NovaSeq 6000. Bioinformatics analysis was performed by Gloria Rendon (Roy J. Carver Biotechnology HCPBio, UIUC). NCBI RefSeq assembly GCF_000006865.1 was used as reference.

RNAseq analysis of WT and resistant mutants of *L. lactis*

A cytolysin-resistant strain of *L. lactis* was treated with vehicle or a 1:1 mixture of $CylL_S''$:$CylL_L''$ (320 nM final concentration) and grown to an $OD_{600}$ equal to 0.5. The parental (WT) strain was treated with vehicle. Once density was achieved, the cells were collected by centrifugation (6,000 ×g for 10 min) then total RNA was extracted using a Monarch® Total RNA Miniprep Kit. Sequencing and RNA-seq analysis were performed at the Carver Biotechnology Center at the University of Illinois. For a complete table of transcriptional changes, see Data S1.

Time-kill assay

*Lactococcus lactis subsp. cremoris* NZ 9000 was streaked on GM17 agar and incubated at 30 °C overnight (*60*). Five colonies were selected from the center of the plate and used to inoculate sterile GM17 media which was incubated overnight at 30 °C with shaking. The overnight culture was diluted in GM17 to an $OD_{600}$ equal to 1 then diluted 100-fold in GM17 media. Aliquots of this diluted sample of bacteria were treated with $CylL_S''$, $CylL_L''$ or a stoichiometric mixture of $CylL_L''$:$CylL_S''$. The total peptide concentration in each aliquot was 32 nM (8 × MIC). After treatment, the aliquots were incubated at 30 °C with shaking. Samples were harvested from each aliquot after 30, 90, 150, and 210 min. These samples were diluted in phosphate-buffered saline and streaked on GM17 agar. Agar plates were incubated overnight at 30 °C. Colonies were counted and used to calculate the density of live cells (CFU/mL) at each time point.

Cell permeabilization assay with SYTOX green

An overnight culture of *Lactococcus lactis subsp. cremoris* NZ 9000 was diluted 100-fold in fresh media then incubated with shaking at 30 °C until the $OD_{600}$ reached 0.4. The cells were then collected by centrifugation (6000 × g for 10 min) and washed with M9 minimal media

(ThermoFisher) containing $MgCl_2$ (2 mM), $CaCl_2$ (0.1 mM) and D-glucose (5 mM) then resuspended in the same media to an $OD_{600}$ equal to 1.0. SYTOX green (Invitrogen™, ThermoFisher) was added to achieve a concentration of 5 μM and the suspension was transferred to a 96-well plate black polystyrene plate (90 μL per well). The fluorescence of SYTOX green was monitored for 10 min in a plate reader pre-heated to 30 °C (Biotek Synergy H4) using an excitation wavelength equal to 488 nm and an emission wavelength equal to 523 nm. After this incubation period, peptide or vehicle diluted in M9 media (10 μL) was added and the emission was monitored for 60 min. $CylL_L''$ and $CylL_S''$ were present in 1:1 ratio at a total peptide concentration of 16 nM (2 × MIC). The concentration of nisin was 600 nM (2 × MIC). The experiment was conducted on three different cultures of *L. lactis subsp. cremoris* ($n$ = 3). Error bars represent the standard of deviation between three trials. Data from these experiments can be found in Data S1.

Assaying the intracellular pH of *L. lactis*

This procedure is based on a literature report (*61*). Overnight cultures of *L. lactis subsp. cremoris* were diluted 40-fold in GM17 media, then incubated at 30 °C with shaking until mid-log phase. The cells were collected by centrifugation (3,000 g for 10 min at 4 °C), then resuspended in Buffer A (50 mM HEPES, 20 mM D-glucose, 1 mM $MgSO_4$ at pH = 7) to an $OD_{600}$ of 0.5. To this suspension of bacteria was added a mixture of 5- and 6-carboxyfluorescein diacetate succinimidyl ester (5(6)-CFDA SE) to a concentration of 3 μM. The cells were incubated with the dye for 30 min at 30 °C with shaking. The cells were collected by centrifugation as before and washed with Buffer A then suspended in Buffer B (50 mM HEPES, 20 mM glucose, 1 mM $MgSO_4$ at pH = 5) to an $OD_{600}$ equal to 0.05. The cells were immediately dispensed into a 96-well plate black polystyrene plate (90 μL per well) and transferred to a plate reader (Biotek Synergy H4). Using an excitation wavelength equal to 490 nm and an emission wavelength of 525 nm, the fluorescence emission intensity was measured for 2 min, then cytolysin (16 nM), nisin (600 nM) or CCCP (20 μg/mL) was added, and emission intensity measurement was continued. After 10 min, CCCP (20 μg/mL) was added and emission intensity measurement was continued for 5 min. This experiment was done in triplicate (*n* = 3). Data from these experiments can be found in Data S1.

Toxicity studies with HepG2 cells

HepG2 cells were revived and maintained following the procedures outlined by the American Type Culture Collection (ATCC). Cells were grown in ATCC-formulated Eagle's Minimum Essential Medium (EMEM) containing 10% fetal bovine serum (Gibco™, ThermoFisher) that was not heat inactivated. Following revival from cryopreservation, the cells were passaged twice then transferred to a 96-well plate at concentration of $10^4$ cells per well. The cells were incubated at 37 °C for 24 h in an atmosphere containing 5% $CO_2$ before the growth media was replaced with serum-free EMEM. The cells were incubated overnight then used for toxicity assays.

Stocks of $CylL_S''$ and $CylL_L''$ were diluted separately in calcium and magnesium free Dulbecco's phosphate buffered saline (DPBS) to ten times the assay concentration, then added to cell-containing wells. The stimulated cells were incubated for 1 h or 3 h at 37 °C, then the lactate dehydrogenase (LDH) activity of the supernatant of each well was determined using a CyQUANT™ LDH Cytotoxicity Assay (ThermoFisher). These experiments were repeated at least 3 times (*n* = 3). Data from these experiments can be found in Data S1.

Confocal microscopy with fluorescently labelled cytolysin

HepG2 cells were seeded into a Nunc™ 96-Well Optical-Bottom Microplate (ThermoFisher) at $10^4$ cells per well and incubated for 24 h at 37 °C in an atmosphere containing 5% $CO_2$. Serum-containing media was removed, and the cells were washed with phosphate-buffered saline (PBS) then replaced with serum-free EMEM containing $CylL_S''$ and $CylL_L''$-NBD (10 µM total peptide concentration). The plate was incubated for 45 min at 37 °C in an atmosphere containing 5% $CO_2$. The media was removed and the cells were washed once with PBS (no Ca/Mg) then fixed with 4% paraformaldehyde. After 15 min of treatment at room temperature, the paraformaldehyde was removed and the cells were washed three times with PBS (no Ca/Mg). A staining solution was prepared by diluting commercial CellBrite Orange (biotium) solution 200-fold in PBS (no Ca/Mg) and Hoechst 33342 (Sigma) to a final concentration of 10 µg/mL. The staining solution was added to the fixed cells. After 15 min at room temperature, the staining solution was removed, and the cells were washed with PBS (no Ca/Mg) three times then imaged under PBS (no Ca/Mg). Imaging was done within 1 h of sample preparation. A Zeiss LSM900 microscope with 63× magnification and immersion oil (Immersol 518F) were used. Images were analyzed using ZEISS ZEN software v3.9.

<u>Pyranine liposome permeabilization assay</u>

Pyranine-encapsulated liposomes were prepared as described elsewhere (*62*). 1,2-Dioleoyl-sn-glycero-3-phosphocholine (DOPC), 1,2-dipalmitoyl-sn-glycero-3-phosphocholine (DPPC), 1,2-dioleoyl-sn-glycero-3-phospho-L-serine (DOPS), 1,2-dioleoyl-sn-glycero-3-phosphoethanolamine (DOPE), 1,2-dioleoyl-sn-glycero-3-phospho-(1'-rac-glycerol) (DOPG), sphingomyelin (egg, chicken) and cholesterol (Chol) were purchased from Avanti. Liposomes composed of 100% DOPC, 7:3 DOPC:DOPG, 7:3 DOPE:DOPG, or DPPC:DOPE:DOPS:

sphingomyelin:Chol 20:20:10:15:35 were prepared using this approach. Briefly, thin films of lipid residue were resuspended in low pH buffer (5 mM Tricine, 5 mM MES, 5 mM NaCl, 1 mM pyranine pH = 6.0), freeze-thawed three times then extruded through a 200 nm polycarbonate filter 21 times. Unencapsulated pyranine was removed by a Sephadex G50 column equilibrated with buffer (5 mM Tricine, 5 mM MES, 5 mM NaCl, pH = 6.0). The prepared liposomes were diluted in high pH buffer (5 mM Tricine, 5 mM MES, 5 mM NaCl, pH = 8.0) in a 96-well black polystyrene plate to a total lipid concentration of 50 µM. The plate was transferred to a plate reader (Biotek Synergy H4) and the pyranine emission intensity was monitored for 8 min. Cytolysin was added to a total concentration of 200 nM and the pyranine emission intensity was monitored for 30 min. Triton X-100 was added to a concentration of 0.1% and the emission intensity was monitored for 5 min. Fraction of permeabilization was determined by min-max normalization of emission intensity data over the course of the entire experiment. Experiments were performed in triplicate (n = 3). Standard of deviation is represented with error bars. Data from these experiments can be found in Data S1.

Broth dilution assay for determining minimum inhibitory concentrations (MICs)

MICs were determined via a broth dilution assay (*63*). *L. lactis subsp. cremoris* was grown at 30 °C in GM17. Overnight cultures were sub-cultured until they reached the mid-log phase of growth, then they were diluted in growth media and combined with serial dilutions of antibiotic in growth media. The final concentration of bacteria was $5 \times 10^5$ CFU/mL. After 16–18 h of incubation, the MIC was determined visually and by $OD_{600}$.

Analysis of cytolysin peptides using SDS-PAGE

Cytolysin peptides were diluted in PBS (pH = 7.4) to a concentration of 100 μM and incubated at room temperature overnight. The resulting solution was combined with 4 × Laemmli sample buffer containing 8% SDS (the final concentration of SDS was 2%), incubated at 37 °C for 30 min, then loaded onto a 4–20% Mini-PROTEAN® TGX™ (BioRad). See Data S2 for unaltered gel images.

Analysis of cytolysin oligomers by high-resolution mass spectrometry

The high-resolution mass spectrometry analyses were performed on a Bruker Maxis Impact II ToF MS instrument (Bruker Daltonics, Inc., Billerica, MA) equipped with an nESI source in positive mode. nESI tip emitters (O.D. = 1.0 mm and I.D. = 0.70 mm) were pulled in-house from quartz capillaries using a Sutter Instruments Co. P2000 laser puller (Sutter Instruments, Novato, CA). A 1:1 $CylL_L''$:$CylL_S''$ ratio solution (total peptide concentration 10 μM) was loaded in a pulled tip capillary housed in a mounted custom-built XYZ stage in front of the MS inlet, where the nESI emitters contained a tungsten wire biased at ~900 V relative to the MS inlet. Ions were softly transferred to avoid potential oligomer disruptions during the oligomerization process between $CylL_L''$ and $CylL_S''$ in the gas-phase.

Negative stain transmission electron microscopy of cytolysin

To image cytolysin, stocks of $CylL_L''$ (620 μM) and $CylL_S''$ (620 μM) in methanol were diluted into MQ $H_2O$ to a total peptide concentration of 50 μM. After incubation at room temperature for 13 h, a 5 μL aliquot was transferred to a glow discharged Carbon Film 200 Mesh, Cu grid and incubated for 1 min before blotting and applying 2% uranyl acetate stain. The staining was allowed to proceed for 15 s before blotting and airdrying. The dry grid was imaged using a JEOL 1400 TEM with an accelerating voltage of 120 kV. See Data S2 for original micrographs.

Negative stain transmission electron microscopy of cytolysin on cells

The procedure for imaging bacteria was based on a previously described method (*64*). Briefly, an overnight culture of *L. lactis subsp. cremoris* NZ9000 was collected by centrifugation (6,000 × g for 10 min), washed with Tris-Glu (10 mM Tris HCl, 5 mM D-glucose, pH 6.0) then resuspended in Tris-Glu (~$10^9$ CFU/mL), and treated with cytolysin (3 μM) or vehicle for 1 min. A glow discharged Carbon Film 200 Mesh, Cu grid previously washed with TBS (150 mM NaCl, 50 mM Tris-HCl, pH 7.4) was transferred to a drop of resuspended bacteria that had been treated with cytolysin. The grid was kept in contact with the bacteria containing drop for 30 s. The grid was transferred to a drop of water for 30 s, then another drop of water for 30 s, followed by two drops of 2% uranyl acetate for 30 s each. Finally, the grid was transferred to a drop of water for 30 s then blotted and allowed to air dry. The dry grid was imaged using a JEOL 1400 TEM with an accelerating voltage of 120 kV. See Data S2 for original micrographs.

Preparation of cytolysin helical oligomers for single particle cryo-EM

Methanol-dissolved stocks of $CylL_L''$ (725 μM) and $CylL_S''$ (1000 μM) were rapidly mixed in milli-Q water to a final concentration of 50 μM in a final volume of 400 μL. The suspension was incubated overnight at room temperature, and the oligomers were then pelleted by centrifugation at 21,000 ×g for 1 h to remove methanol. The supernatant was carefully removed by aspiration and the resulting pellet resuspended in 200 μL of fresh milli-Q water. A 3 μL aliquot of the resulting suspension was vitrified on grids using a Mark IV Vitrobot (ThermoFisher). Grids were glow discharged using an easiGlowTM (Pelco) 30 min prior to sample vitrification. Grid type and blotting time was optimized for the sample using a double-sided blot with a constant force of 0, in a 100% relative humidity chamber at 4 °C, and a 5 s wait prior to plunging in liquid ethane before storing in liquid nitrogen. The condition that yielded the best distribution of cytolysin oligomers was found to result from using QuantifoilTM R2/1 400 mesh copper grids with an additional 2 nm layer of carbon thin film and a blotting time of 5 s.

Cryo-EM grids were screened on a Tundra Cryo TEM (ThermoFisher) and data was collected using a Talos Arctica (ThermoFisher) operating at 200 kV and a Titan Krios (ThermoFisher) microscope operating at 300 kV. The Tundra was equipped with a Falcon-C direct detector, the Talos Arctica was equipped with a K3 direct electron detector (Gatan), and the Titan Krios was equipped with a Falcon-4 direct electron detector and a Selectris energy filter. SerialEM software version 4.2 was used for collections with the Talos Arctica and EPU software version 3.8.1 was used for collection with the Titan Krios. Final data collection on grids of cytolysin helical oligomers was performed using a Titan Krios microscope and a total of 12,689 movies were acquired at a pixel size of 0.74 Å, a total dose of 54.8 e− /$Å^2$, dose per frame of 0.62 e− /$Å^2$ at a defocus range of −0.7 to −1.8 µm.

Detergent extraction of cytolysin oligomers from liposomes

Methanol-dissolved stocks of $CylL_L''$ (725 µM) and $CylL_S''$ (1000 µM) were rapidly mixed with DOPC liposomes to a final concentration of 50 µM in a buffer of 50 mM HEPES-KOH (pH 7.5) and 150 mM NaCl to a final volume of 5 mL. The reaction was incubated overnight at room temperature and then divided into 100 µL aliquots, which were centrifuged at 21,000 ×g for 1 h. Supernatants were removed carefully by a pipette, saved, and each resulting pellet was resuspended in a solution of 50 mM HEPES-KOH (pH 7.5) and 150 mM NaCl with a unique detergent from the Detergent ScreenTM (Hampton) at 1× concentration. The resulting suspensions were incubated at 37 °C for 1 h with rotation and then centrifuged at 21,000 ×g for 1 h. The supernatant containing putative solubilized cytolysin oligomers were carefully removed and the remaining pellet resuspended in 100 µL 50 mM HEPES-KOH (pH 7.5) and 150 mM NaCl. Samples were analyzed by 4–20% SDS-PAGE (BioRad) to determine which detergents were able to solubilize cytolysin oligomers.

Cryo-EM Image Processing and Model building

Movie frames were imported into cryoSPARC for patch-based motion correction and CTF estimation, blob picking, 2D classification, template-based particle picking and filament tracing. Helical symmetry parameters were determined from multiple complementary sources of information. Occasional top-views of tube segments enabled rough estimation of approximately 17–26 subunits per turn. 2D class averages indicated that features repeated with near-translational periodicity along the helical axis across multiple rungs, consistent with a near-integer number of subunits per turn. The helical pitch was estimated directly from the 2D class averages and corroborated by a summed power spectrum, computed using a previously described custom script that sums the power spectra of all particles after in-plane rotational alignment (*65*).

Helical symmetry parameters were systematically screened by running independent helical refinements across a range of subunit counts per turn, testing values of 19, 20, 21, 22, 23, 24, and 25, each paired with a pitch of 22.5 Å. In each refinement, automatic symmetry search was enabled around the input values. Convergence on the correct solution was confirmed by the emergence of well-resolved secondary structure features and side chains in the reconstruction. The final refined parameters were a twist of –15.1°, corresponding to 23.9 subunits per turn, and a rise of 0.974 Å per subunit, corresponding to a pitch of 23.3 Å. Global and local CTF refinement and reference-based motion correction were subsequently applied (*66*).

An initial model of the helical assembly was generated using ModelAngelo (*67*), followed by manual model rebuilding in ISOLDE (*68*). Non-canonical features were added to the structure in Coot (*69*) and the model was iteratively built and refined using Coot and Phenix (*70*). The protomer model was validated using MolProbity in Phenix (*71*) (Table S1). To construct the superhelical assembly, the protomer models were added to the map through transformation about the helical axis in UCSF ChimeraX (*72*). Figure panels containing structural information were created using ChimeraX version 1.10 and PyMol version 3.0.4.

Genome mining for substrates of LanMs

Substrates of class II lanthipeptide synthetases (LanMs) were collected from UniProt and Refseq databases using several approaches. Members of the two-component *Enterococcus faecalis* cytolysin (EFC) Pfam (PF16934) were collected from UniProt. HMMER was used in combination with the HMM profile of PF16934 to expand this collection of substrates. Additional substrates were mined by mapping accession from the Domain of Unknown Function (DUF4135) Pfam (PF13575) to Refseq accessions followed by exploration of nearby sequences using RODEO (*73*). Biosynthetic gene clusters containing LanJ enzymes (which reduce dehydro amino acids to D-amino acids) were removed (*74*). Finally, previously reported substrates were added (*57, 75*). The leader-core boundary was estimated by searching the leader peptide motif for GG, GA, GS or $(S/T)_2$ sequences and appointing the core region as starting immediately after GG, GA, GS or six residues N-terminal to $(S/T)_2$. The estimated core sequences were analyzed for $S/T(X)_3C$

sequences and grouped according to the number of S/T(X)$_3$C sequences present. The EFI-EST web resource was used to construct a sequence similarity network (*76, 77*). The nonredundant set of core sequences were aligned using Clustal Omega version 1.2.3 (*78*) using five duramycin sequences as an outgroup. A phylogenetic tree was calculated from the multiple sequence alignment using FastTree version 2.1.11 (*78*) and visualized and annotated in iTOL (*79*). A complete list of substrate sequences can be found in Data S1.

<u>The impact of size filtration and dialysis on the hemolytic activity of cytolysin</u>

Defibrinated rabbit erythrocytes (Hemostat Laboratories) were washed with cold PBS as described previously (*57*).

*In vitro dialysis*

CylL$_L$″ and CylL$_S$″ were diluted in PBS (100 μL) such that the total peptide concentration was 4 μM. Washed rabbit erythrocytes were diluted 50-fold in PBS and 1.5 mL of this suspension was transferred to a Slide-A-Lyzer™ MINI microcentrifuge tube (Thermo) and incubated at 37 °C for 30 min. The Slide-A-Lyzer™ MINI Dialysis Device (10 kDa) was placed in contact with the warmed solution of erythrocytes then the cytolysin solution was added to the dialysis device. Care was taken to ensure that the membrane was in contact with both solutions throughout the experiment. A negative control was prepared by adding 100 μL of PBS not containing cytolysin to the dialysis device. A positive control was prepared by adding the same amount of cytolysin present in the sample to the 1.5 mL erythrocyte solution. The apparatuses were incubated at 37 °C for 90 min. Afterwards, the dialysis device was removed, and the intact erythrocytes were pelleted by centrifugation (1000 × g for 10 min). The amount of hemoglobin in the supernatant was evaluated visually. Raw data can be found in Data S1.

*Centrifugal filtration*

Washed rabbit erythrocytes were diluted 20-fold in PBS and the suspension was transferred to a 96-well microtiter plate (90 µL per well). The plate was incubated at 37 °C for at least 30 min. Solutions of cytolysin were prepared in PBS then transferred to Amicon® Ultra Centrifugal Filters equipped with 10 kDa or 100 kDa molecular weight cutoff membranes. The solutions were centrifuged (4000 × g for 30 min) and the filtrate was tested for hemolytic activity by immediately adding to the pre-warmed 96-well plate (60 µL). A positive control was prepared by adding cytolysin directly to the plate without any filtration. A negative control was prepared by adding PBS to the plate. The plate was incubated at 37 °C for 1 h. The plate was centrifuged (1000 × g for 10 min) then 20 µL of the supernatant was transferred to 180 µL of PBS. The absorbance of the diluted supernatant was measured at 415 nm using a plate reader (Biotek Synergy H4). The experiment was done in triplicate ($n$ = 3). Raw data can be found in Data S1.

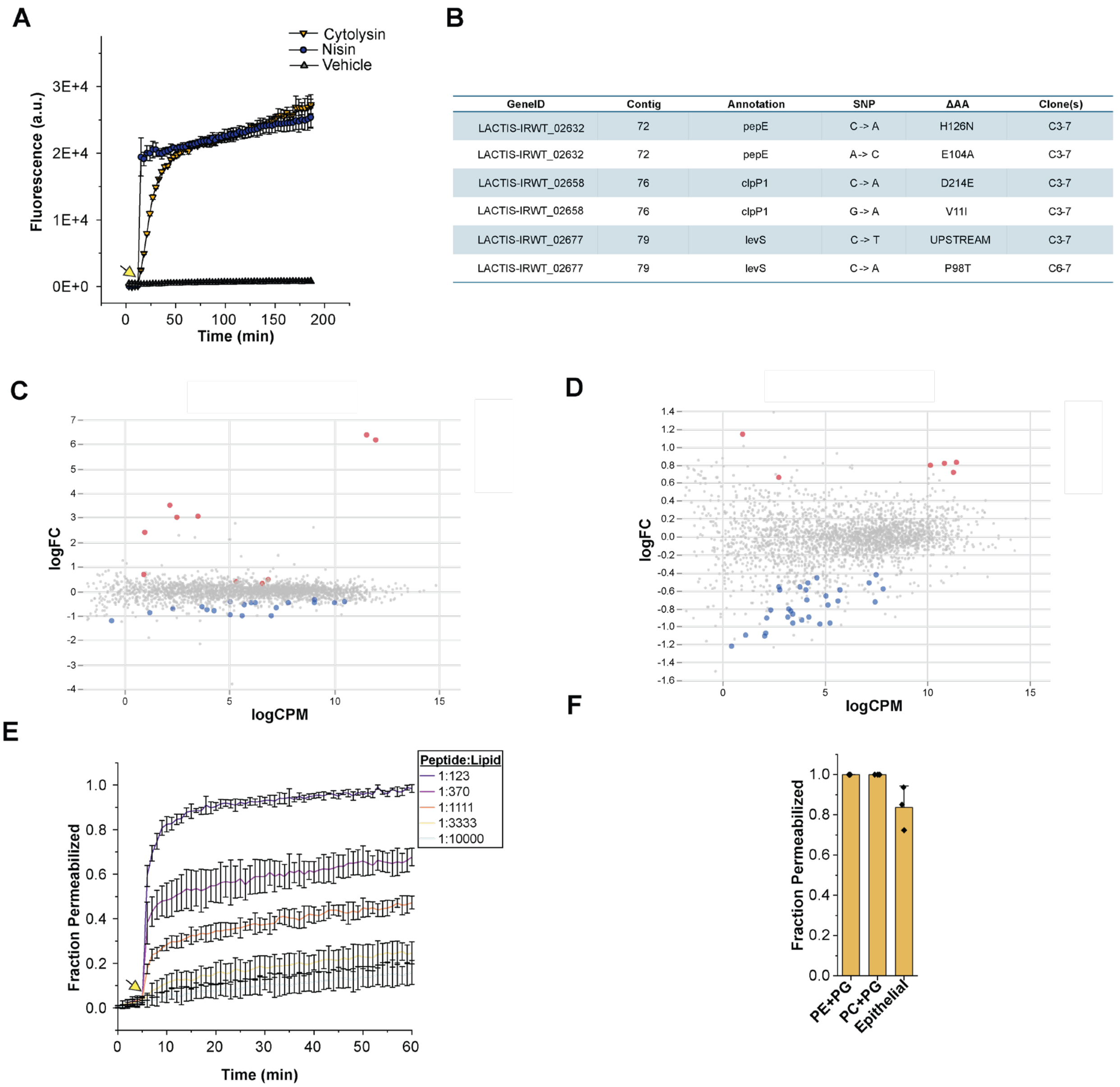


| GeneID | Contig | Annotation | SNP | ΔAA | Clone(s) |
|---|---|---|---|---|---|
| LACTIS-IRWT_02632 | 72 | pepE | C -> A | H126N | C3-7 |
| LACTIS-IRWT_02632 | 72 | pepE | A-> C | E104A | C3-7 |
| LACTIS-IRWT_02658 | 76 | clpP1 | C -> A | D214E | C3-7 |
| LACTIS-IRWT_02658 | 76 | clpP1 | G -> A | V11I | C3-7 |
| LACTIS-IRWT_02677 | 79 | levS | C -> T | UPSTREAM | C3-7 |
| LACTIS-IRWT_02677 | 79 | levS | C -> A | P98T | C6-7 |

**Fig. S1.**

**Cytolysin permeabilizes gram-positive bacteria and resistance towards cytolysin is conferred through a passive resistance mechanism.** A) Membrane permeabilization of *L. lactis subsp. cremoris* to SYTOX green induced by cytolysin or nisin. The yellow arrow indicates when the peptides were added. CylL$_L$″ and CylL$_S$″ were present in 1:1 ratio at a total peptide concentration of 16 nM (2 × MIC). The concentration of nisin was 600 nM (2 × MIC). Data from three separate trials were averaged and presented ($n$ = 3). B) Summary of the mutations corresponding to the emergence of resistance to cytolysin in *L. lactis subsp. cremoris*. *pepE* encodes a serine peptidase specific for dipeptides with an N-terminal Asp (*80*). *clpP1* encodes a caseinolytic protease isoform involved in cell homeostasis (*81*). *levS* encodes for a mucin-binding protein displayed on the surface of *L. lactis* (*82*). C) and D) RNAseq analysis indicates a passive resistance mechanism to cytolysin. Genes with global False Discovery Rate (FDR)-adjusted p-value < 0.2 are colored in red or blue indicating upregulation or down regulation, respectively. A complete list of genes and expression levels can be found in Data S1. C) Comparison of the transcriptional profiles of a parental WT strain and a mutant cytolysin resistant strain of *L. lactis subsp. cremoris* treated with vehicle. The most upregulated genes (top right corner) correspond to two adjacent genes that encode for the ATP-binding protein and transmembrane domain of the ABC transporter (*83*). D) Comparison of the transcriptional profiles of the cytolysin resistant strain treated with vehicle or cytolysin (320 nM). E) Permeabilization of DOPC liposomes induced by $CylL_L''$:$CylL_S''$ (1:1) at varying peptide:lipid (P:L) ratios. The yellow arrow indicates when the peptides were added. Data from three separate trials were averaged ($n$ = 3) then plotted. F) Comparing the membrane permeabilization activity of $CylL_L''$:$CylL_S''$ in a 1:1 ratio over a range of liposome compositions. Model membranes for epithelial cells were composed of PC:PE:phosphatidylserine (PS):sphingomyelin:cholesterol in a 20:20:10:15:35 ratio. The P:L ratio was 1:250. Data from three separate trials ($n$ = 3) is presented. For E) and F) min-max normalization of the emission intensity data yielded fraction permeabilized. Raw data can be found in Data S1.

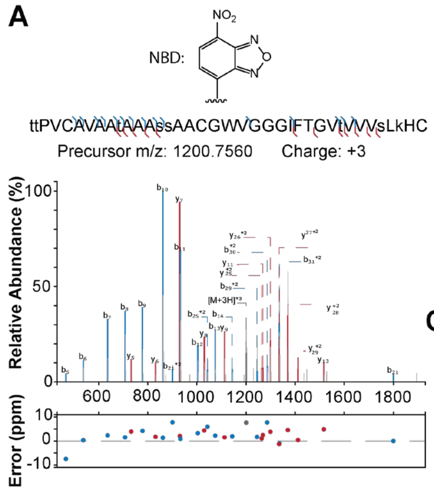


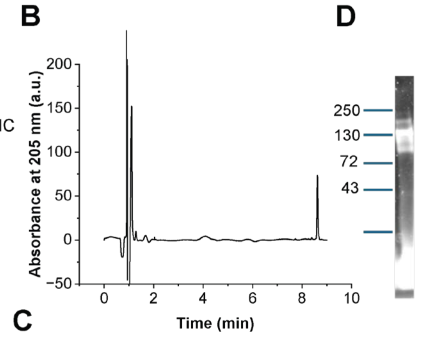


C

| Peptide | MIC (nM) | EC50 of hemolysis (nM) |
|---|---|---|
| $CylL_S$":$CylL_L$" | 4 | 71 ± 3 |
| $CylL_S$":$CylL_L$"-NBD | 64 | 196 ± 12 |

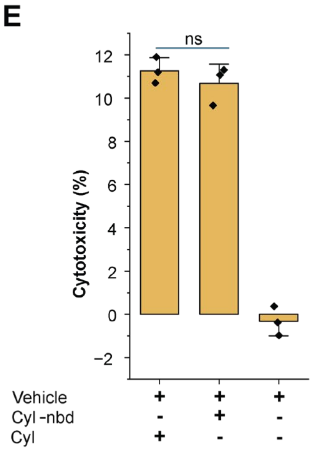


**Fig. S2.**

**Characterization of $CylL_L$″-NBD.** A) High-resolution MS-MS analysis of a purified sample of $CylL_L$″-NBD. Lowercase letters indicate modifications. Ser/Thr residues were dehydrated according to the structure of cytolysin (*84*). The fragmentation data verify that Lys36 was modified by the mass of the NBD group. The annotations and plot were created using the Interactive Peptide Spectral Annotator (*85*). B) UV-vis trace collected during LC-MS analysis of a sample of $CylL_L$″-NBD. A wavelength of 205 nm was used. The trace was baseline corrected using Origin Pro. The retention time of $CylL_L$″-NBD is 10.6 min. C) The antibiotic and hemolytic activities of cytolysin ($CylL_S$″:$CylL_L$″ in a 1:1 ratio) and fluorescently labelled cytolysin ($CylL_S$″:$CylL_L$″-NBD in a 1:1 ratio). D) Negative image of the fluorescence signal from SDS-PAGE analysis of a stoichiometric mixture of $CylL_L$″-NBD and $CylL_S$″. The gel was analyzed by an iBright™ CL750 Imaging System in fluorescence mode. $CylL_L$″-NBD and $CylL_S$″ form a similar higher molecular weight band around 130 kDa as the WT peptides. An uncropped image was added to Data S2. E) LDH release from HepG2 cells 1 h after treatment with stoichiometric mixtures of $CylL_L$″-NBD and $CylL_S$″ (Cyl-nbd) or $CylL_L$″ and $CylL_S$″ (Cyl). A total peptide concentration of 10 µM was used. This experiment was done in triplicate ($n$ = 3). The bar represents the mean, and the error bar represents the standard of deviation. Samples were compared with a two-sample $t$ test. ns indicates a $P$ value > 0.05. Raw data is provided in Data S1 and Data S2.

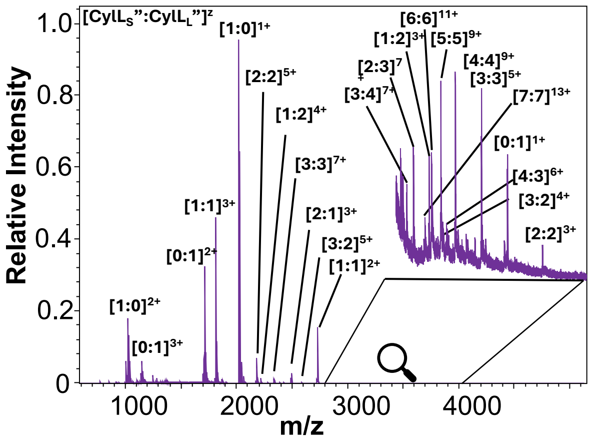

**Fig. S3.**

**Detection of cytolysin heterooligomers by high-resolution mass spectrometry.** High-resolution MS spectrum of a stoichiometric mixture of $CylL_L''$ and $CylL_S''$ diluted in 1:1 MeOH:$H_2O$. The MS data highlights the first steps of the oligomerization process between $CylL_L''$ and $CylL_S''$ in the gas-phase, showing the formation $CylL_L''$:$CylL_S''$ complexes up to 7:7 stoichiometry. The *m/z* peaks were labelled with the combination of subunits that correspond to the observed masses. Raw data can be found in Data S3.

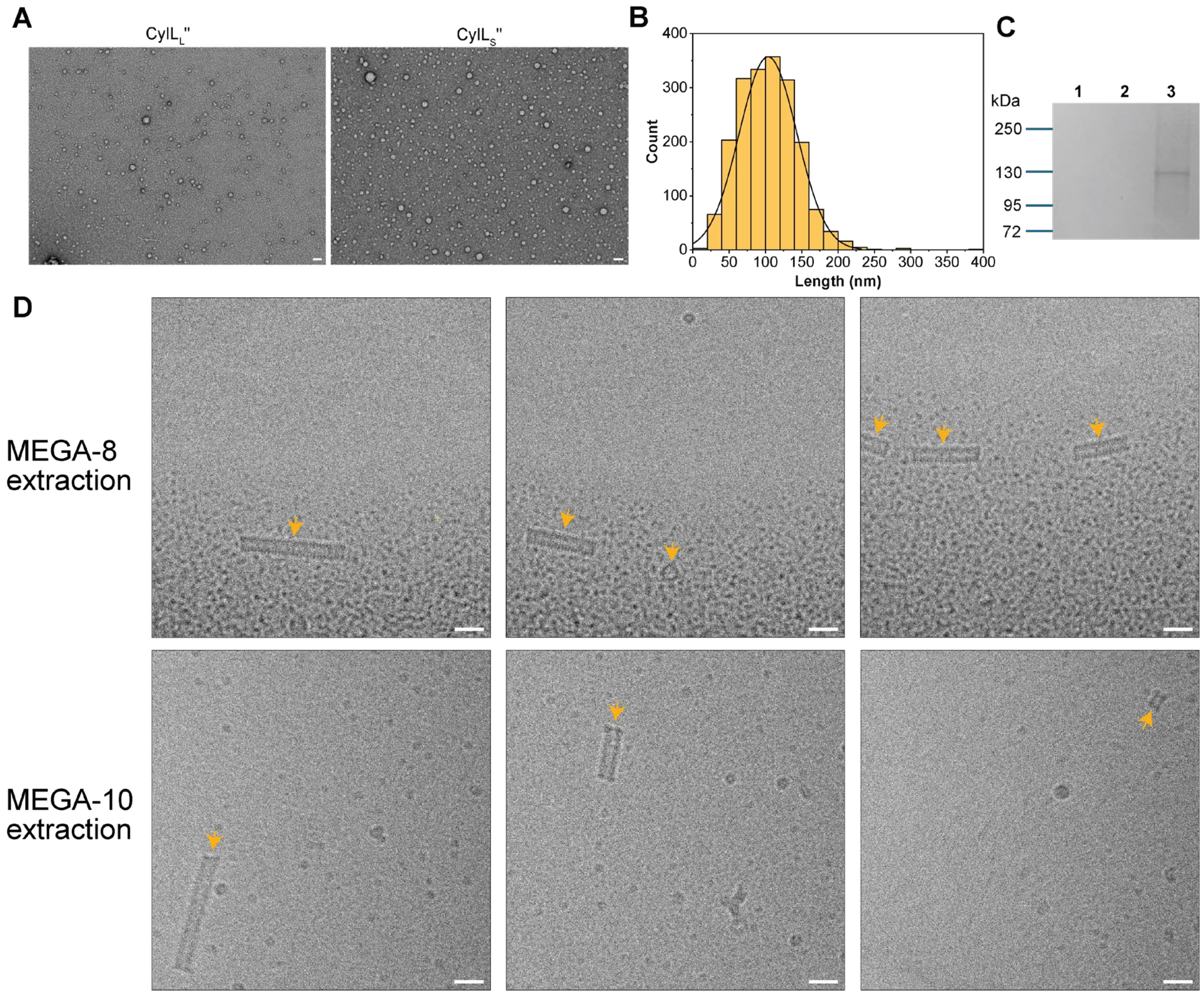


**Fig. S4.**

**The properties of cytolysin nanotubes.** A) Negatively stained samples of CylL$_S$″ and CylL$_L$″ analyzed by TEM. Samples were diluted to a concentration of 50 μM in 50 mM HEPES-KOH (pH 7.5) and 150 mM NaCl and incubated overnight at room temperature before analysis. Scale bars represent 50 nm. B) Length distribution of cytolysin nanotube assemblies with lengths greater than 10 nm. ImageJ was used to measure the lengths of 1,927 pipes from 24 micrographs (*86*). The black line represents a fit to the normal distribution, which was calculated using Origin Pro. C) Samples of CylL$_L$″ (lane 1), CylL$_S$″ (lane 2) and a 1:1 mixture of CylL$_S$″:CylL$_L$″ (lane 3) analyzed by SDS-PAGE. D) Cryo-EM micrographs of cytolysin assemblies extracted from DOPC liposomes with non-ionic detergents. The detergents used are shown to the left of micrographs; extractions were performed with 79 mM MEGA-8 and 7 mM MEGA-10, in a buffer containing 50 mM HEPES-KOH (pH 7.5) and 150 mM NaCl. Arrows indicate putative nanotubes and particles in different orientations. Scale bars represent 20 nm. Original micrographs can be found in Data S2.

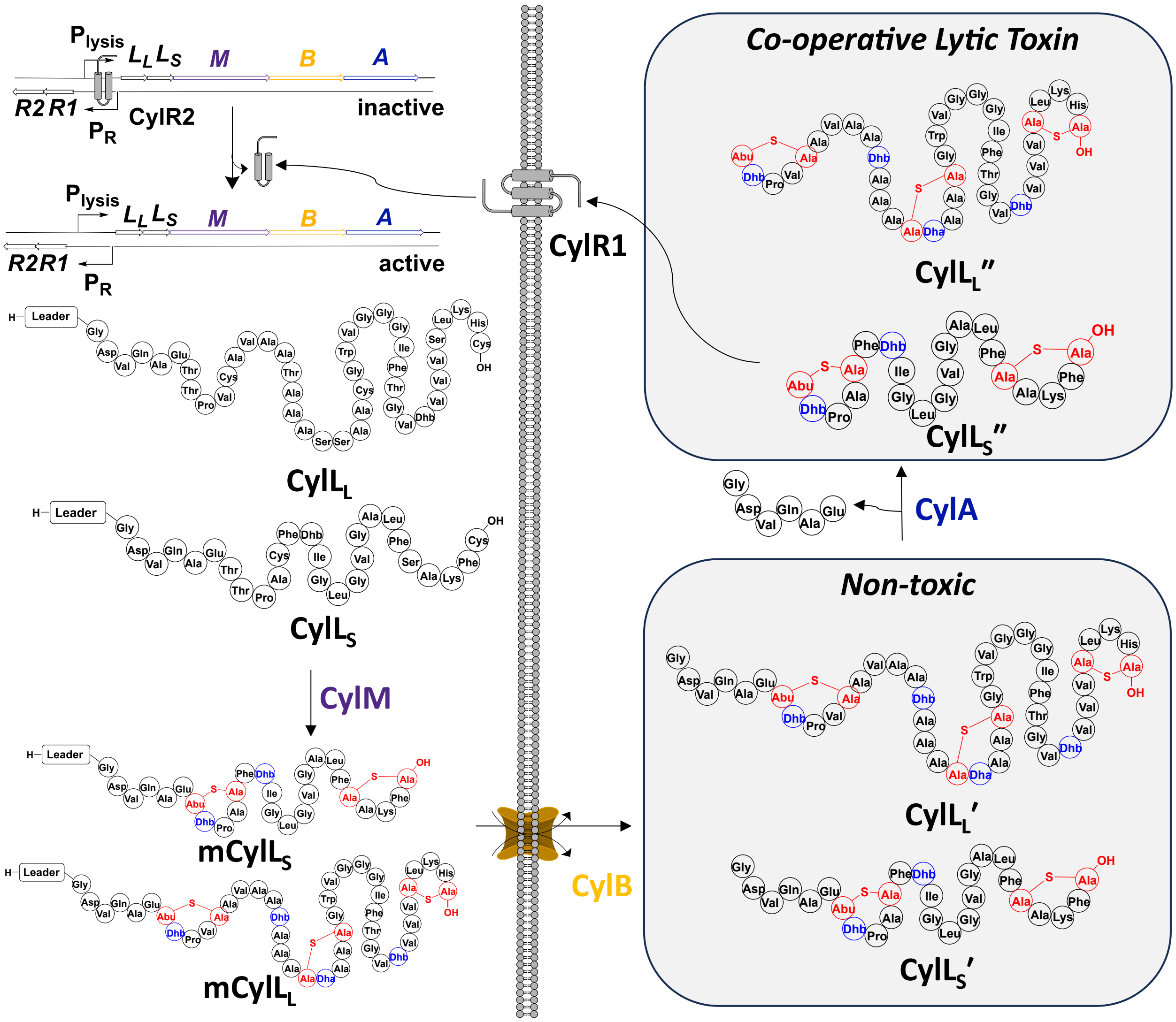


**Fig. S5.**

**The biosynthesis of cytolysin in *Enterococcus faecalis*.** The substrates CylL$_L$ and CylL$_S$ are ribosomally synthesized then modified by the bifunctional enzyme CylM (*87*). The dehydration domain of CylM converts Ser/Thr to dehydroalanine (Dha) and dehydrobutyrine (Dhb), respectively. The cyclization domain catalyzes Michael-type addition of the Cys side chain thiol to Dha and Dhb, forming lanthionine and methyllanthionine bridges, respectively. The fully modified full-length precursors, mCylL$_S$ and mCylL$_L$, are the substrates of the transporter-protease CylB which removes most of the N-terminal leader peptide and generates the inactive precursors CylL$_L$′ and CylL$_S$′. An extracellular protease called CylA converts CylL$_L$′ and CylL$_S$′ to CylL$_L$″ and CylL$_S$″ through removal of an N-terminal hexapeptide sequence (GDVQAE). CylL$_S$″ induces cytolysin expression through interaction with the response regulator CylR1 and CylR2 (*88*).

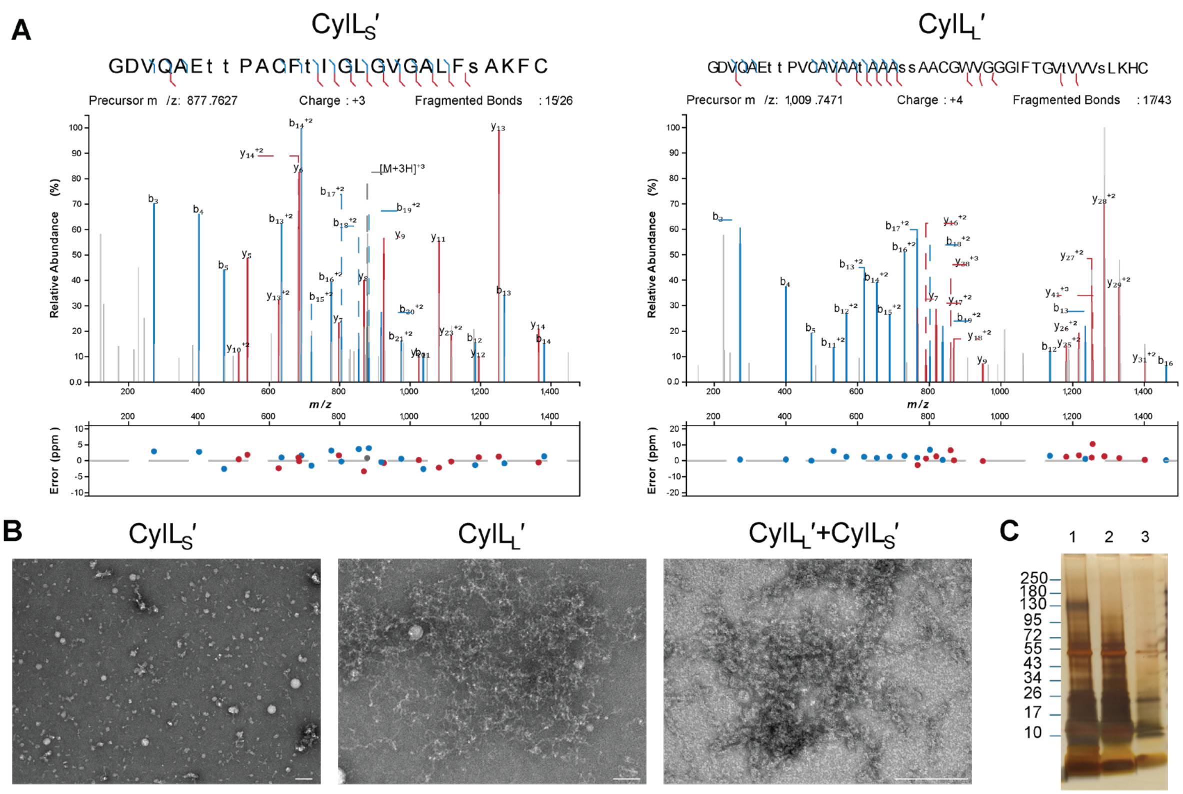


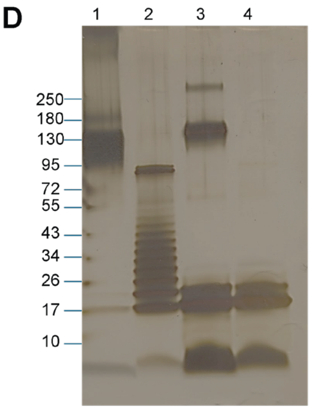


E

| Peptide | MIC (nM) | EC50 of hemolysis (nM) |
|---|---|---|
| CylL$_S$″:CylL$_L$″ | 4 | 71 ± 3 |
| CylL$_S$″:CylL$_L$″ C5A | 128 | 222 ± 31 |
| CylL$_S$″ C5A:CylL$_L$″ | 4 | 141 ± 15 |
| CylL$_S$″ C5A:CylL$_L$″ C5A | >256 | >4000 |
| CylL$_S$′:CylL$_L$′ | >256 | >4000 |

**Fig. S6.**

**Characterization and bioactivity of WT cytolysin and analogs.** A) MS-MS analysis of the cytolysin precursors CylL$_S$′ and CylL$_L$′. B) Negatively stained samples of CylL$_S$′, CylL$_L$′ and a stoichiometric mixture of the two subunits analyzed by TEM. Samples were diluted to a concentration of 50 µM in 50 mM HEPES-KOH (pH 7.5) and 150 mM NaCl and incubated overnight at room temperature before staining and analysis. White scale bars represent 50 nm. C) SDS-PAGE analysis of pre-cytolysin peptides individually or as a stoichiometric mixture. Lane 1: CylL$_L$′ and CylL$_S$′. Lane 2: CylL$_L$′. Lane 3: CylL$_S$′. D) SDS-PAGE analysis of stoichiometric mixtures of WT cytolysin and CylL$_L$″ C5A or CylL$_S$″ C5A. Lane 1: CylL$_L$″ and CylL$_S$″. Lane 2: CylL$_S$″ and CylL$_L$″ C5A. Lane 3: CylL$_S$″ C5A and CylL$_L$″. Lane 4: CylL$_L$″ C5A and CylL$_S$″ C5A. E) The antibiotic and hemolytic activities of cytolysin and analogs thereof. See Data S2 and Data S1 for original images and raw data.

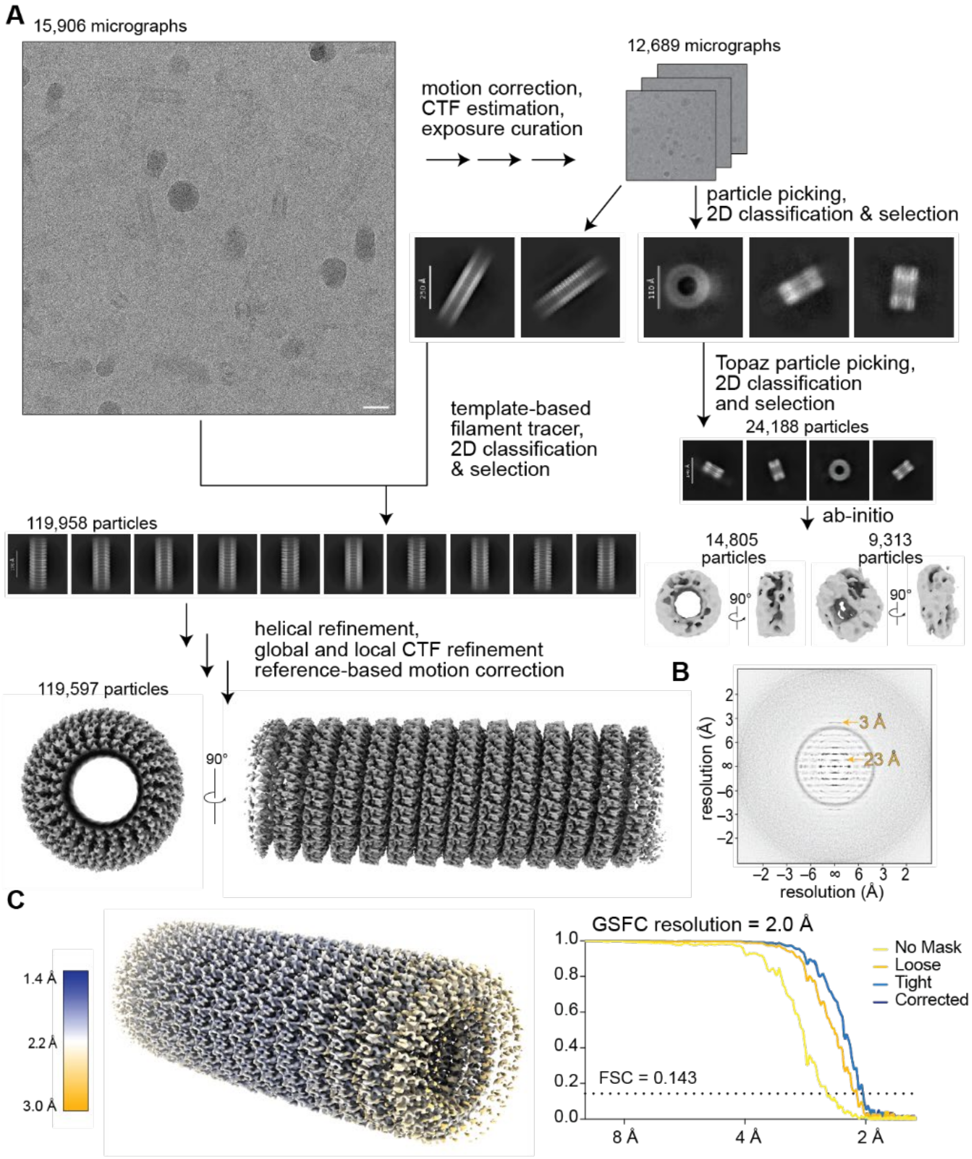


**Fig. S7.**

**Cryo-EM data processing of cytolysin oligomers.** A) Workflow from representative cryo-EM micrograph of cytolysin oligomers, 2D classification, and helical refinement or 3D reconstruction. The helical reconstruction was improved with global CTF refinement, local CTF refinement, and reference-based motion correction. The small pore-like assemblies were not processed beyond ab initio models due to low particle count. All processing was performed with cryoSPARC (*66*). Scale bar = 50 nm. B) The sum in-plane rotated power spectra of cytolysin tube segments from the entire dataset. The results give rise to a layer-line pattern typical for helical assemblies where the line visible at 23 Å is consistent with the 23 Å pitch and lines are observed past 3 Å, consistent with the high-resolution data. C) Left, local resolution estimate of final map 2.0 Å map colored according to gradient bar on the left. Right, Fourier shell correlation (FSC) curves versus resolution of the cytolysin helical oligomer. The resolution was estimated at an FSC of 0.143.

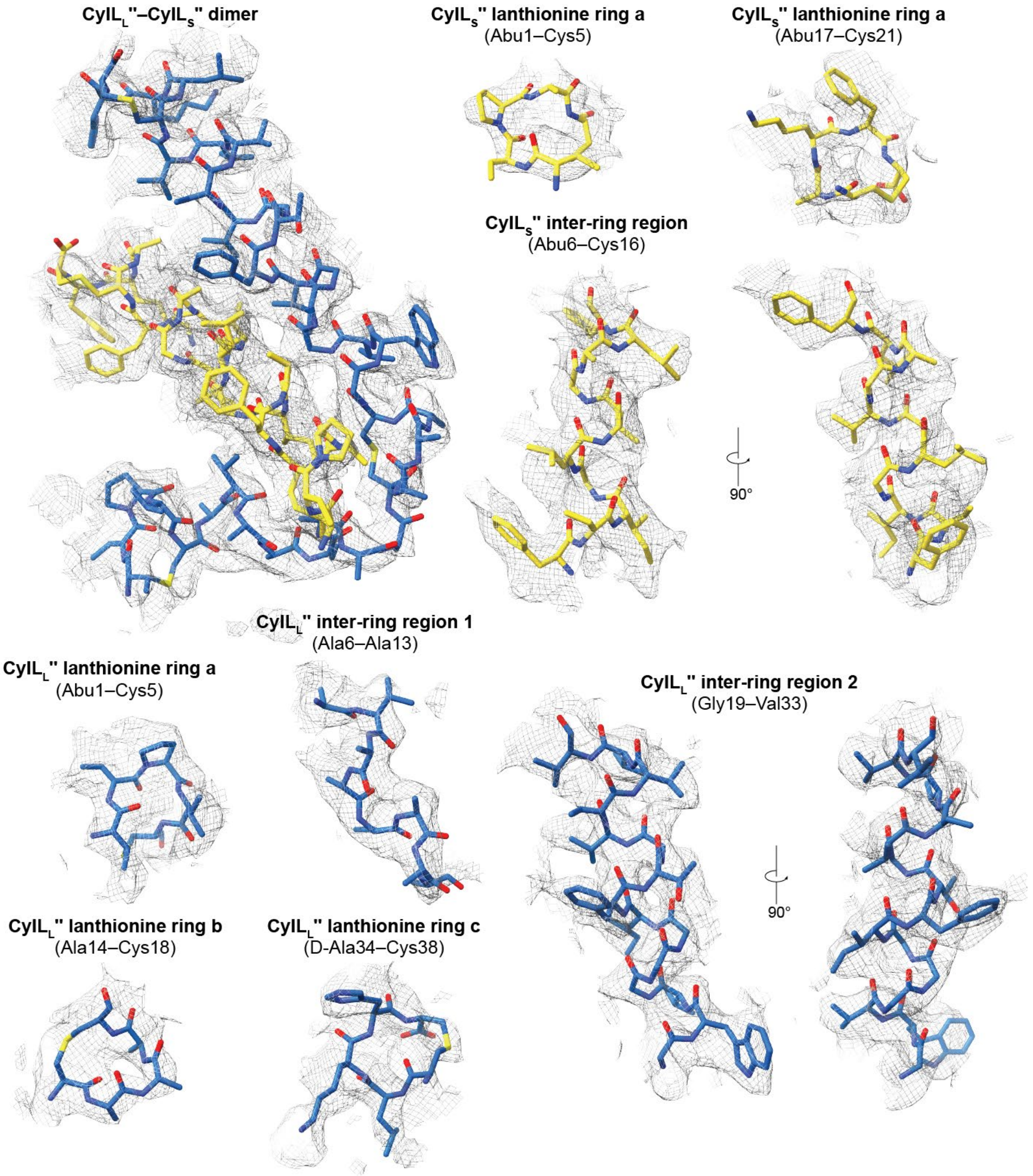


**Fig. S8.**

**Cryo-EM model to map fitting of cytolysin.** Regions of model to map fit quality for the CylL$_S$″–CylL$_L$″ dimer from the nanotube structure. The map surface has been contoured to 4.5σ.

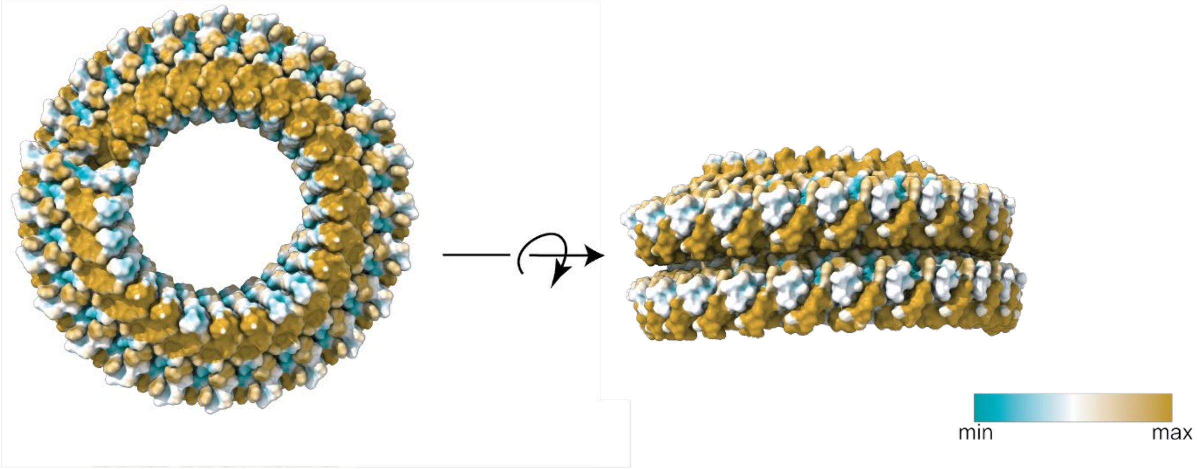


**Fig. S9.**

**Surface of the assembly generated by 36 dimers (1.5 helical turns) of CylL$_S$″ and CylL$_L$″ colored according to lipophilicity**. Molecular lipophilicity was calculated and plotted using USCF Chimera X 1.10.1 (*89*). Cyan represents the most hydrophilic atoms, and golden rod represents the most hydrophobic residues.

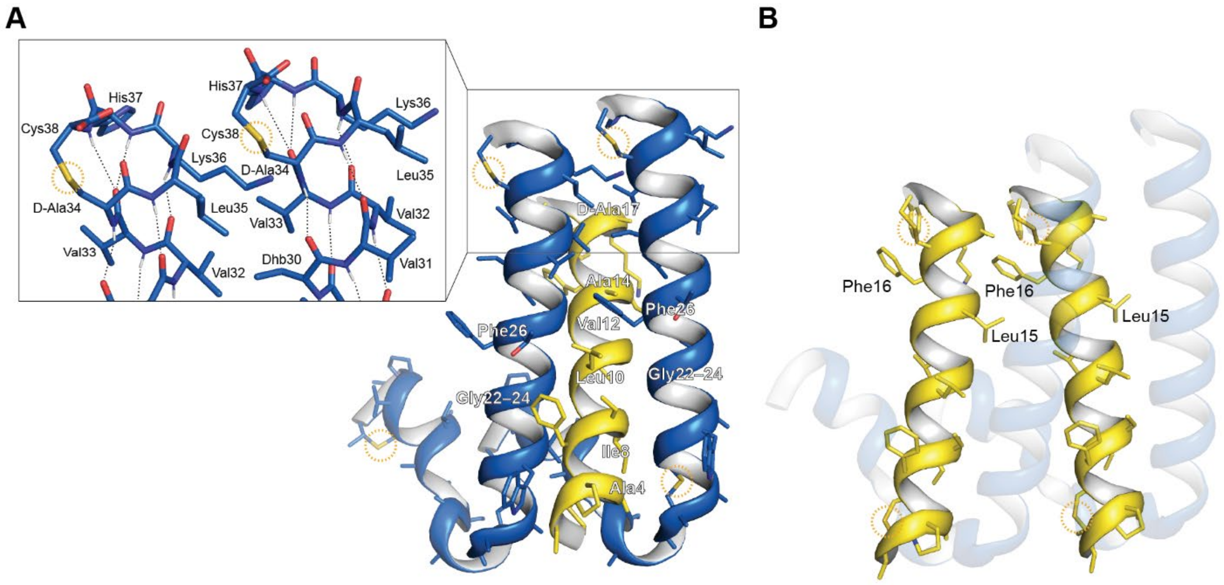


**Fig. S10.**

**Additional CylL$_L$″-CylL$_L$″, CylL$_S$″-CylL$_L$″, and CylL$_S$″-CylL$_S$″ interactions.** A) Key residues involved in the interaction between CylL$_S$″ and adjacent units of CylL$_L$″. The inset highlights the interdigital pattern of hydrophobic residues near the C-termini of adjacent CylL$_L$″ subunits. B) Key residues involved in the interaction between CylL$_S$″ monomers in adjacent dimers within a helical turn.

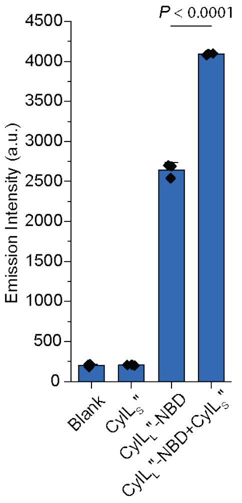


**Fig. S11.**

**The fluorescence emission intensity of CylL$_L$″-NBD increases upon addition of CylL$_S$″.** NBD is an environmentally sensitive fluorophore that displays an increase in quantum yield when moved from a hydrophilic to a hydrophobic environment (*90*). Peptide or vehicle was diluted into PBS (pH = 7.4) at room temperature, incubated at room temperature for 10 min, then the fluorescence emission intensity was determined using a plate reader (Biotek Synergy H4) with an excitation wavelength equal to 460 nm and an emission wavelength equal to 540 nm. The total peptide concentration was 500 nM. This experiment was performed in triplicate ($n$ = 3). A two-sample $t$ test was used for hypothesis testing.

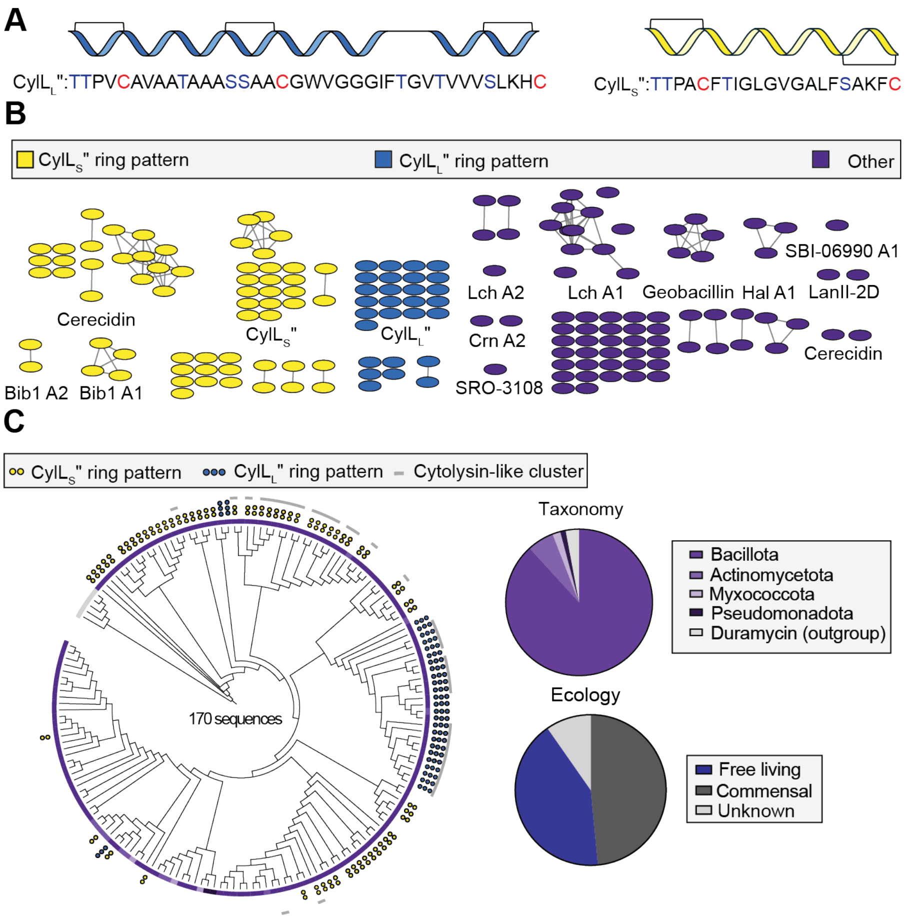


**Fig. S12.**

**Genome mining reveals conservation of helix-templating thioethers at the N- and C-termini of lanthipeptide substrates.** A) Diagram depicting the ring pattern of CylL$_L$″ and CylL$_S$″. S/T(X)$_3$C motifs at the N- and C-termini template helical conformations and this pattern was used to form heuristics to guide genome mining towards α-helical lanthipeptides. B) Sequence similarity network constructed from the core sequences of class II lanthipeptide substrates. Sequences that were identical were consolidated prior to SSN construction. Nodes are colored according to the ring pattern predicted from their sequences. Clusters containing two or more nodes were labelled with the highest similarity hit from a BLAST of the MIBiG database with a representative sequence (*91*). If this did not yield a hit, the cluster was left unlabelled. C) A tree constructed from core sequences colored according to phylum. A grey bar at the outer periphery indicates if the sequences were part of a BGC that encoded CylL$_S$″-like and CylL$_L$″-like components. The colored dots above each node indicate the ring pattern of the constituent sequences. Three blue dots indicate a CylL$_L$″-like pattern. Two yellow dots indicate a CylL$_S$″-like pattern. Sequences were classified based on the taxonomy and ecology of their producing organisms. The pie charts illustrate the taxonomic and ecological distributions. A complete list of sequences can be found in Data S1.

**A**

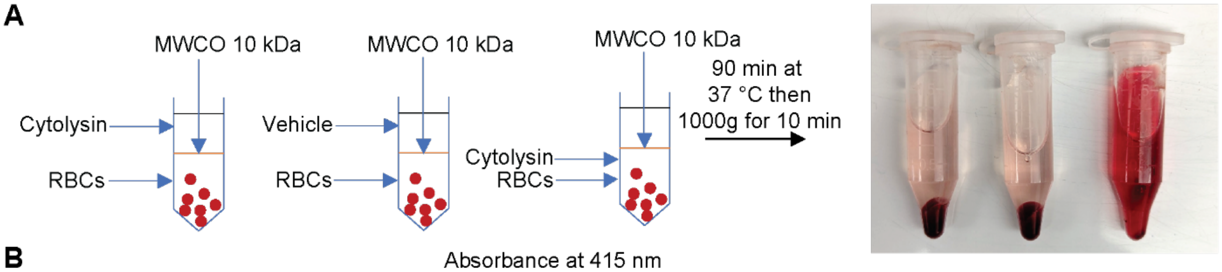


**B**

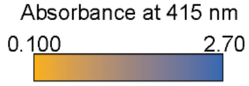


| | 10 kDa MWCO | | | 100 kDa MWCO | | | No filtration | | |
|---|---|---|---|---|---|---|---|---|---|
| Concentration of cytolysin (μM) | Trial 1 | Trial 2 | Trial 3 | Trial 1 | Trial 2 | Trial 3 | Trial 1 | Trial 2 | Trial 3 |
| 2.0 | 0.153 | 0.15 | 0.147 | 0.145 | 0.137 | 0.15 | 2.272 | 2.012 | 1.775 |
| 1.0 | 0.148 | 0.143 | 0.144 | 0.149 | 0.137 | 0.134 | 2.522 | 2.257 | 2.465 |
| 0.50 | 0.148 | 0.143 | 0.15 | 0.148 | 0.131 | 0.135 | 2.662 | 2.385 | 2.204 |
| 0.25 | 0.149 | 0.142 | 0.143 | 0.14 | 0.131 | 0.13 | 2.510 | 2.511 | 2.603 |
| 0.13 | 0.14 | 0.138 | 0.137 | 0.128 | 0.134 | 0.13 | 2.199 | 2.009 | 2.244 |
| 0.063 | 0.145 | 0.139 | 0.137 | 0.136 | 0.128 | 0.128 | 0.795 | 1.158 | 1.221 |
| 0.031 | 0.155 | 0.144 | 0.146 | 0.126 | 0.126 | 0.127 | 0.177 | 0.178 | 0.180 |
| 0.016 | 0.146 | 0.141 | 0.162 | 0.128 | 0.128 | 0.13 | 0.117 | 0.109 | 0.120 |

**Fig S13.**

**Samples of cytolysin are inactivated by dialysis or centrifugal filtration.** A) To the left is a visual description of the dialysis experiments. Cytolysin (4 μM) or vehicle in 100 μL of phosphate buffered saline (PBS) was added to a dialysis chamber in contact with 1.5 mL of 5% rabbit erythrocytes (RBCs) suspended in PBS. The molecular weight cutoff (MWCO) of this membrane was 10 kDa. A positive control sample was created by adding the same amount of cytolysin to the solution of RBCs. All samples were incubated for 90 min at 37 °C. Intact RBCs were pelleted by centrifugation after removing the dialysis chamber. The picture to the right was taken immediately after centrifugation. B) Samples of cytolysin at various concentrations were filtered using centrifugal filters (Amicon®) with MWCOs of 10 kDa or 100 kDa. The filtrate was tested for hemolytic activity next to an unfiltered control (these wells are under the heading 'No filtration'). Values within grid squares represent the absorbance at 415 nm of the supernatant of the corresponding well which is proportional to the amount of hemoglobin released during RBC lysis. Wells are colored with a linear gradient from orange (no hemolysis) to blue (complete hemolysis).

**Table. S1. Cryogenic-electron microscopy data summary table.**

**Cryo-EM data collection, refinement and validation statistics**

| | *EfCytolysin nanotube* (EMD-75911) (PDB 11PC) |
|---|---|
| **Data collection and processing** | |
| Magnification | 165,000 |
| Voltage (kV) | 300 |
| Electron exposure (e–/Å$^2$) | 54.8 |
| Exposure time | 2.58 |
| Number of fractions | 88 |
| Pixel size (Å) | 0.74 |
| Detector | Falcon4 + selectris |
| Dose rate detector (e$^-$/pix/s) | 1.64 |
| Number of movies | 15,906 |
| Defocus range (μm) | −0.7 to −1.8 |
| Initial particle images (no.) | 356,404 |
| Final particle images (no.) | 119,597 |
| Map resolution (Å) | 2.0 |
| FSC threshold | 0.143 |
| Map resolution range (Å) | 1.44–2.07 |
| Helical rise (Å) | 0.97 |
| Helical twist | –15.05° |
| Final number of asym. Units | 5,142,671 |
| **Refinement** | |
| Initial model used | ModelAngelo of unmodified peptides |
| Sequences | fasta files from PDB 9ve9, 9vgt |
| Model resolution (Å) | 2.1 |
| FSC threshold | 0.143 |
| Model resolution range (Å) | 2.0–3.7 |
| Map sharpening *B* factor (Å$^2$) | −30.2 |
| Model composition | |
| Non-hydrogen atoms | 395 |
| Protein residues | 55 |
| *B* factors (Å$^2$) | |
| Protein | 36.01–78.19 |
| R.m.s. deviations | |
| Bond lengths (Å) | 0.004 |
| Bond angles (°) | 1.492 |
| Validation | |
| MolProbity score | 2.28 |
| Clashscore | 14.77 |
| Poor rotamers (%) | 3.70 |
| Ramachandran plot | |
| Favored (%) | 96.97 |
| Allowed (%) | 3.03 |
| Disallowed (%) | 0.00 |

**Data S1. (separate file)**
This file contains the tabulated data behind the figures.

**Data S2. (separate file)**
This file contains unedited gel images and transmission electron micrographs.

72. E. C. Meng *et al.*, UCSF ChimeraX: Tools for structure building and analysis. *Protein Sci.* **32**, e4792 (2023).
73. J. I. Tietz *et al.*, A new genome-mining tool redefines the lasso peptide biosynthetic landscape. *Nat. Chem. Biol.* **13**, 470-478 (2017).
74. X. Yang, W. A. van der Donk, Post-translational introduction of D-alanine into ribosomally synthesized peptides by the dehydroalanine reductase NpnJ. *J. Am. Chem. Soc.* **137**, 12426-12429 (2015).
75. M. C. Walker *et al.*, Precursor peptide-targeted mining of more than one hundred thousand genomes expands the lanthipeptide natural product family. *BMC Genom.* **21**, 387 (2020).
76. R. Zallot, N. Oberg, J. A. Gerlt, The EFI web resource for genomic enzymology tools: Leveraging protein, genome, and metagenome databases to discover novel enzymes and metabolic pathways. *Biochemistry* **58**, 4169-4182 (2019).
77. N. Oberg, R. Zallot, J. A. Gerlt, EFI-EST, EFI-GNT, and EFI-CGFP: Enzyme Function Initiative (EFI) web resource for genomic enzymology tools. *J. Mol. Biol.* **435**, 168018 (2023).
78. F. Sievers *et al.*, Fast, scalable generation of high-quality protein multiple sequence alignments using Clustal Omega. *Mol. Syst. Biol.* **7**, 539 (2011).
79. I. Letunic, P. Bork, Interactive Tree Of Life (iTOL) v5: an online tool for phylogenetic tree display and annotation. *Nucleic Acids Res.*, W293-W296 (2021).
80. P. Yadav *et al.*, Structure of Asp-bound peptidase E from Salmonella enterica: Active site at dimer interface illuminates Asp recognition. *FEBS Letters* **592**, 3346-3354 (2018).
81. E. Zeiler *et al.*, Structural and functional insights into caseinolytic proteases reveal an unprecedented regulation principle of their catalytic triad. *Proc. Natl. Acad. Sci. U. S. A.* **110**, 11302-11307 (2013).
82. W. Tsuchiya *et al.*, Cell-surface protein YwfG of Lactococcus lactis binds to α-1,2-linked mannose. *PLOS ONE* **18**, e0273955 (2023).
83. M. Ogura, K. Tsukahara, K. Hayashi, T. Tanaka, The Bacillus subtilis NatK–NatR two-component system regulates expression of the natAB operon encoding an ABC transporter for sodium ion extrusion. *Microbiology* **153**, 667-675 (2007).
84. W. Tang, W. A. van der Donk, The sequence of the enterococcal cytolysin imparts unusual lanthionine stereochemistry. *Nat. Chem. Biol.* **9**, 157-159 (2013).
85. D. R. Brademan, N. M. Riley, N. W. Kwiecien, J. J. Coon, Interactive peptide spectral annotator: A versatile web-based tool for proteomic applications. *Mol. Cell. Proteom.* **18**, S193-S201 (2019).
86. J. Schindelin, C. T. Rueden, M. C. Hiner, K. W. Eliceiri, The ImageJ ecosystem: An open platform for biomedical image analysis. *Mol. Reprod. Dev.* **82**, 518-529 (2015).
87. D. Van Tyne, M. J. Martin, M. S. Gilmore, Structure, function, and biology of the *Enterococcus faecalis* cytolysin. *Toxins (Basel)* **5**, 895-911 (2013).
88. P. S. Coburn, C. M. Pillar, B. D. Jett, W. Haas, M. S. Gilmore, *Enterococcus faecalis* senses target cells and in response expresses cytolysin. *Science* **306**, 2270-2272 (2004).
89. E. F. Pettersen *et al.*, UCSF ChimeraX: Structure visualization for researchers, educators, and developers. *Protein Sci.* **30**, 70-82 (2021).
90. S. Fery-Forgues, J.-P. Fayet, A. Lopez, Drastic changes in the fluorescence properties of NBD probes with the polarity of the medium: involvement of a TICT state? *J. Photochem. Photobiol. A: Chem.* **70**, 229-243 (1993).

91. S. A. Kautsar *et al.*, MIBiG 2.0: a repository for biosynthetic gene clusters of known function. *Nucleic Acids Res.* **48**, D454-D458 (2020).
